# Validation of lipid-related therapeutic targets for coronary heart disease prevention using human genetics

**DOI:** 10.1101/2020.11.11.377747

**Authors:** María Gordillo-Marañón, Magdalena Zwierzyna, Pimphen Charoen, Fotios Drenos, Sandesh Chopade, Tina Shah, Jorgen Engmann, Juan-Pablo Casas, Nishi Chaturvedi, Olia Papacosta, Goya Wannamethee, Andrew Wong, Reecha Sofat, Mika Kivimaki, Jackie F Price, Alun D Hughes, Tom R Gaunt, Deborah A Lawlor, Anna Gaulton, Aroon D Hingorani, Amand F Schmidt, Chris Finan

## Abstract

Drug target Mendelian randomization (MR) studies use DNA sequence variants in or near a gene encoding a drug target, that alter its expression or function, as a tool to anticipate the effect of drug action on the same target. Here, we applied MR to prioritize drug targets for their causal relevance for coronary heart disease (CHD). The targets were further prioritized using genetic co-localization, protein expression profiles from the Human Protein Atlas and, for targets with a licensed drug or an agent in clinical development, by sourcing data from the British National Formulary and clinicaltrials.gov. Out of the 341 drug targets identified through their association with circulating blood lipids (HDL-C, LDL-C and triglycerides), we were able to robustly prioritize 30 targets that might elicit beneficial treatment effects in the prevention or treatment of CHD. The prioritized list included NPC1L1 and PCSK9, the targets of licensed drugs whose efficacy has been already proven in clinical trials. To conclude, we discuss how this approach can be generalized to other targets, disease biomarkers and clinical end-points to help prioritize and validate targets during the drug development process.

**One Sentence Summary:** We provide genetic support for lipid-modifying drug targets for coronary heart disease prevention using drug target Mendelian randomization and further prioritization based on clinical and biological evidence.

## Introduction

Genome-wide association studies (GWAS) in patients and populations test relationships between natural sequence variation (genotype) and disease risk factors, biomarkers and clinical end-points using population-based cohort and case-control studies.

Mendelian randomization (MR) analysis uses genetic variants (mostly identified from GWAS) as instrumental variables to identify which disease biomarkers (e.g. blood lipids such as low- and high-density lipoprotein cholesterol and triglycerides) may be causally related to disease end-points (e.g. coronary heart disease; CHD) *(1, 2)*. The established approach utilizes multiple independent variants associated with the biomarker of interest, selected from throughout the genome. These genetic instruments are used to derive an estimate of the effect of a change in the level of the biomarker on disease risk. We refer to this approach as MR analysis for causal biomarkers or ‘biomarker MR’. For example, biomarker MR studies have validated the causal role of elevated low-density lipoprotein cholesterol (LDL-C) on coronary heart disease risk, supporting the findings from randomized controlled trials of different LDL-C lowering drug classes *(3–8)*. However, such studies have been equivocal on the role of high-density lipoprotein cholesterol (HDL-C) and triglycerides (TG) in CHD *(3, 4)*. Clinical trials of these lipid fractions have also been seemingly contradictory. For example, using niacin to raise HDL-C did not reduce CHD risk *(9)*, but raising HDL-C by inhibiting cholesteryl ester transfer protein (CETP) with anacetrapib *was* effective in preventing CHD events *(10)*.

Genetic effects (like drug action) are mediated through proteins (according to Crick’s Central Dogma), and variation in the genome is inherited at random from parents to offspring (according to Mendel’s Laws), much like treatment allocation in a clinical trial *(11)*. We and others have shown that variants in a gene encoding a specific drug target, that alter its expression or function, can be used as a tool to anticipate the effect of drug action on the same target *(12)*. We have referred to this application of Mendelian randomization as ‘drug target MR’ *(12)*. In contrast to a biomarker MR, where the variants comprising the genetic instrument are selected from across the genome, in a drug target MR analysis, variants are selected from the gene of interest or neighboring genomic region because these variants are most likely to associate with the expression or function of the encoded protein (acting in *cis*). The estimate from a drug target MR helps infer whether, and in what direction, a drug that acts on the encoded protein, whether an antagonist, agonist, activator or inhibitor, will alter disease risk. The conceptual and analytical differences between drug target and biomarker MR (Table 1) are important because a narrow interpretation of a biomarker MR analysis of HDL-C and CHD might suggest that CETP inhibitors, which raise HDL-C, should not be regarded as a valid therapeutic intervention to reduce CHD risk. Yet, the causal effect on CHD per SD increase in HDL-C from the drug target MR using variants in *CETP* (0.87; 95%CI: 0.84, 0.90), and the odds ratio for CHD from allocation to the CETP-inhibitor anacetrapib in a placebo-controlled trial (0.93; 95%CI: 0.86, 0.99) are consistent with the view that *targeting CETP* is an effective therapeutic approach to prevent CHD (Fig. 1) *(10, 12)*. This implies that, regardless of the findings of a biomarker MR analysis, other similarly effective but yet unexploited drug targets might exist for the prevention or treatment of CHD, and be identified through their association with blood lipids.

**Table 1.** Main conceptual differences between *biomarker* and *drug target* MR approaches.

|  | **Biomarker MR** | **Drug target MR** |
| --- | --- | --- |
| **Aim** | Causal effect of a biomarker | Causal relevance of a drug target |
| **SNP selection** | Genome-wide | Locus specific |
| **Ideal exposure** | Clinically relevant quantitative trait | mRNA or protein expression of the encoded gene |
| **MR methods** | Any described MR method | Methods accounting for residual genetic correlation to maximize power |

**Fig. 1.**
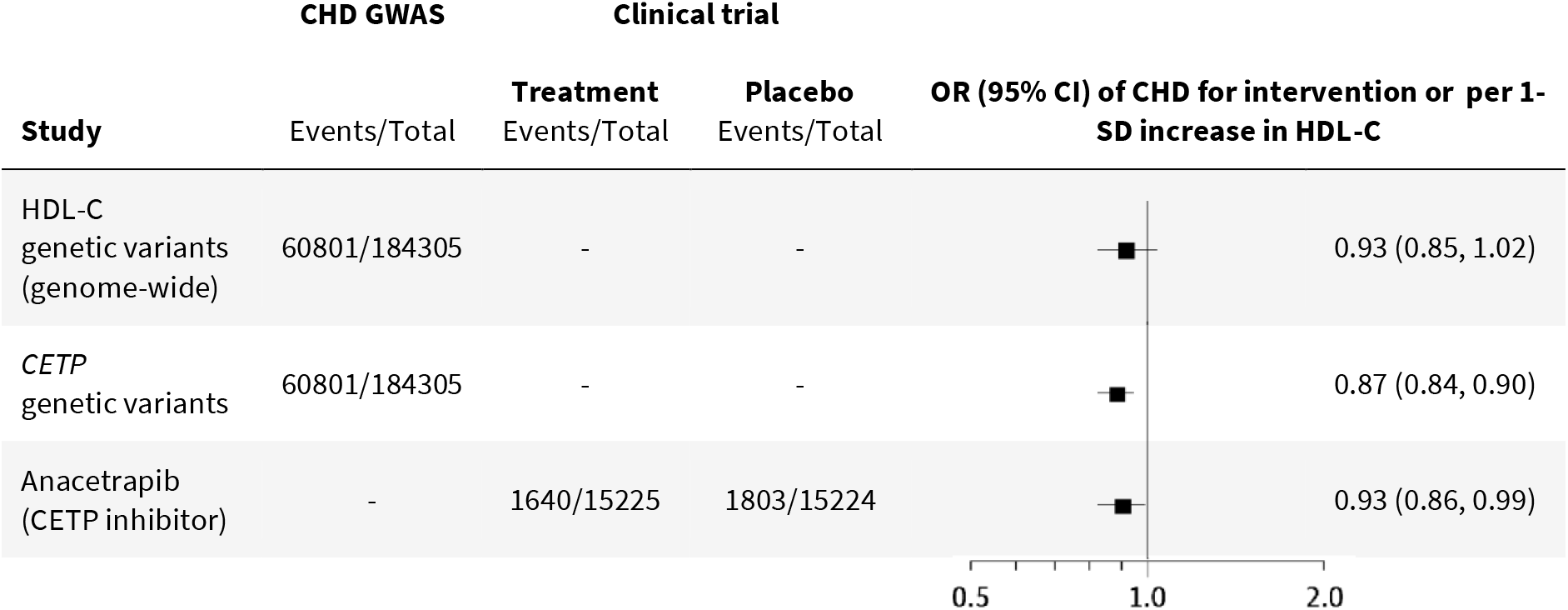
HDL-C, CETP inhibitor and CHD: biomarker vs drug target MR. Forest plot of the HDL-C biomarker MR estimate (Holmes et al, 2015), drug target MR estimate of CETP level and function using HDL-C as a proxy (Schmidt et al, 2020) and odds ratio of anacetrapib clinical trial (HPS3/TIMI55–REVEAL Collaborative Group, 2017). OR = odds ratio; CI = confidence interval; SD = standard deviation.

In this study, we applied drug target MR on a set of druggable proteins identified through genetic associations with circulating blood lipids, and assessed their causal relevance for CHD. To place the findings in context, we first re-evaluated causal effect estimates for LDL-C, HDL-C, and TG on CHD using *biomarker MR*, based on summary statistics from GWAS of blood lipids and CHD. Next, we used these data to select genes associated with blood lipids that encode druggable targets, and tested the effects of these drug targets on CHD using *drug target MR*. For a set of replicated, prioritized drug targets, we performed a phenome-wide scan of genetic associations of variants within the encoding gene with additional disease biomarkers and end-points beyond CHD. We sourced data from clinicaltrials.gov and the British National Formulary (BNF) for drugs in clinical phase development and approved medicines, respectively, to identify agents that might be pursued rapidly in clinical phase testing for treatment or prevention of CHD. Finally, we discuss how this approach might be generalized to other drug targets and clinical end-points, providing a route to translating findings from GWAS into new drug development.

## Results

### Biomarker MR analysis of LDL-C, HDL-C and TG on CHD

Previous biomarker MR studies have shown a causal effect of LDL-C and TG on CHD risk, while the causal role of HDL-C remains uncertain *(4)*. As an initial step, to confirm the robustness of our analytical pipeline and contextualize further analyses, we first replicated previously reported biomarker MR estimates using genetic variants from the Global Lipid Genetic Consortium (GLGC) *(13)* to instrument causal effects of the three lipid sub-fractions on CHD, using summary statistics from the CardiogramPlusC4D Consortium GWAS *(14)*. Causal estimates were obtained through univariable Mendelian randomization, with Egger horizontal pleiotropy correction applied through a model selection framework *(15)*. The odds ratio (OR) for CHD per standard deviation (SD) higher concentration of the corresponding blood lipid fraction was 1.50 (95% confidence interval (CI): 1.39, 1.63) for LDL-C, 0.95 (95% CI: 0.90, 1.01) for HDL-C and 1.10 (95% CI: 1.01, 1.21) for TG. These findings were replicated in an independent analysis using summary statistics from a GWAS meta-analysis of lipids measured using a nuclear magnetic resonance (NMR) spectroscopy platform *(16, 17)*, and genetic associations with CHD risk derived from UK Biobank *(18)*. The odds ratio for CHD per SD increase in LDL-C and TG in the replication dataset were 1.28 (95% CI: 1.25, 1.31) and 1.23 (95% CI: 1.14, 1.32), respectively, and 0.89 (95% CI: 0.83, 0.96) per SD increase in HDL-C. These genome-wide biomarker MR estimates confirmed the previously reported causal effect of LDL-C and TG on CHD risk but illustrate the equivocal role of HDL-C. To account for the correlation between the lipid fractions and evaluate their direct independent effect on CHD, we performed a multivariable MR (MVMR) analysis in the discovery datasets, which assessed genetic associations with the three lipid subfractions and CHD risk in a single model. The MVMR analysis generated an OR of 1.53 (95% CI: 1.44, 1.62) per SD higher LDL-C, 0.91 (95% CI: 0.86, 0.95) per SD higher HDL-C and 1.09 (95% CI: 1.01, 1.17) per SD higher TG (table S1).

### Drug target MR analysis

Drug target MR was used to determine the effect on CHD of perturbing druggable proteins that influence one or more of the three lipid fractions. First, genes previously shown to encode druggable proteins were selected in regions around variants associated with one or more of the major circulating lipid subfractions applying a *P* value < 1×10^−6^. This identified 341 genes; 149 for an association with LDL-C, 180 for HDL-C and 154 for TG *(19)*. One hundred forty genes (41%) were associated with a single lipid subfraction, 101 (30%) were associated with two subfractions and 100 (29%) were associated with all three subfractions (fig. S1, table S2). Subsequently, we performed a drug target MR analysis on CHD accounting for genetic correlation between variants (see Methods). In the absence of direct measures of the encoded protein, we proxied the effect of genetic drug target perturbation through the downstream effect on one or more of the three lipid sub-fractions. Of the 341 drug targets, the effect estimates for 165 excluded a null effect on CHD, with 131 of these estimates being consistent with a protective effect via decreasing LDL-C or TG and/or increasing HDL-C (Fig. 2, table S3). When weighted by LDL-C, eighty-seven targets showed a significant effect on CHD after orientating towards an increasing LDL-C direction, with the first and third quartiles (Q) of the CHD OR of 1.93 and 3.32. Similarly, the Q1 and Q3 after orientating the OR towards an increasing HDL-C direction were 0.22 and 0.53 for the 49 significant HDL-C instrumented targets, and for the 49 significant TG instrumented targets these were 1.95 and 4.35, respectively.

**Fig. 2.**
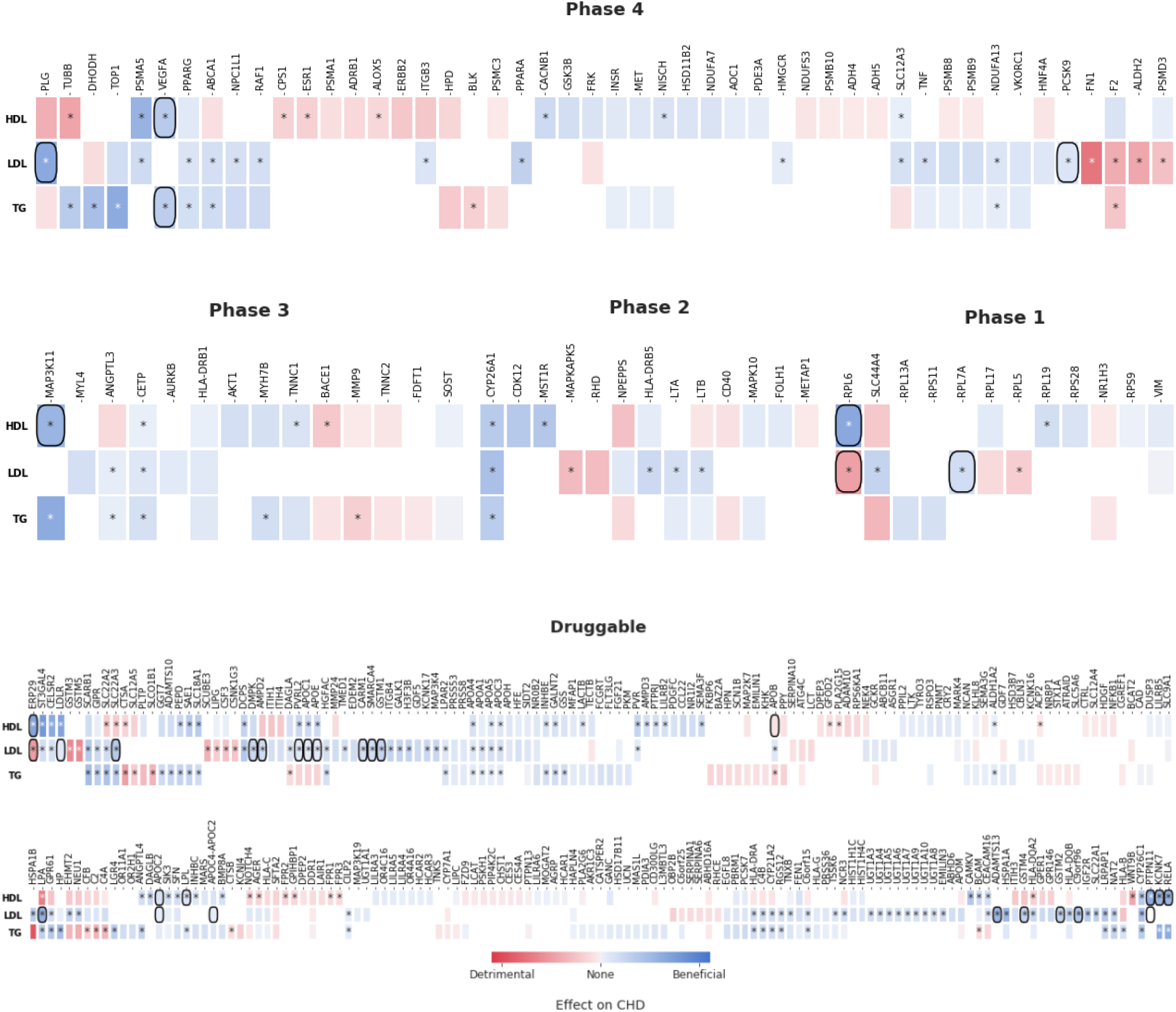
Drug target MR estimates on CHD. Analyses were performed using genetic associations with LDL-C, HDL-C and TG from the Global Lipid Genetic Consortium (GLGC) with CHD events from the CardiogramPlusC4D Consortium. Drug targets are grouped by clinical phase according to ChEMBL database. Blue indicates a beneficial effect on CHD risk, and red a detrimental effect per SD difference with respect to the indicated lipid sub-fraction. Significant estimates are indicated with an asterisk (*). Co-localization of genetic effects on the relevant lipid sub-fraction and CHD at the same locus is indicated by a square around the cell.

### Genetic rediscoveries of indications and mechanism-based adverse effects

We investigated if the drug target MR analysis rediscovered the mechanism of action of drugs with a license for lipid modification or compounds with a different indication but with reported lipid-related effects. To do so, compounds with reported lipid indications (CHD or non-CHD) or adverse effects were extracted from the BNF website (https://bnf.nice.org.uk/), which comprises prescribing information for all UK licensed drugs. Out of the 341 druggable genes included in the analysis, five encoded the targets of drugs with a lipid-modifying indication (PCSK9, PPARG, PPARA, NPC1L1, HMGCR) of which NPC1L1, HMGCR and PCSK9 are targets of drugs used in CHD prevention; and 6 encoded a protein target of a drug with reported lipid-related adverse effects (ADRB1, TNF, ESR1, FRK, BLK and DHODH) (table S4). To include outcome and side effect data of candidates in clinical phase development, the 341 drug targets were mapped to compound data available in a clinicaltrials.gov curated database. This database differentiates between endpoints monitored throughout the trial (‘outcomes’), and unanticipated harmful episodes during the study that may be on-target or off-target effects of the trial agent (‘adverse events’). Of the 341 drug targets, 23 had reported lipid related outcomes and 40 had reported lipid-related adverse events (table S4).

The pool of druggable targets that were modeled using higher LDL-C as a proxy for the pharmacological action on a drug target included 14 targets of clinically used drugs, three of which were licensed for CHD treatment by lowering LDL-C (HMGCR, PCSK9 and NPC1L1). The non-CHD indications of clinically used drugs included dyslipidemias (PPARA), type 2 diabetes (PPARG and NDUFA13), autoimmune diseases (TNF), neoplasms (RAF1 and PSMA5), circulatory disorders (ABCA1, PLG, ITGB3 and F2) and alcohol-dependency (ALDH2) (Table 2). With the exception of F2, instrumenting the target action through an higher LDL-C effect was associated with a higher CHD risk. Two drug targets were for compounds already in phase 3 trials for CHD prevention (ANGPTL3 and CETP). Their effect on CHD instrumented through an higher LDL-C effect was similar in magnitude to that observed for previously licensed drugs, with OR 1.21 (95%CI 1.11; 1.33) and 1.49 (95%CI 1.29, 1.72), respectively. Lastly, three targets were in phase 2 trials of compounds developed for other indications (CYP26A1, LTA and LTB). The remaining 82 of the 101 targets had not yet been drugged by compounds in clinical phase development.

**Table 2.**
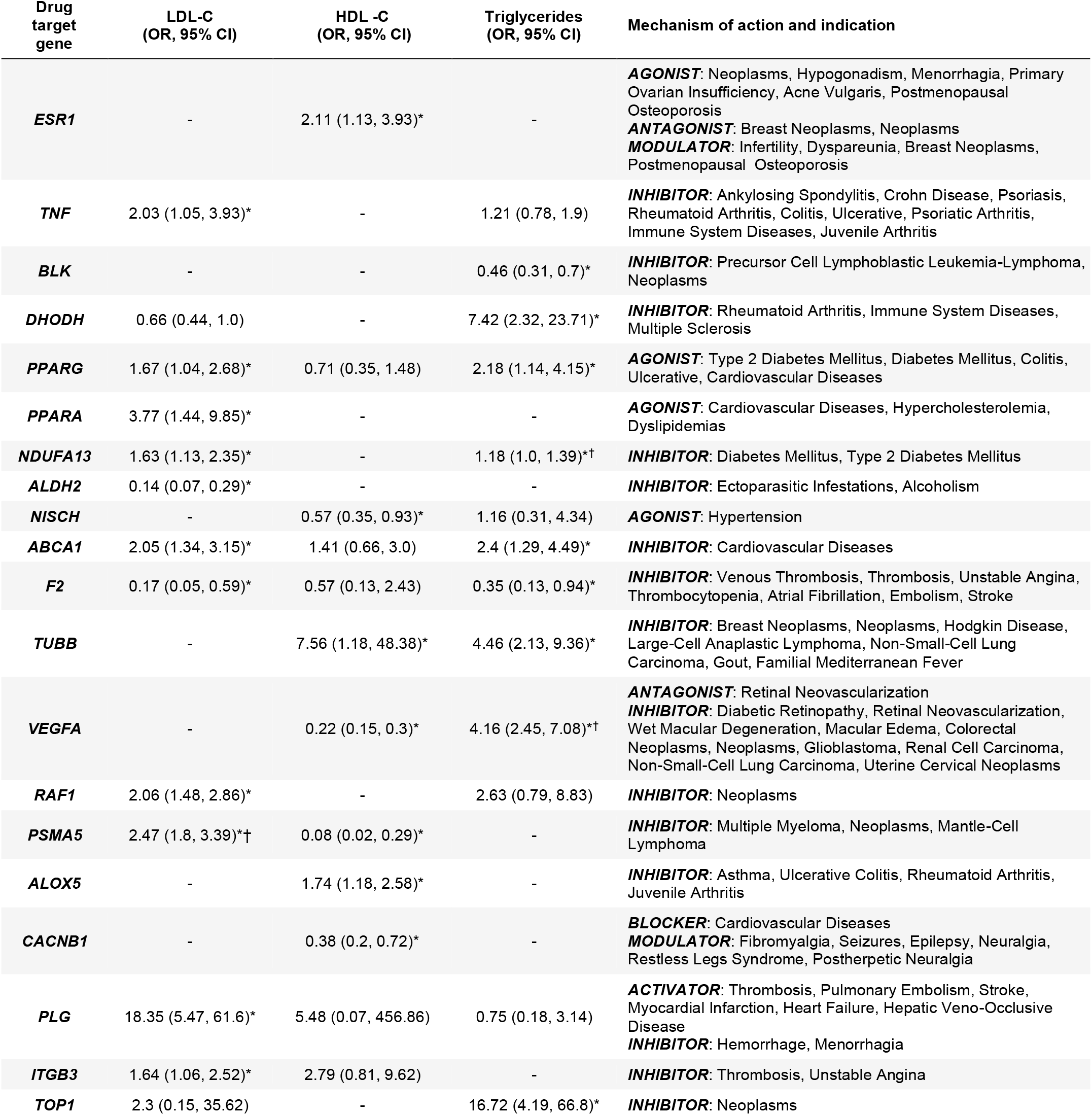
Univariable drug target MR estimates for drug targets approved for indications other than lipid-lowering. These drug targets showed lipid records in clinicaltrials.gov and/or the British National Formulary (BNF). **\***indicates significance in the discovery analysis; † indicates significance in both original and validation study and concordant direction of effect. OR = odds ratio of CHD per 1-standard deviation increase in LDL-C, HDL-C or triglycerides; CI = confidence interval.

| Drug target gene | LDL-C (OR, 95% CI) | HDL -C (OR, 95% CI) | Triglycerides (OR, 95% CI) | Mechanism of action and indication |
| --- | --- | --- | --- | --- |
| *ESR1* | - | 2.11 (1.13, 3.93)* | - | **AGONIST:** Neoplasms, Hypogonadism, Menorrhagia, Primary Ovarian Insufficiency, Acne Vulgaris, Postmenopausal Osteoporosis<br>**ANTAGONIST:** Breast Neoplasms, Neoplasms<br>**MODULATOR:** Infertility, Dyspareunia, Breast Neoplasms, Postmenopausal Osteoporosis |
| *TNF* | 2.03 (1.05, 3.93)* | - | 1.21 (0.78, 1.9) | **INHIBITOR:** Ankylosing Spondylitis, Crohn Disease, Psoriasis, Rheumatoid Arthritis, Colitis, Ulcerative, Psoriatic Arthritis, Immune System Diseases, Juvenile Arthritis |
| *BLK* | - | - | 0.46 (0.31, 0.7)* | **INHIBITOR:** Precursor Cell Lymphoblastic Leukemia-Lymphoma, Neoplasms |
| *DHODH* | 0.66 (0.44, 1.0) | - | 7.42 (2.32, 23.71)* | **INHIBITOR:** Rheumatoid Arthritis, Immune System Diseases, Multiple Sclerosis |
| *PPARG* | 1.67 (1.04, 2.68)* | 0.71 (0.35, 1.48) | 2.18 (1.14, 4.15)* | **AGONIST:** Type 2 Diabetes Mellitus, Diabetes Mellitus, Colitis, Ulcerative, Cardiovascular Diseases |
| *PPARA* | 3.77 (1.44, 9.85)* | - | - | **AGONIST:** Cardiovascular Diseases, Hypercholesterolemia, Dyslipidemias |
| *NDUFA13* | 1.63 (1.13, 2.35)* | - | 1.18 (1.0, 1.39)*† | **INHIBITOR:** Diabetes Mellitus, Type 2 Diabetes Mellitus |
| *ALDH2* | 0.14 (0.07, 0.29)* | - | - | **INHIBITOR:** Ectoparasitic Infestations, Alcoholism |
| *NISCH* | - | 0.57 (0.35, 0.93)* | 1.16 (0.31, 4.34) | **AGONIST:** Hypertension |
| *ABCA1* | 2.05 (1.34, 3.15)* | 1.41 (0.66, 3.0) | 2.4 (1.29, 4.49)* | **INHIBITOR:** Cardiovascular Diseases |
| *F2* | 0.17 (0.05, 0.59)* | 0.57 (0.13, 2.43) | 0.35 (0.13, 0.94)* | **INHIBITOR:** Venous Thrombosis, Thrombosis, Unstable Angina, Thrombocytopenia, Atrial Fibrillation, Embolism, Stroke |
| *TUBB* | - | 7.56 (1.18, 48.38)* | 4.46 (2.13, 9.36)* | **INHIBITOR:** Breast Neoplasms, Neoplasms, Hodgkin Disease, Large-Cell Anaplastic Lymphoma, Non-Small-Cell Lung Carcinoma, Gout, Familial Mediterranean Fever |
| *VEGFA* | - | 0.22 (0.15, 0.3)* | 4.16 (2.45, 7.08)*† | **ANTAGONIST:** Retinal Neovascularization<br>**INHIBITOR:** Diabetic Retinopathy, Retinal Neovascularization, Wet Macular Degeneration, Macular Edema, Colorectal Neoplasms, Neoplasms, Glioblastoma, Renal Cell Carcinoma, Non-Small-Cell Lung Carcinoma, Uterine Cervical Neoplasms |
| *RAF1* | 2.06 (1.48, 2.86)* | - | 2.63 (0.79, 8.83) | **INHIBITOR:** Neoplasms |
| *PSMA5* | 2.47 (1.8, 3.39)*† | 0.08 (0.02, 0.29)* | - | **INHIBITOR:** Multiple Myeloma, Neoplasms, Mantle-Cell Lymphoma |
| *ALOX5* | - | 1.74 (1.18, 2.58)* | - | **INHIBITOR:** Asthma, Ulcerative Colitis, Rheumatoid Arthritis, Juvenile Arthritis |
| *CACNB1* | - | 0.38 (0.2, 0.72)* | - | **BLOCKER:** Cardiovascular Diseases<br>**MODULATOR:** Fibromyalgia, Seizures, Epilepsy, Neuralgia, Restless Legs Syndrome, Postherpetic Neuralgia |
| *PLG* | 18.35 (5.47, 61.6)* | 5.48 (0.07, 456.86) | 0.75 (0.18, 3.14) | **ACTIVATOR:** Thrombosis, Pulmonary Embolism, Stroke, Myocardial Infarction, Heart Failure, Hepatic Veno-Occlusive Disease<br>**INHIBITOR:** Hemorrhage, Menorrhagia |
| *ITGB3* | 1.64 (1.06, 2.52)* | 2.79 (0.81, 9.62) | - | **INHIBITOR:** Thrombosis, Unstable Angina |
| *TOP1* | 2.3 (0.15, 35.62) | - | 16.72 (4.19, 66.8)* | **INHIBITOR:** Neoplasms |

When using higher HDL-C as a proxy for pharmacological action, MR of four drug targets with compounds approved for non-CHD indications showed a directionally beneficial effect on CHD (VEGFA, PSMA5, CACNB1 and NISCH), suggesting potential for indication expansion (Table 2). Three were targets for drugs approved for non-CHD indications but which showed a potentially detrimental effect direction on CHD when instrumented through increasing HDL-C concentration (ESR1, ALOX5, TUBB). Both CYP26A1 and CETP were associated with lower CHD risk when the effect on CHD was instrumented through an elevation of HDL-C. The remaining 65 of the 74 targets have not yet been drugged by compounds in clinical phase development.

Lastly, the set of druggable targets with compounds developed for non-CHD indications that were modeled using higher TG as a proxy for the pharmacological action on the target included PPARG, DHODH, VEGFA, TOP1, TUBB, NDUFA13, ABCA1, BLK, and F2 (Table 2). Of these, instrumenting the CHD effect through higher TG via drug action on BLK or F2 increased CHD risk. For the remaining targets, which included CETP, ANGPTL3 and CYP26A1, instrumenting the target effect through lowering TG levels decreased the risk of CHD, while the remaining 52 of the 64 targets have not been drugged by licensed compounds or clinical candidates yet.

### Independent validation of the drug target MR estimates

To help verify the MR findings, an independent two sample drug target MR analysis was conducted using summary statistics from a GWAS of blood lipids measured using an NMR spectroscopy platform *(16, 17)*, and genetic associations with CHD risk derived from UK Biobank *(18)*. The validation analysis identified 47 significant MR estimates (*P* value < 0.05), of which 39/47 (83%) showed a concordant direction of effect with the initial analysis (Table 3) corresponding to 30 drug targets. Replicated targets included the licensed LDL-lowering drug targets PCSK9 and NPC1L1 (Table 4). While the majority of the replicated drug targets were anticipated to decrease CHD risk via lowering LDL-C concentration based on the univariable results, 9 of the drug targets analyzed were significantly associated with lower CHD when the effects were modelled through HDL-C and/or TG (fig. S2).

**Table 3.**
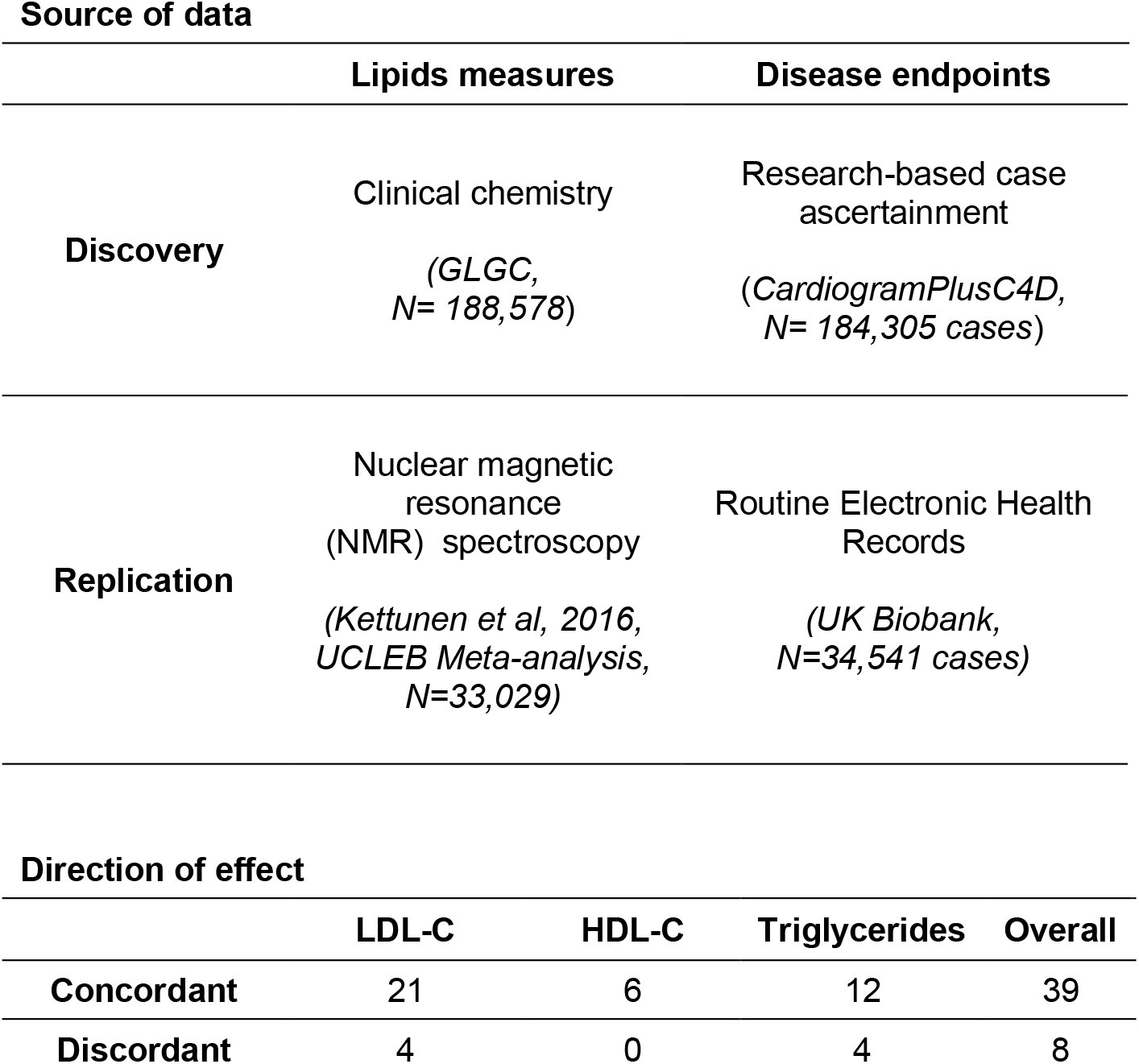
Replication of drug target MR findings. The discovery and replication analyses used different data sources for both exposure and outcome. 145 replication MR analyses were performed in which the gene boundaries included genetic associations exceeding the pre-specified significance threshold (*P* ≤ 1×10^−4^).

**Table 4.** Tissue specificity for replicated genes encoding drug targets. The tau value is a measure of tissue specificity with values between 0 and 1, where 1 indicates high specificity for a single tissue. The tissue with the highest expression of the gene is indicated in the top tissue column. * indicates significance in the discovery analysis, † indicates significance in both original and validation study and concordant direction of effect. OR = odds ratio of CHD per 1-standard deviation increase in LDL-C, HDL-C or triglycerides; CI = confidence interval.

| Drug target gene | LDL-C (OR, 95% CI) | HDL-C (OR, 95% CI) | Triglycerides (OR, 95% CI) | Tissue specificity index (tau) | Top tissues (Z-score >1) |
| --- | --- | --- | --- | --- | --- |
| **APOA5** | 2.05 (1.4, 3.02)*† | 0.72 (0.6, 0.87)*† | 1.21 (1.12, 1.31)*† | 1.00 | liver |
| **SLC12A3** | 1.94 (1.43, 2.63)* | 0.89 (0.86, 0.93)*† | 0.75 (0.24, 2.33) | 0.98 | kidney |
| **CEACAM16** | 1.66 (1.31, 2.11)*† | 0.46 (0.27, 0.79)* | 0.56 (0.25, 1.27) | 0.98 | pancreas, tonsil |
| **APOC3** | 2.04 (1.72, 2.42)*† | 0.67 (0.58, 0.78)* | 1.26 (1.12, 1.41)*† | 0.95 | liver |
| **APOA4** | 1.51 (1.23, 1.86)*† | 0.53 (0.38, 0.74)* | 1.27 (1.14, 1.43)*† | 0.94 | small intestine, colon, duodenum |
| **APOB** | 1.5 (1.18, 1.9)*† | 1.23 (0.72, 2.12) | 0.53 (0.29, 0.98)*† | 0.94 | liver, small intestine |
| **APOA1** | 1.88 (1.49, 2.36)*† | 0.84 (0.63, 1.11) | 1.25 (1.12, 1.4)*† | 0.93 | liver, small intestine |
| **NPC1L1** | 2.01 (1.48, 2.73)*† | - | 2.56 (0.75, 8.68) | 0.92 | small intestine, colon, duodenum, liver |
| **GPR61** | 1.97 (1.56, 2.5)*† | 3.02 (0.77, 11.91) | 5.14 (1.43, 18.48)* | 0.91 | cerebral cortex, adrenal gland, eye, thyroid gland |
| **PCSK9** | 1.6 (1.45, 1.77)*† | - | - | 0.87 | liver, lung, pancreas |
| **APOC1** | 1.31 (1.22, 1.41)*† | 0.39 (0.25, 0.59)* | 0.51 (0.17, 1.47) | 0.85 | liver |
| **CETP** | 1.49 (1.29, 1.72)* | 0.91 (0.87, 0.95)*† | 1.98 (1.63, 2.4)*† | 0.76 | lymph node, liver, placenta, spleen |
| **ADAMTS13** | 11.18 (4.37, 28.59)*† | - | - | 0.72 | liver |
| **CILP2** | 1.19 (1.01, 1.39)* | - | 1.18 (1.0, 1.39)*† | 0.71 | testis, gallbladder, ovary, thyroid gland |
| **LPL** | - | 0.63 (0.49, 0.82)*† | 1.68 (1.46, 1.92)*† | 0.68 | adipose tissue, breast, heart muscle, seminal vesicle |
| **APOE** | 1.3 (1.2, 1.41)*† | 0.39 (0.26, 0.59)* | 0.5 (0.17, 1.45) | 0.58 | liver, adrenal gland |
| **CELSR2** | 1.97 (1.78, 2.18)*† | 0.06 (0.04, 0.09)* | - | 0.58 | cerebral cortex, fallopian tube, skin |
| **ALDH1A2** | - | 0.89 (0.81, 0.99)* | 1.28 (1.07, 1.54)*† | 0.55 | endometrium, blood, cervix, uterine, fallopian tube, eye, seminal vesicle, testis |
| **ANGPTL4** | - | 0.48 (0.28, 0.83)*† | 3.38 (1.02, 11.22)*† | 0.50 | liver, adipose tissue, breast, cerebral cortex, pancreas |
| **PVR** | 1.31 (1.12, 1.54)*† | 0.32 (0.11, 0.91)* | - | 0.45 | liver, heart muscle |
| **NDUFA13** | 1.63 (1.13, 2.35)* | - | 1.18 (1.0, 1.39)*† | 0.43 | testis, blood, heart muscle, skeletal muscle |
| **CARM1** | 2.27 (1.68, 3.05)*† | - | - | 0.38 | skeletal muscle |
| **VEGFA** | - | 0.22 (0.15, 0.3)* | 4.16 (2.45, 7.08)*† | 0.33 | thyroid gland, endometrium, heart muscle, liver, skeletal muscle, urinary bladder |
| **SIK3** | 1.15 (0.57, 2.31) | 0.46 (0.29, 0.73)*† | 1.08 (0.98, 1.18) | 0.27 | cerebral cortex, ovary, parathyroid gland, testis, thyroid gland |
| **TMED1** | 2.06 (1.5, 2.83)*† | - | - | 0.26 | blood, heart muscle, liver, placenta, skeletal muscle |
| **PSMA5** | 2.47 (1.8, 3.39)*† | 0.08 (0.02, 0.29)* | - | 0.23 | liver, cerebral cortex, kidney, skeletal muscle, thyroid gland |
| **SMARCA4** | 2.22 (1.98, 2.49)*† | 0.01 (0.0, 0.02)* | - | 0.19 | cerebral cortex, bone marrow, esophagus, skeletal muscle, skin, testis, tonsil |
| **RPL7A** | 2.29 (1.57, 3.36)*† | - | - | 0.19 | salivary gland, endometrium, lymph node, ovary, pancreas |

### Discriminating independent lipid effects

After considering each lipid sub-fraction as a single measure on disease risk in the univariable drug target MR analyses, we performed a multivariable drug target MR analysis including LDL-C, HDL-C and TG in a single model to account for potential pleiotropic effects of target perturbation via the other lipid sub-fractions. Twenty-six of the replicated targets had sufficient data (3 or more variants) for the multivariable analysis. This analysis prioritized a single lipid fraction for 12 targets (SLC12A3, APOB, APOA1, PVRL2, APOE, APOC1, CELSR2, GPR61, PCSK9 and CEACAM16 through LDL-C; LPL through HDL-C; and ALDH1A2 through TG) (table S5). For SMARCA4 and APOA5, the analysis prioritized both LDL-C and TG, and for RPL7A both LDL-C and HDL-C. Due to the limited number of variants in VEGFA, CILP2, NDUFA13 and ANGPTL4, multivariable MR analysis could not distinguish the lipid fraction through which CHD was likely affected. Additionally, the presence of horizontal pleiotropy in the MVMR analysis based on heterogeneity tests suggested that PCSK9, LPL, APOC1, APOE, PVRL2, APOB, APOC3, CETP, APOA1 and CELSR2 may affect CHD through additional pathways beyond the lipid sub-fractions LDL-C, HDL-C and TG included in the current model.

### Co-localization between loci for lipids and CHD

Co-localization analyses are often performed to facilitate the mapping of genetic variants to causal genes in a disease GWAS by assessing whether associations with gene expression and disease outcome share a causal variant. Here, we applied co-localization analysis using blood lipids as an intermediate trait instead of gene expression data as a parallel validation step to assess if the genetic associations with each lipid sub-fraction and CHD were consistent with a shared causal variant *(20)*. Twenty-eight out of a total of 33 co-localization signals overlapped a significant finding in the discovery MR, which corresponded to 25 genes encoding a drugged or druggable target (Fig. 2). Moreover, 11 of the replicated drug targets showed evidence of co-localization between the lipid sub-fraction and CHD. These included 9 targets replicated for lowering LDL-C levels (SMARCA4, PVLR2, APOE, APOC1, CARM1, RPL7A, ADAMTS13, PCSK9 and C9orf96), one target replicated for raising HDL-C levels (LPL), and one target replicated for lowering TG levels (VEGFA).

### Tissue expression to aid drug target prioritization

While many tissues are involved in lipid homeostasis, the liver is considered the mechanistic effector organ for many therapeutics targeting lipid metabolism *(21)*. To investigate if the replicated drug target genes were specifically expressed in liver or any other particular tissue, we extracted their RNAseq expression profiles from the Human Protein Atlas *(22)* and calculated two commonly used tissue specificity metrics: the tau and z-scores *(23)*. Whilst tau summarizes the overall tissue distribution of a given gene and helps to distinguish between broadly expressed house-keeping genes (tau = 0) and tissue-specific genes (tau = 1), z-scores quantify how elevated the expression of a particular gene is in a particular tissue compared to other tissues. Among the 30 replicated genes, 28 had available RNAseq data, of which 15 (54%) showed elevated expression in the liver compared to other tissues (z-score > 1) (Table 4, fig. S3). These genes included the known lipid-lowering drug targets, PCSK9 and NPC1L1. Furthermore, eight genes were highly specific to the liver as indicated by high tau values (tau > 0.8). Other tissues showing elevated expression of the replicated drug target genes were gastrointestinal tissues such as small intestine and colon (e.g. APOA4, APOB) and kidney (SLC12A3). Regarding the expression distribution of the targets, 9 showed tau values below 0.5, indicating that they are broadly expressed and suggesting that, when developing a drug, the possibility of observing adverse effects increases *(24)*.

### Phenome-wide scan of replicated drug target candidates

The identification of potential mechanism-based adverse effects of a target represents an important aspect when prioritizing clinical candidates in the drug development pipeline. To explore potential effects of target perturbation on clinical endpoints other than CHD (whether beneficial or adverse), we performed a phenome-wide scan in 103 disease traits of the 30 drug targets replicated via drug target MR (Methods, Fig. 3, fig. S4-32). Besides genome-wide significant associations with diseases of the circulatory system, variants in six drug target genes showed genome-wide significant associations with type 2 diabetes (NDUFA13, CILP2, PVRL2, VEGFA, APOC1, LPL), five with Alzheimer’s disease (APOC1, PVR, PVRL2, APOE, CEACAM16), four with asthma (SMARCA4, CETP, VEGFA, ALDH1A2) and four with gout (APOA1, APOC3, APOA4, APOA5). Notably, the PheWAS rediscovered the mechanism of action of metformin, a drug targeting NDUFA13 and licensed for type 2 diabetes *(25)*.

**Fig. 3.**
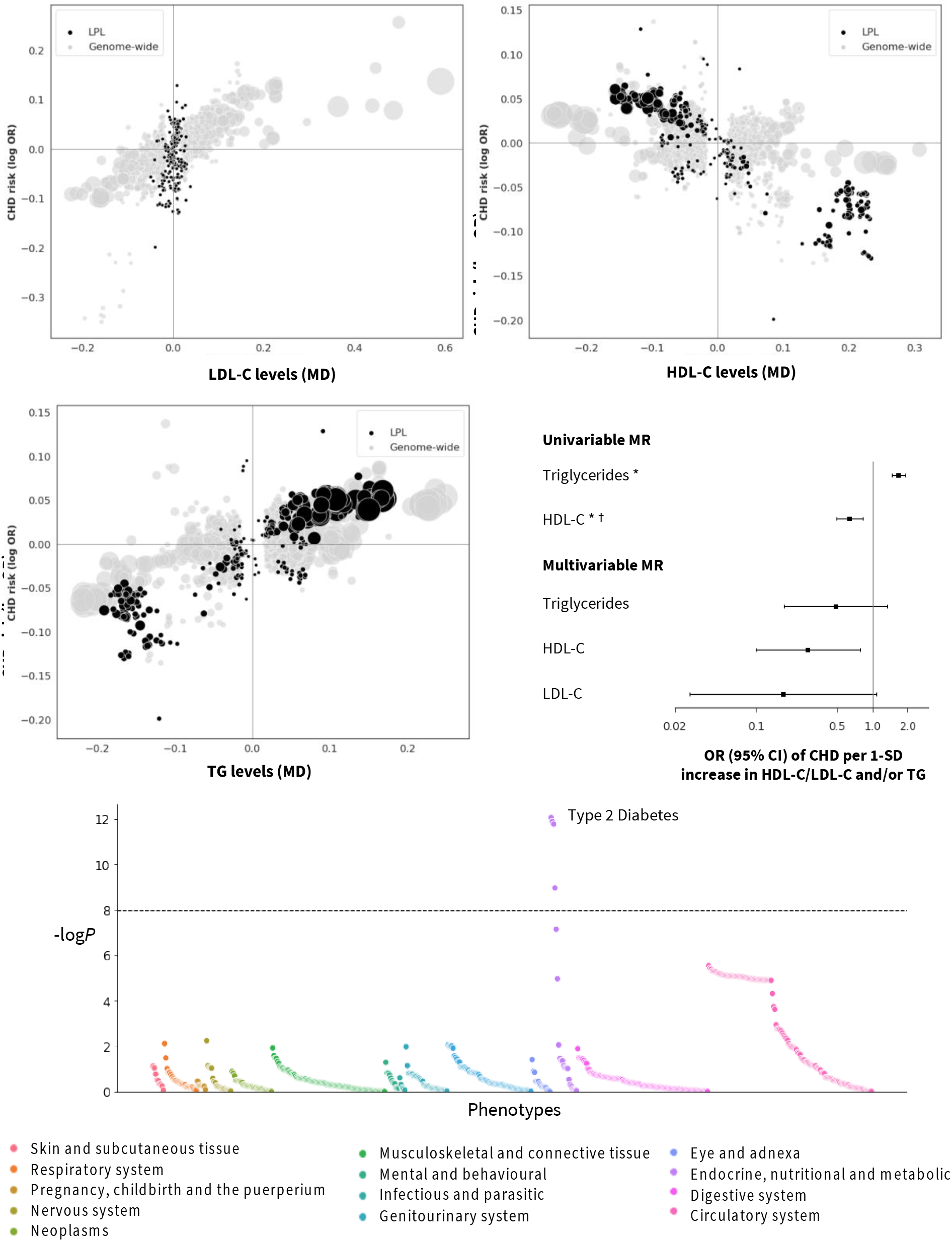
Prioritized target: lipoprotein lipase (LPL). The top and middle left panels show genetic associations at the locus (± 50kbp) in black vs genome-wide associations (grey, *P* value < 1×10^−6^). The x-axis shows the per allele effect on the corresponding lipid expressed as mean difference (MD) from GLGC and the y-axis indicates the per allele effect on CHD expressed as log odds ratios (OR) from CardiogramPlusC4D. The marker size indicates the significance of the association with the lipid sub-fraction (*P*-value). The middle right panel shows the result of the univariable and multivariable (drug target) *cis*-MR results. An asterisk (*) indicates the MR estimates as being replicated, and a dagger (†) that the lipid effect and CHD signals are co-localized. The bottom panel shows disease associations at the locus with 103 clinical end points from UK Biobank and GWAS Consortia.

## Discussion

By combining publicly available GWAS datasets on blood lipids and coronary heart disease and applying MR approaches with drug information and clinical data, we have genetically-validated and prioritized drug targets for CHD prevention.

We identified 131 drug target genes associated with CHD risk from a set of 341 druggable genes overlapping associations with one or more of the major blood lipid fractions.

Importantly, these effects were observed not only for genes associated with LDL-C, but also TG or HDL-C. The set of targets included NPC1L1, HMGCR and PCSK9, which are known targets of LDL-lowering drugs, whose efficacy in CHD prevention has been proven in clinical trials. We performed an independent replication study both to corroborate the targets and the direction of the effects. We replicated the findings in independent datasets (UCLEB Consortium and UK Biobank) in which lipids were measured using a different platform (NMR spectroscopy in UCLEB) and the disease end-points ascertained by linkage to routinely recorded health data (UK Biobank). The validation study replicated 83% (39/47) of the initial estimates, including the mechanism of action of current lipid-modifying drug targets PCSK9 and NPC1L1 and the suggested mechanism of action of compounds under investigation for lipid modification through TG or HDL-C, such as CETP inhibitors *(26, 27)*.

As a positive control step, our (genome-wide) biomarker MR analysis replicated previous findings on the potential causal relevance of LDL-C, TG, and HDL-C *(4, 10, 28)*. Importantly, contrary to previous studies, here we replicated findings using a completely independent set of NMR-spectroscopy measured lipids data and CHD cases sourced from UK Biobank. While the causal relevance of LDL-C for CHD has been robustly proven through successful drug development of for example statins, there are as yet no compounds licensed for CHD prevention through effects on HDL-C and TG. Hence, the causal relevance of the lipid sub-fraction, while supported by the current biomarker analyses, cannot be concluded definitively. It is therefore essential to highlight that, while our drug target analysis uses genetic associations with these lipid sub-fractions as weights, our inference throughout has been on the therapeutic relevance of perturbing the proteins encoded by the corresponding genes which are the main category of molecular target for drug action. The genetic associations with the corresponding lipids are merely used as a proxy for protein activity and/or concentration, serving to orientate the MR effects in the direction of a therapeutic effect. They do not provide comprehensive evidence on the pathway through which perturbation of such targets causally affects CHD. Nevertheless, co-localization and multivariable MR do provide insight on the potential relevance of lipid pathways in mediating the effects of drug target perturbation. Due to the potential for weak instrument bias, attenuating results towards the null, non-significant results should not be over-interpreted as proof of absence *(29)*.

The set of 30 replicated drug targets also included lipoprotein lipase (LPL), a target that could potentially decrease CHD risk through both TG-lowering and HDL-C elevation, with an effect through HDL-C further endorsed by the co-localization and multivariable MR analyses (Fig. 6). In contrast to current lipid-lowering drug targets which are specifically expressed in the liver, LPL shows highest specific expression in adipose tissue which suggests tissues beyond the liver may be relevant to target lipid metabolism. Several pharmacological attempts have been pursued to target LPL *(30, 31)*, and gene therapy has also been applied to treat LPL deficiency by introducing extra copies of the functional enzyme in patients with hypertriglyceridemia *(32)*. The approval of gene therapy interventions and the known indirect activation of LPL by drugs targeting other proteins, such as fibrates *(33)* and metformin *(34)*, suggest that the previous failure of compounds targeting LPL in initial trials may have been idiosyncratic. LPL activity is also modulated by another protein in the replicated dataset, apolipoprotein A5 (ApoA5), which is exclusively expressed in liver tissue. While the univariable drug target MR analysis of ApoA5 suggested that all three sub-fractions affected by ApoA5 perturbation may contribute to the effect on CHD risk, the multivariable MR suggest that ApoA5 (partially) affects CHD through LDL-C and TG-mediated pathways. Regardless of the mediating lipid or lipids, the genetic findings in relation to both LPL and ApoA5 are consistent and point to this as an important potentially targetable pathway in atherosclerosis, supporting prior work *(35)*.

To provide an indicative genetic profile of a drug target and hypothesise about potential mechanism-based adverse effects, repurposing opportunities or expansion of the indication portfolio of a drug target, we performed a PheWAS of variants in and around the replicated set of targets on 103 traits. While in some cases PheWAS highlighted associations with particular clinical endpoints, for example, the rediscovery of already known indications or biological pathway, such as the associations of type 2 diabetes with variants in *NDUFA13* or the association of Alzheimer’s Disease with *APOE*, further research is needed to evaluate the causal role of the target in the corresponding disease and the beneficial or detrimental effects of modulating those targets pharmacologically.

Some limitations of this study are noteworthy. First, we only included genes regarded as encoding druggable proteins, which currently comprise approximately 25% of all protein coding genes *(19)*. As new knowledge advances, additional proteins will become druggable, and alternative therapeutic strategies such as antisense oligonucleotides and gene therapy may extend the range of mechanisms that can be targeted. The approach we describe is in fact agnostic to therapeutic modality and could be adapted accordingly. Notably, antisense oligonucleotides efficiently delivered to the liver *(36)*, where 54% of the prioritised targets in our analysis showed elevated expression compared to other tissues. Second, we assigned variants to druggable genes based on genomic proximity, which may be as reliable as other approaches in mapping causal genes *(37–39)*. However, simple genomic proximity might result in misleading assignment of the causal gene in a region containing multiple genes in high LD (e.g. *PVRL2*, *APOC1* and *APOE* are all located in a region of LD in Chr19:45349432-45422606, GRCh37). In an effort to account for this, all the druggable genes (± 50kbp) that overlap one of the genetic variants associated with LDL-C, HDL-C or TG were included in the analysis, and we provided information on proximity of the variant to the gene, a gene distance rank value (in base pairs), and previous gene prioritisation data by the Global Lipids Genetics Consortium (GLGC) *(13)* to inform scenarios in which the causal gene may be a non-druggable gene but reside in the same region (table S2).

We used *cis*-MR to evaluate the relevance of each drug target to CHD, which poses additional challenges and choices: defining the locus of interest, the significance threshold for the association with the exposure and the LD threshold to prune correlated instruments. Since an agreement on the choice of a general LD threshold and flanking region has yet to be reached, we used a window of 50kbp and LD threshold of 0.4, which showed the most consistent estimates in a grid-search in the discovery data using the four positive control examples: PCSK9, NPC1L1, HMGCR and CETP. Based on previous studies showing that using less stringent P-value thresholds often results in improved performance in *cis*-MR settings, we relaxed the threshold below genome-wide significance to select the genetic associations to instrument the exposure; and accounted for LD correlation by pruning and LD modelling during the MR analysis *(12, 40)*.

To validate our findings with independent data sources, we conducted a second drug target MR, although several drug target genes could not be evaluated in the validation analysis because the gene boundaries did not include genetic associations exceeding the pre-specified significance threshold (*P* ≤ 1×10^−4^), likely related to the “modest” sample size of the NMR replication data (N=33,029). Beyond univariable MR analyses, we attempted to further validate the findings with a multivariable extension of the inverse-variance weighted (IVW) and MR Egger methods, however, in some cases we observed imprecise estimates in line with previous studies which attributed this to the inclusion of highly correlated exposures in the model *(41)*. To further evaluate if the association signals in the exposure and outcome datasets shared a causal genetic variant, we performed co-localization analyses. Because these analyses were originally developed to find evidence of co-localization between mRNA expression and a disease and not for an intermediate trait and a disease, the default prior probabilities used in the analysis may not be the optimal for these pairs of traits. In addition, the single-causal-variant assumption in genetic co-localization methods may not always be satisfied even when prior conditional analyses are performed, with regions with multiple causal variants potentially yielding false negative results *(42)*.

The effect directions of the replicated drug targets were compared to results from clinical trials using data from the clinicaltrials.gov registry, however, the lack of precision in annotation of events associated with lipid perturbations (e.g. hyperlipidaemia) in this dataset hinders the assignment of reported lipid abnormalities to a particular lipid sub-fraction. Moreover, the proportion of clinical trials with reported results in clinicaltrials.gov is less than 54.2% *(43)*, suggesting that additional drug candidates with lipid effects might have been investigated but were not included in this analysis because of the lack of accessible data. Furthermore, our analysis relied on mapping clinical trial interventions to compounds known to act through binding to the targets of interest, which could potentially miss clinical trials of compounds annotated with less synonyms (such as research codes for compounds used by individual trial sponsors). Lastly, we performed a PheWAS spanning over 100 clinical endpoints, 80 of which were derived from UK Biobank. While this enabled screening for associations with a wide range of diseases, genetic associations derived from diagnostic codes in electronic health record datasets might suffer from limited case numbers and inaccurate case and control definitions, which would reduce the power to detect true associations. To increase the power to detect associations, we included data from publicly available GWAS with the largest sample sizes for such phenotypes.

In summary, we have shown an approach to move from GWAS signals to drug targets and disease indications. We illustrated its potential using genetic association data on lipids and CHD data, but the approach could also be applied in other settings where there are GWAS of diseases and biomarkers thought to be potentially causally related. For example, with the increasing available data on inflammatory biomarkers, this approach could be used to evaluate the causal role of anti-inflammatory drug targets, such as IL6R, in CHD, Alzheimer’s disease and major depression, following up on associations described in several studies *(44–46)*, to identify potential new indications for anti-inflammatory agents established in the treatment of autoimmune conditions. Similarly, recent genetic studies on coagulation factor levels *(47)* can be harnessed to instrument the effect of modulating druggable targets for thrombotic disorders, such as FXI or FXII, which are emerging as potential targets for new anticoagulant drugs *(48, 49)*.

When used as a screening tool, the approach could help reduce the efficacy problem in drug discovery by genetically validating targets in the earlier phases of the drug development pipeline.

## Materials and Methods

### Data sources

To determine the causal role and replicate previously reported results on the causal effect of LDL-C, HDL-C and TG on CHD, we obtained genetic estimates from the Global Lipids Genetics Consortium (188,577 individuals) *(13)* and from CardiogramPlusC4D (60,801 cases and 123,504 controls) *(14)*.

Independent replication data were sourced using lipids exposure data from a GWAS meta-analysis of metabolic measures by the University College London–Edinburgh-Bristol (UCLEB) Consortium *(50)* and Kettunen *et al. (13)* utilizing NMR spectroscopy measured lipids (joint sample size up to 33,029). Independent CHD data was obtained from a publicly available GWAS of 34,541 cases and 261,984 controls in UK Biobank *(18)*.

### Drug target gene selection

Analyses were conducted using Python v3.7.3. To estimate the causal effect of modulating the level of each lipid sub-fraction via a druggable gene on CHD, variants associated with LDL-C, HDL-C and/or TG with a *P* ≤ 1×10^−6^ were selected. Druggable genes overlapping a 50kbp region around the selected variants were extracted, resulting in 341 associated drug target genes (149 for LDL-C, 180 for HDL-C and 154 for TG). The set of genes in the druggable genome were identified as described previously *(19)*, and identifiers were updated to Ensembl version 95 (GRCh37), used in this analysis. All of these IDs were also present in Ensembl 95 (GRCh37), used in this analysis. Because we only scanned for genetic associations with the druggable genome, protein-coding genes that were the ‘true’ causal gene but not yet druggable would be missed and the association mis-assigned. To mitigate this and provide information about potential effects through non-druggable genes, we provide the minimum distance from the variant to the gene, where variants located within a gene were given a distance of 0bp, a gene distance rank value according to their base pair distance, and indicated the druggable genes prioritized by GLGC (table S2).

### Instrument selection

For the biomarker or genome-wide MR analyses, a *P* threshold of 1×10^−6^ was used to select exposure variants associated with LDL-C, HDL-C and/or TG. For *cis*-or drug target MR analyses, variants from/within the 341 selected genes (±50kbp) were selected based on a *P* ≤ 1×10^−4^. In both settings, variants were filtered on a MAF > 0.01 and LD clumped to an r^2^ 0.4. These parameters showed the most consistent estimates in a grid-search in the discovery data using the positive control examples: PCSK9, NPC1L1, HMGCR and CETP (fig. S33). To account for residual correlation between variants in the MR analyses, we applied a novel generalized least squares framework with a LD reference dataset derived from UK Biobank *(51)* as described in Detailed materials and methods.

### Statistical analysis

As a validation step, a biomarker MR analysis was conducted for each lipid sub-fraction to replicate previous findings using genetic associations across the genome. A model-selection framework was used to decide between competing inverse-variance weighted (IVW) fixed-effects, IVW random-effects, MR-Egger fixed effects or MR-Egger random-effects models *(15)*. While IVW models assume an absence of directional horizontal pleiotropy, Egger models allow for possible directional pleiotropy at the cost of power. After removing variants with large heterogeneity (*P* < 0.001 for Cochran’s Q test) or leverage, we re-applied this model selection framework and used the final model (Detailed materials and methods).

Additionally, we conducted biomarker and drug target multivariable MR analyses using genetic associations with the three lipid sub-fractions and CHD risk in a single regression model.

### Co-localization analysis

To estimate the posterior probability of each druggable gene sharing the same causal variant for the exposure lipid and CHD risk *(52)* we performed co-localization analyses. First, we conducted a stepwise conditional analysis using GCTA-COJO v1.92.4 with genotype data from 5,000 individuals randomly selected from UK Biobank *(53)*. Colocalization analyses were performed using a Python implementation of “coloc” v3.2-1 *(20)*. The default prior probabilities were used to estimate if a SNP was associated only with the lipid sub-fraction (*p_1_*= 10^−4^), only with CHD risk (*p_2_* = 10−4), or with both traits (*p_12_* = 10^−5^). For each drug target gene, all variants from/within the gene boundaries (±50kbp) with a MAF > 0.01 were included. A posterior probability above 0.8 was considered sufficient evidence of colocalization based on previous observations *(20)*.

### Drug indications and adverse effects

To evaluate if the drug target MR and colocalization analyses rediscovered known drug indications, adverse effects or predicted repurposing opportunities, drug information and clinical trial data was extracted for the set of 341 druggable targets. Drug target genes were mapped to UniProt identifiers and indications and clinical phase for compounds that bind the target were extracted from the ChEMBL database (version 25) *(54)*. Drug indications and lipid adverse effects data for licensed drugs were extracted from the British National Formulary (BNF) website (https://bnf.nice.org.uk/) in July, 2019.

To further examine the effects of the drugs and clinical candidates that are known to act through binding to the 341 druggable targets, relevant clinical trial data were downloaded from the clinicaltrials.gov registry. Compound name and synonyms were extracted from ChEMBL database (version 25) *(54)* and used to identify clinical trials with matching interventions. In case of non-exact matches, the results were inspected manually to ensure that only relevant trial records were used in the analysis. Lipid-related trial outcomes and adverse events were identified by searching the relevant fields within the trial records with the keywords: lipo*, lipid*, ldl*, hdl*, cholest* and triglyceride*. For adverse events, the search was limited to the trial arm in which the drug of interest was administered (as opposed to placebo or active control used in the study) and only adverse events that affected at least one study participant were included.

### Tissue expression analysis

To further characterize the genes prioritized by the MR pipeline, their tissue expression was analyzed as follows. First, RNAseq data were downloaded from Human Protein Atlas (HPA) *(22)*, which captures baseline expression of human genes and proteins across a panel of diverse healthy tissues and organs. For each included gene and tissue, HPA provides a consensus Normalized eXpression value (NX), obtained by normalizing TPM (transcripts per million) values from three independent transcriptomics datasets: GTEx *(55)*, Fantom5 *(56)*, and HPA’s own RNAseq experiments *(57)*.

The downloaded NX values were then used to investigate if the prioritized target genes were specifically expressed in any of the included tissues. Two commonly used tissue specificity metrics were calculated for each gene: tau and z-score *(23)*. Tau summarizes the overall tissue distribution of a given gene and ranges from 0 to 1, where 0 indicates ubiquitous expression across all included tissues (house-keeping genes) and 1 indicates narrow expression (highly tissue-specific genes). While tau provides a single summary measure of the tissue specificity, z-scores are calculated for individual tissues separately to quantify how elevated the gene expression is in a particular tissue compared to others. Here, higher z-score values indicate higher tissue specificity. See Kryuchkova-Mostacci *et al. (23)* for details on the calculation and interpretation of the two metrics.

### Phenome-wide scan of replicated drug target genes

To explore the effect spectrum associated to prioritized drug targets, we performed a phenome-wide scan of 103 disease endpoints. These included genome-wide summary statistics for 80 ICD10 main diagnoses in UK Biobank, which were released by Neale Lab (1st August 2018, http://www.nealelab.is/uk-biobank/), and downloaded using a Python implementation of MR Base API *(58)*. The variants in-and-around the prioritized drug target genes allowing for a boundary region of 50kbp were extracted, palindromic variants were inferred using the API default MAF threshold of 0.3 and removed *(59)*. The Ensembl REST Client was used to gather positional information for the variants *(60)*.

We attempted to maximize the power to detect genetic associations by sourcing data from 23 publicly available GWAS with the largest sample sizes for such phenotypes (table S6). All the GWAS clinical endpoints and UK Biobank ICD10 main diagnoses were grouped according to ICD10 chapters.

## Supplementary Materials

Fig. S1. Overlap between genes encoding druggable targets associated with the major lipid sub-fractions.

Fig. S2. The sets of assigned genes associated with LDL-C, HDL-C, TG that encode druggable targets.

Fig. S3. Tissue expression profile of the replicated drug target genes.

Fig. S4. Prioritized target: SMARCA4

Fig. S5. Prioritized target: APOC1.

Fig. S6. Prioritized target: NPC1L1.

Fig. S7. Prioritized target: SLC12A3.

Fig. S8. Prioritized target: PVR.

Fig. S9. Prioritized target: APOB.

Fig. S10. Prioritized target: CETP.

Fig. S11. Prioritized target: TMED1.

Fig. S12. Prioritized target: APOA5.

Fig. S13. Prioritized target: APOA4.

Fig. S14. Prioritized target: APOC3.

Fig. S15. Prioritized target: VEGFA.

Fig. S16. Prioritized target: APOA1.

Fig. S17. Prioritized target: ALDH1A2.

Fig. S18. Prioritized target: PVRL2.

Fig. S19. Prioritized target: APOE.

Fig. S20. Prioritized target: CARM1.

Fig. S21. Prioritized target: PSMA5.

Fig. S22. Prioritized target: CELSR2.

Fig. S23. Prioritized target: RPL7A.

Fig. S24. Prioritized target: GPR61.

Fig. S25. Prioritized target: CILP2.

Fig. S26. Prioritized target: ADAMTS13.

Fig. S27. Prioritized target: ANGPTL4.

Fig. S28. Prioritized target: SIK3.

Fig. S29. Prioritized target: PCSK9.

Fig. S30. Prioritized target: C9orf96.

Fig. S31. Prioritized target: NDUFA13.

Fig. S32. Prioritized target: CEACAM16.

Fig. S33. Drug target MR of positive control examples.

Table S1. Causal odds ratios (95% CI) for CHD per standard deviation increase in each lipid sub-fraction from a biomarker MR analysis.

Table S2. Proximity to GWAS SNP, distance rank and previous evidence of druggable genes near genetic associations with LDL-C, HDL-C and TG.

Table S3. Univariable drug target MR estimates.

Table S4. Univariable MR estimates of drug targets with lipid records in clinicaltrials.gov and/or BNF.

Table S5. Multivariable drug target MR estimates.

Table S6. Publicly available GWAS data used in the PheWAS.

## Acknowledgments

The authors are grateful to the studies and consortia that provided summary association results and to the participants of the biobanks and research cohorts. This research has been conducted using the UK Biobank Resource under Application Number 12113. UK Biobank was established by the Wellcome Trust medical charity, Medical Research Council, Department of Health, Scottish Government, and the Northwest Regional Development Agency. It has also had funding from the Welsh Assembly Government and the British Heart Foundation.

## Author contributions

MGM, ADH, AFS and CF contributed to the idea and design of the study. MGM performed the Mendelian randomization and PheWAS analyses. MGM and MZ performed the tissue expression and clinical trial analyses. MGM, ADH, AFS, CF and drafted the manuscript. MGM, MZ, ADH, AFS and CF contributed to the first draft of the manuscript. All authors contributed to and approved the final version of the manuscript.

## Data and materials availability

All data are publicly available, as described in the methods section. Please contact MGM for access to specific files, data, or analysis scripts.

