## Supplementary Materials for "Validation of lipid-related therapeutic targets for coronary heart disease prevention using human genetics"

### Detailed materials and methods

*Linkage disequilibrium reference dataset*

LD reference matrices were created by extracting a random subset of 5,000 unrelated individuals of European ancestry from UK Biobank. Variants with a MAF < 0.001, and imputation quality < 0.3 were excluded. To ensure that SNPs with lower MAF have higher confidence, variants were removed if MAF < 0.005 and genotype probability < 0.9; MAF < 0.01 and genotype probability < 0.8; MAF < 0.03 and genotype probability < 0.6.

*Extended statistical analysis*

A drug target MR analysis was applied to the set of 341 genes using the model selection framework and heterogeneity tests described in the Methods section. The influence of parameter selection in the drug target MR performance was explored in a grid-search of several $r^2$ and gene boundaries combinations using the positive control examples PCSK9, NPC1L1, HMGCR and CETP, where the lipid perturbation is the intended indication. The estimates were consistent in the discovery and validation analysis (fig. S33), with the exception of HMGCR which showed inconsistencies when using NMR-spectroscopy measured lipid data and CHD risk derived from UK Biobank.

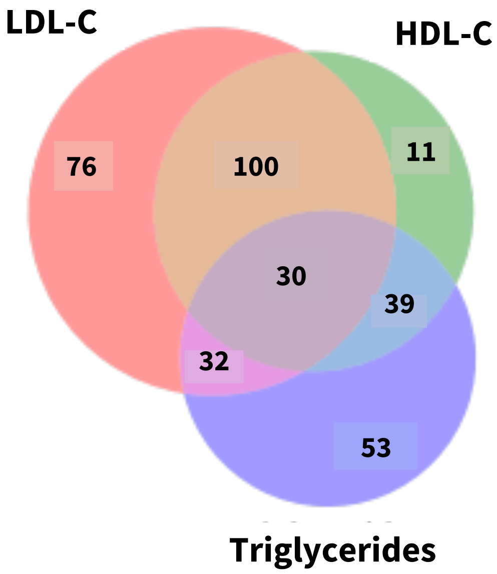


**Fig. S1**. **Overlap between genes encoding druggable targets associated with the major lipid subfractions.**

The Venn diagram shows genes exhibiting overlapping or exclusive associations with LDL-C, HDL-C and/or TG.

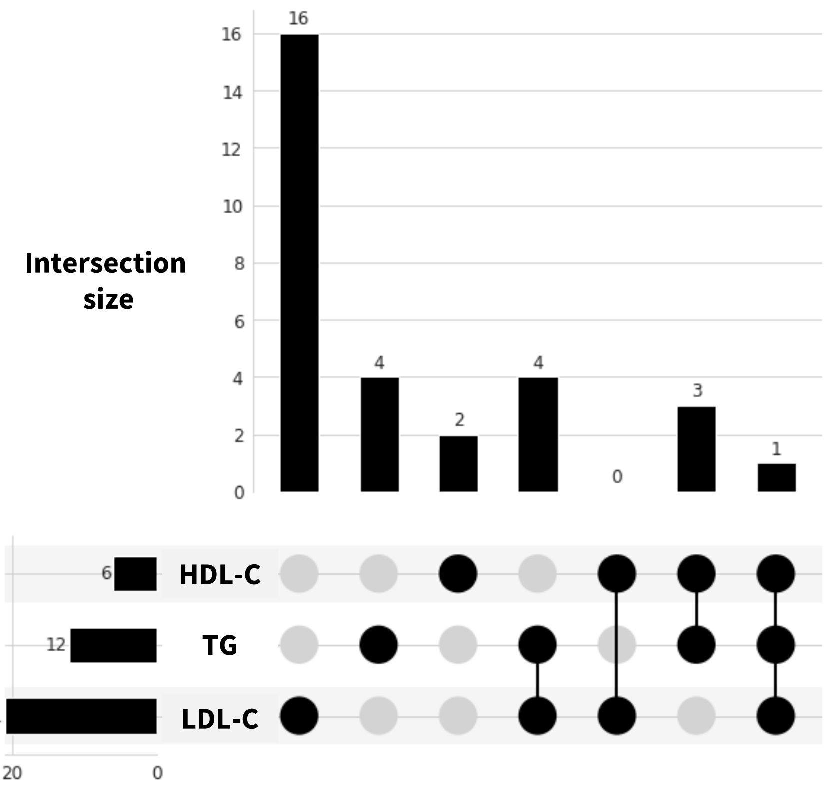


**Fig. S2. The sets of assigned genes associated with LDL-C, HDL-C, TG that encode druggable targets.** Genes encoding druggable targets were included if they demonstrated concordant direction of effect in the discovery and validation studies on CHD showing a causal effect of one or more lipid sub-fractions.

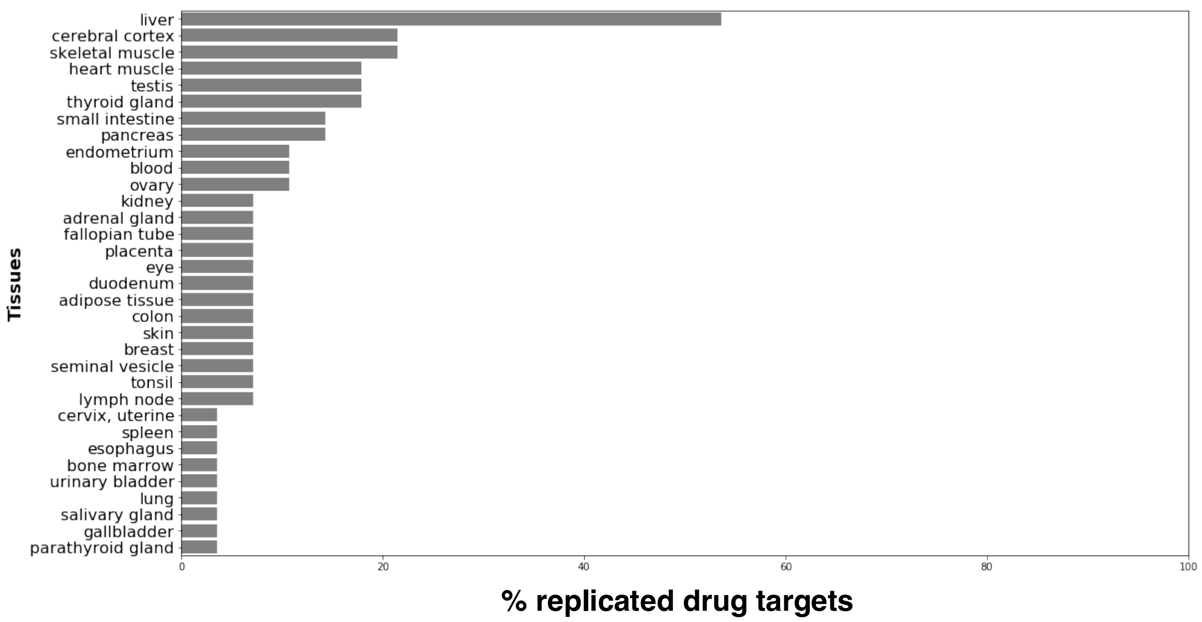


**Fig. S3. Tissue expression profile of the replicated drug target genes.** The tissue specificity metric, z-score, which quantifies how elevated the gene expression is in a particular tissue compared to others, was calculated for the set of 30 replicated genes encoding a druggable target. Of the 28 genes with RNAseq data available, 15 genes in the validated set (54%) were highly expressed in liver (z-score > 1), 13 (46%) did not have elevated liver expression.

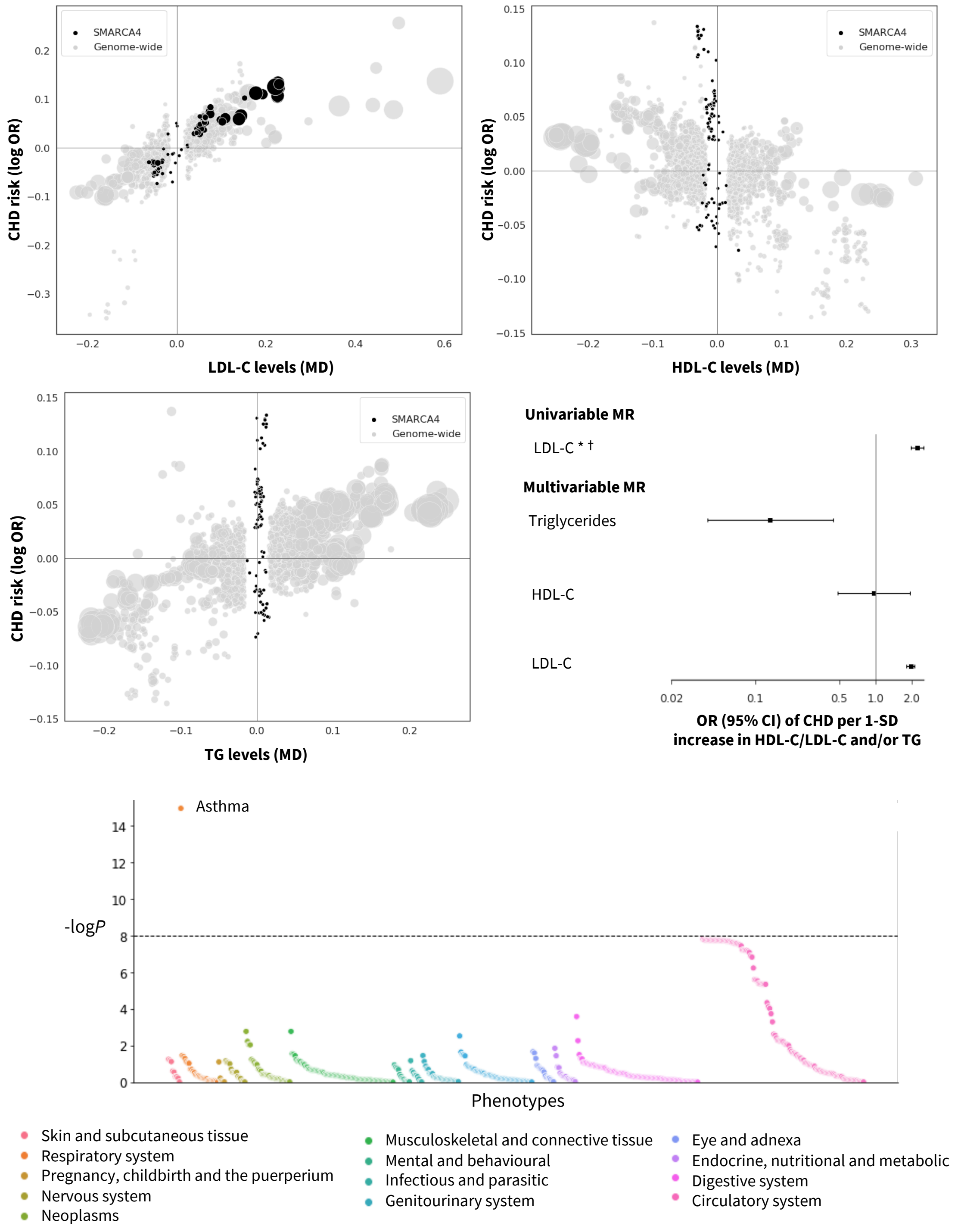


**Fig. S4. Prioritized target: SMARCA4.** The top and middle left panels show genetic associations at the locus (± 50kbp) in black vs genome-wide associations (grey, *P* value < $1 \times 10^{-6}$). The x-axis shows the per allele effect on the corresponding lipid expressed as mean difference (MD) from GLGC and the y-axis indicates the per allele effect on CHD expressed as log odds ratios (OR) from CardiogramPlusC4D. The marker size indicates the significance of the association with the lipid sub-fraction (*P*-value). The middle right panel shows the result of the univariable and multivariable (drug target) *cis*-MR results. An asterisk (*) indicates the MR estimates as being replicated, and a dagger (†) that the lipid effect and CHD signals are co-localized. The bottom panel shows disease associations at the locus with 103 clinical end points from UK Biobank and GWAS Consortia.

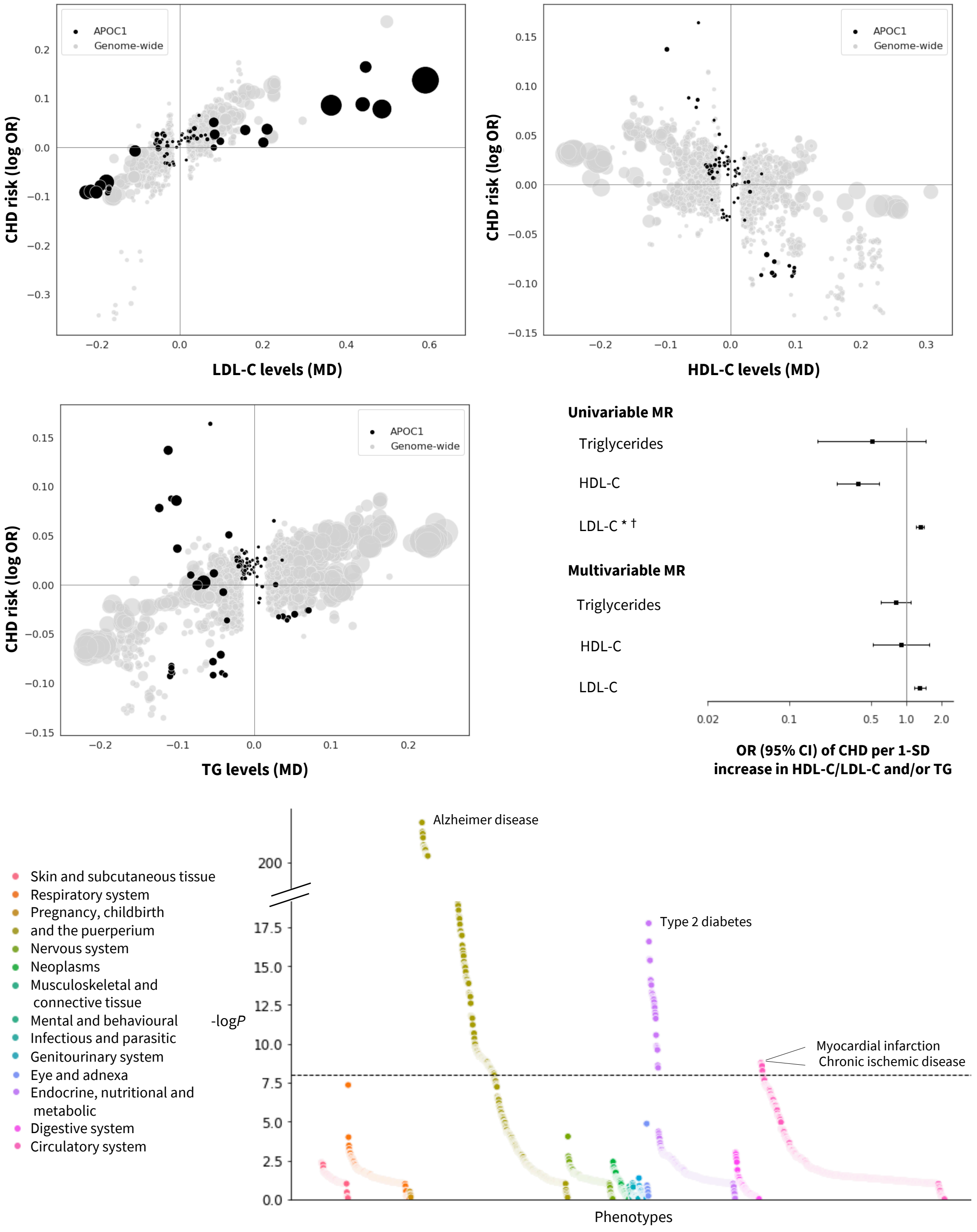


**Fig. S5. Prioritized target: APOC1.** The top and middle left panels show genetic associations at the locus (± 50kbp) in black vs genome-wide associations (grey, *P* value < 1x10$^{-6}$). The x-axis shows the per allele effect on the corresponding lipid expressed as mean difference (MD) from GLGC and the y-axis indicates the per allele effect on CHD expressed as log odds ratios (OR) from CardiogramPlusC4D. The marker size indicates the significance of the association with the lipid sub-fraction (*P*-value). The middle right panel shows the result of the univariable and multivariable (drug target) *cis*-MR results. An asterisk (*) indicates the MR estimates as being replicated, and a dagger (†) that the lipid effect and CHD signals are co-localized. The bottom panel shows disease associations at the locus with 103 clinical end points from UK Biobank and GWAS Consortia.

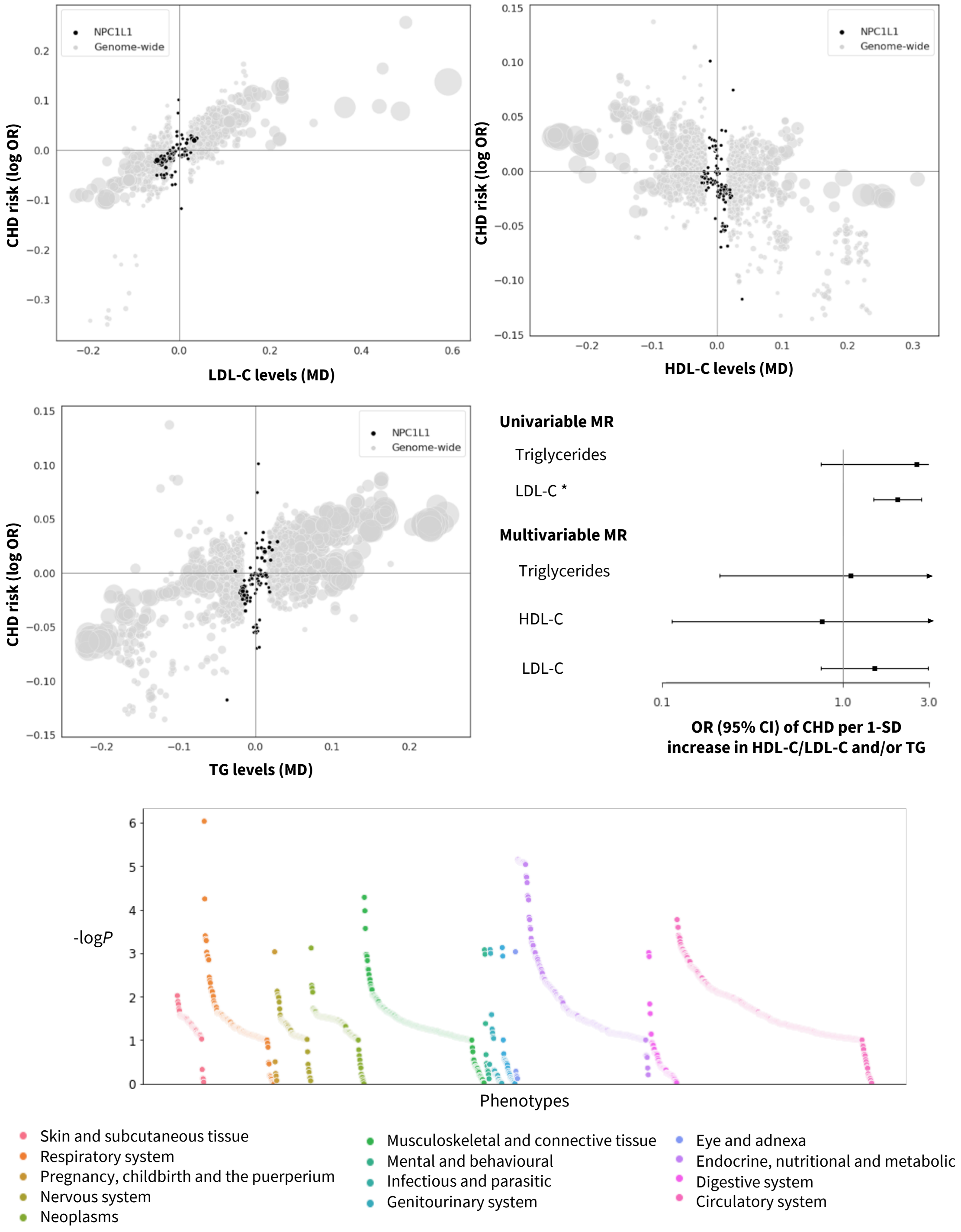


**Fig. S6. Prioritized target: NPC1L1.** The top and middle left panels show genetic associations at the locus (± 50kbp) in black vs genome-wide associations (grey, $P$ value < $1x10^{-6}$). The x-axis shows the per allele effect on the corresponding lipid expressed as mean difference (MD) from GLGC and the y-axis indicates the per allele effect on CHD expressed as log odds ratios (OR) from CardiogramPlusC4D. The marker size indicates the significance of the association with the lipid sub-fraction ($P$-value). The middle right panel shows the result of the univariable and multivariable (drug target) *cis*-MR results. An asterisk (*) indicates the MR estimates as being replicated, and a dagger (†) that the lipid effect and CHD signals are co-localized. The bottom panel shows disease associations at the locus with 103 clinical end points from UK Biobank and GWAS Consortia.

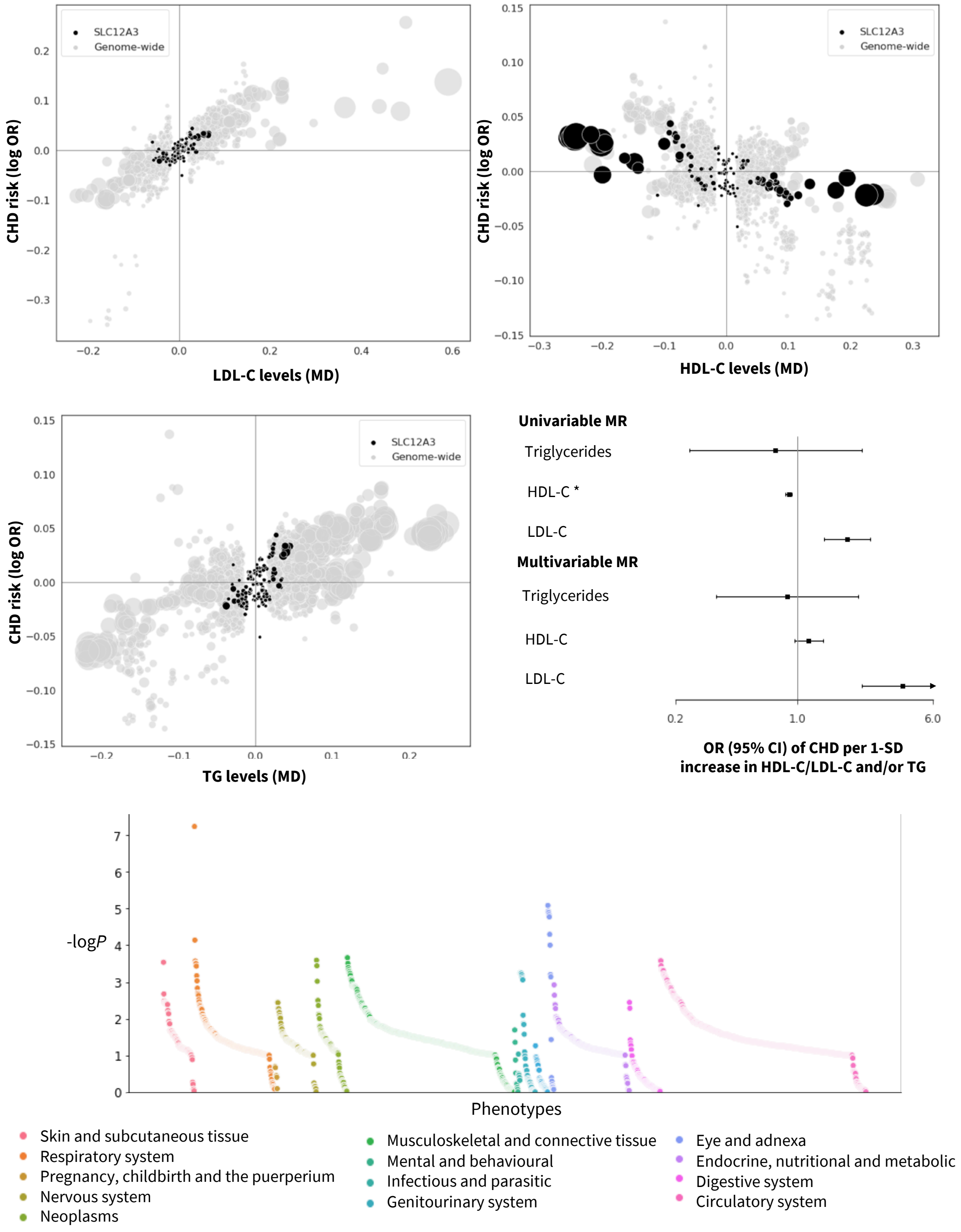


**Fig. S7. Prioritized target: SLC12A3.** The top and middle left panels show genetic associations at the locus (± 50kbp) in black vs genome-wide associations (grey, $P$ value < $1x10^{-6}$). The x-axis shows the per allele effect on the corresponding lipid expressed as mean difference (MD) from GLGC and the y-axis indicates the per allele effect on CHD expressed as log odds ratios (OR) from CardiogramPlusC4D. The marker size indicates the significance of the association with the lipid sub-fraction ($P$-value). The middle right panel shows the result of the univariable and multivariable (drug target) *cis*-MR results. An asterisk (*) indicates the MR estimates as being replicated, and a dagger (†) that the lipid effect and CHD signals are co-localized. The bottom panel shows disease associations at the locus with 103 clinical end points from UK Biobank and GWAS Consortia.

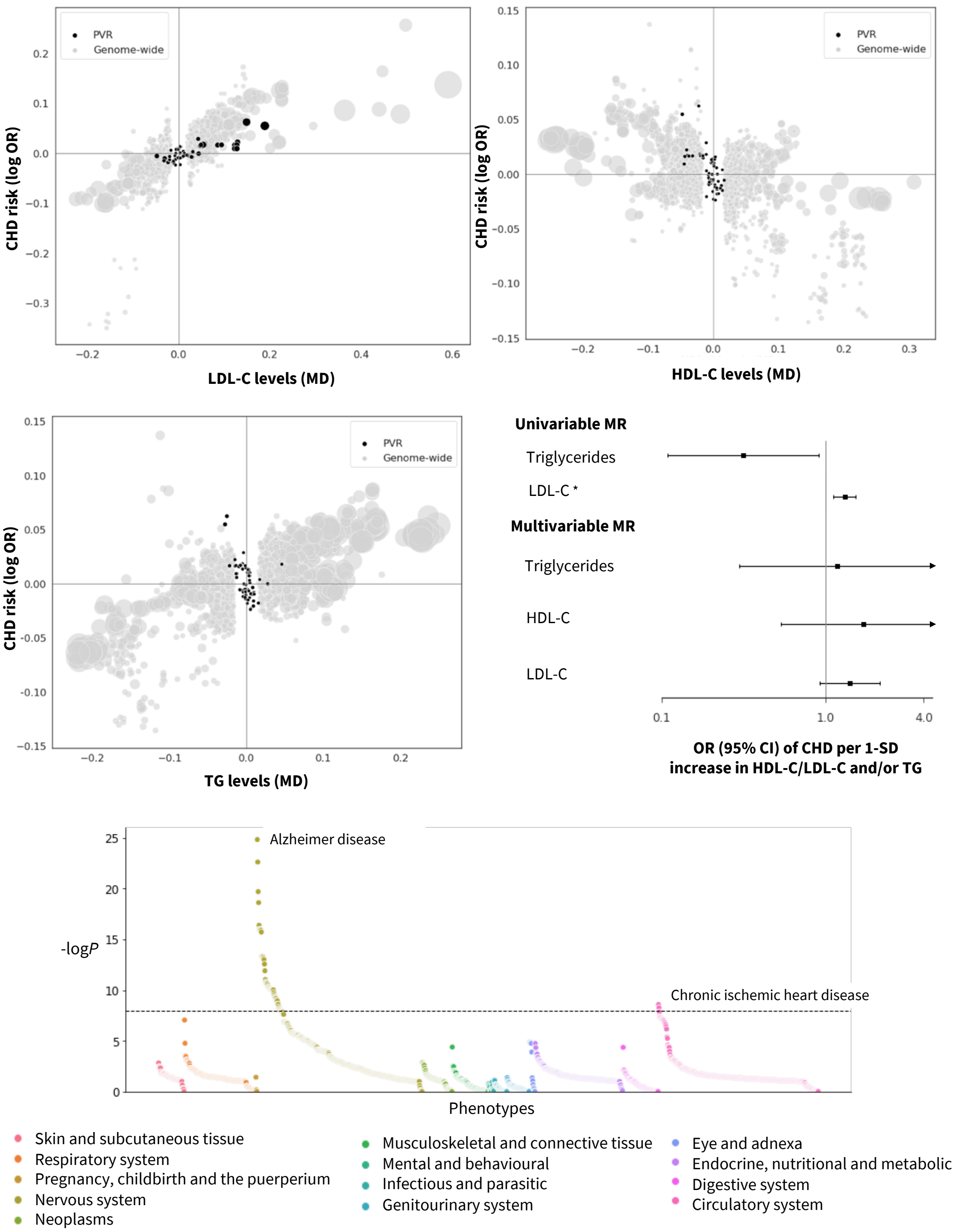


**Fig. S8. Prioritized target: PVR.** The top and middle left panels show genetic associations at the locus (± 50kbp) in black vs genome-wide associations (grey, *P* value < $1x10^{-6}$). The x-axis shows the per allele effect on the corresponding lipid expressed as mean difference (MD) from GLGC and the y-axis indicates the per allele effect on CHD expressed as log odds ratios (OR) from CardiogramPlusC4D. The marker size indicates the significance of the association with the lipid sub-fraction (*P*-value). The middle right panel shows the result of the univariable and multivariable (drug target) *cis*-MR results. An asterisk (*) indicates the MR estimates as being replicated, and a dagger (†) that the lipid effect and CHD signals are co-localized. The bottom panel shows disease associations at the locus with 103 clinical end points from UK Biobank and GWAS Consortia.

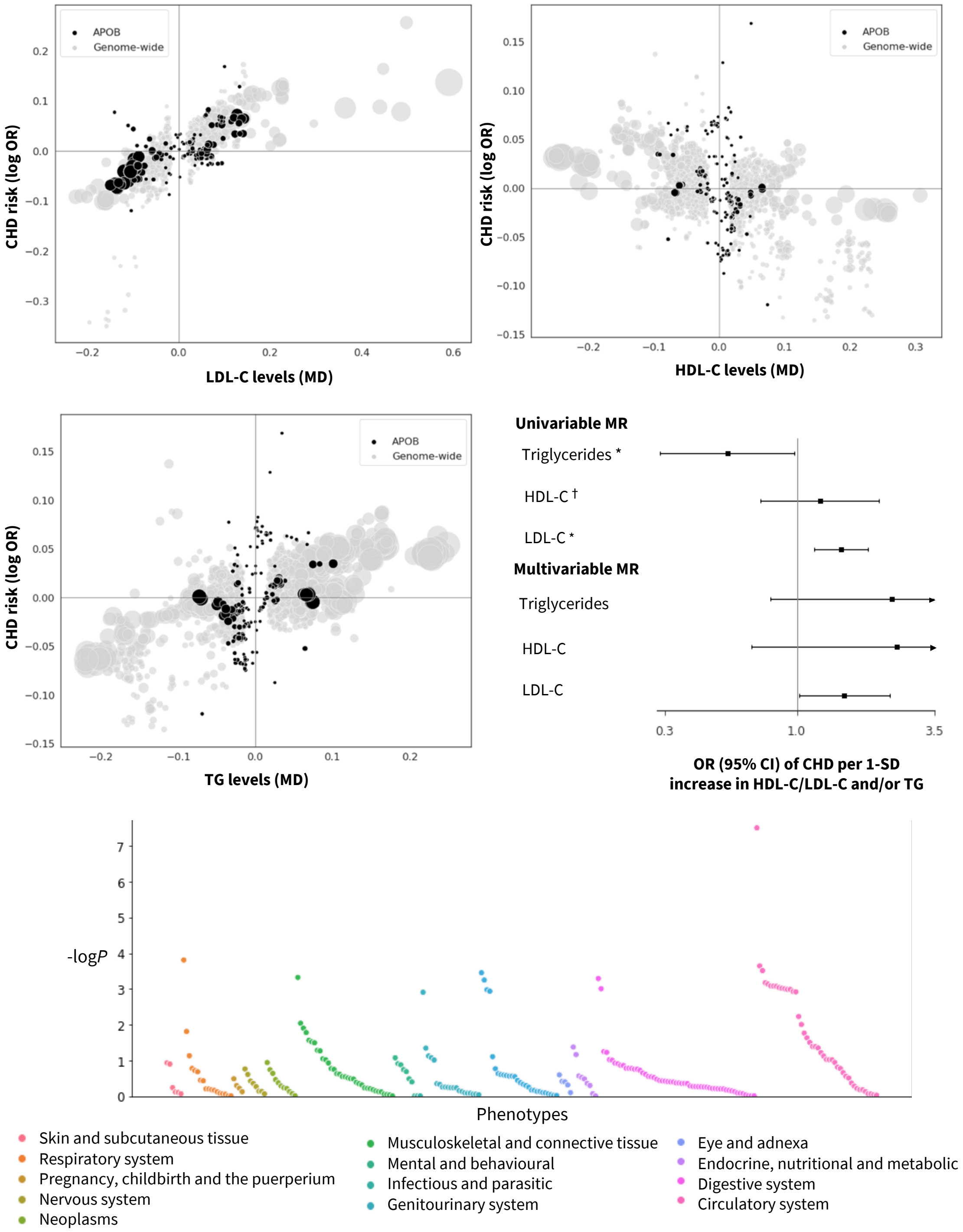


**Fig. S9. Prioritized target: APOB.** The top and middle left panels show genetic associations at the locus (± 50kbp) in black vs genome-wide associations (grey, $P$ value < $1x10^{-6}$). The x-axis shows the per allele effect on the corresponding lipid expressed as mean difference (MD) from GLGC and the y-axis indicates the per allele effect on CHD expressed as log odds ratios (OR) from CardiogramPlusC4D. The marker size indicates the significance of the association with the lipid sub-fraction ($P$-value). The middle right panel shows the result of the univariable and multivariable (drug target) *cis*-MR results. An asterisk (*) indicates the MR estimates as being replicated, and a dagger (†) that the lipid effect and CHD signals are co-localized. The bottom panel shows disease associations at the locus with 103 clinical end points from UK Biobank and GWAS Consortia.

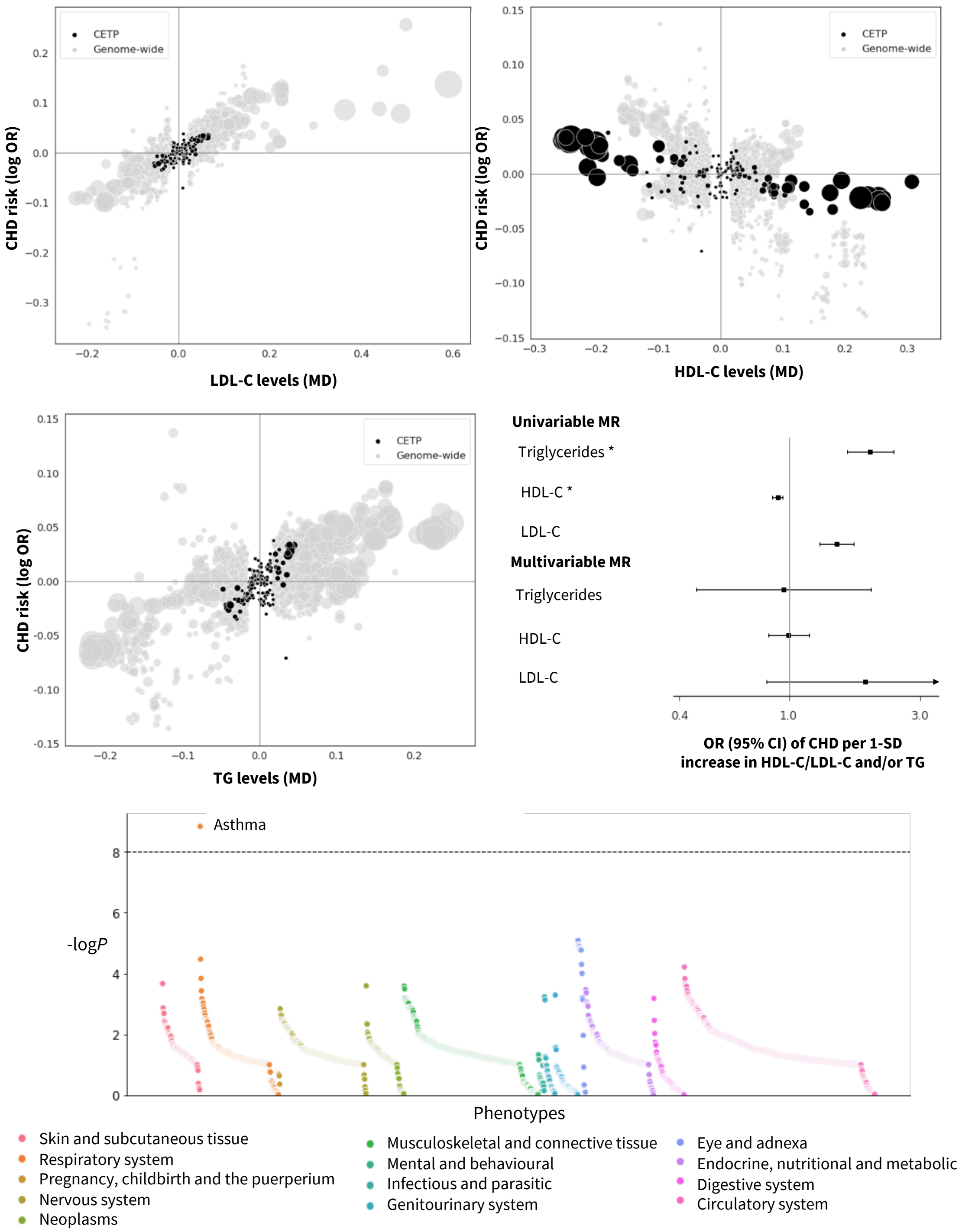


**Fig. S10. Prioritized target: CETP.** The top and middle left panels show genetic associations at the locus (± 50kbp) in black vs genome-wide associations (grey, $P$ value < $1x10^{-6}$). The x-axis shows the per allele effect on the corresponding lipid expressed as mean difference (MD) from GLGC and the y-axis indicates the per allele effect on CHD expressed as log odds ratios (OR) from CardiogramPlusC4D. The marker size indicates the significance of the association with the lipid sub-fraction ($P$-value). The middle right panel shows the result of the univariable and multivariable (drug target) *cis*-MR results. An asterisk (*) indicates the MR estimates as being replicated, and a dagger (†) that the lipid effect and CHD signals are co-localized. The bottom panel shows disease associations at the locus with 103 clinical end points from UK Biobank and GWAS Consortia.

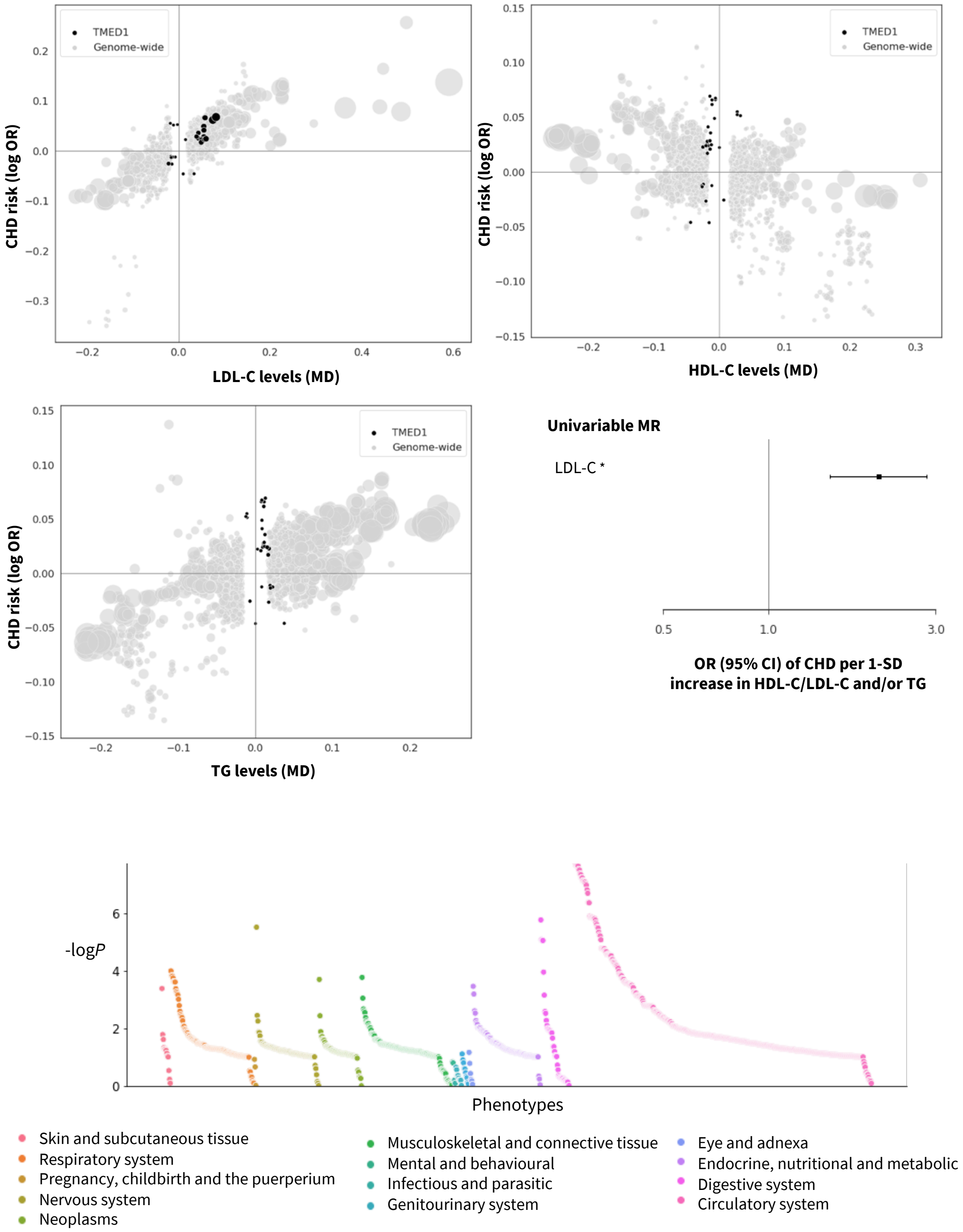


**Fig. S11. Prioritized target: TMED1.** The top and middle left panels show genetic associations at the locus (± 50kbp) in black vs genome-wide associations (grey, *P* value < $1x10^{-6}$). The x-axis shows the per allele effect on the corresponding lipid expressed as mean difference (MD) from GLGC and the y-axis indicates the per allele effect on CHD expressed as log odds ratios (OR) from CardiogramPlusC4D. The marker size indicates the significance of the association with the lipid sub-fraction (*P*-value). The middle right panel shows the result of the univariable and multivariable (drug target) *cis*-MR results. An asterisk (*) indicates the MR estimates as being replicated, and a dagger (†) that the lipid effect and CHD signals are co-localized. The bottom panel shows disease associations at the locus with 103 clinical end points from UK Biobank and GWAS Consortia.

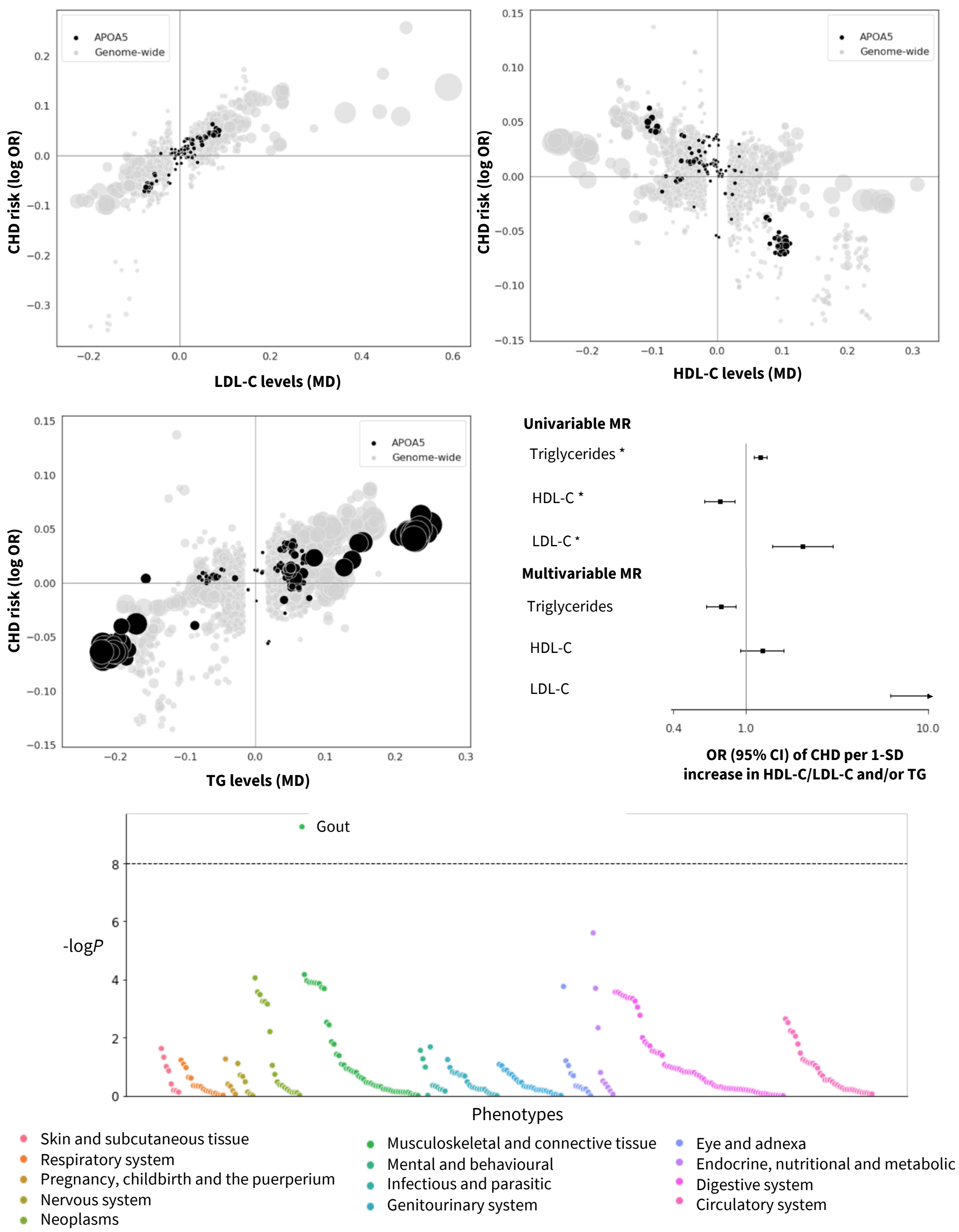


**Fig. S12. Prioritized target: APOA5.** The top and middle left panels show genetic associations at the locus (± 50kbp) in black vs genome-wide associations (grey, *P* value < $1x10^{-6}$). The x-axis shows the per allele effect on the corresponding lipid expressed as mean difference (MD) from GLGC and the y-axis indicates the per allele effect on CHD expressed as log odds ratios (OR) from CardiogramPlusC4D. The marker size indicates the significance of the association with the lipid sub-fraction (*P*-value). The middle right panel shows the result of the univariable and multivariable (drug target) *cis*-MR results. An asterisk (*) indicates the MR estimates as being replicated, and a dagger (†) that the lipid effect and CHD signals are co-localized. The bottom panel shows disease associations at the locus with 103 clinical end points from UK Biobank and GWAS Consortia.

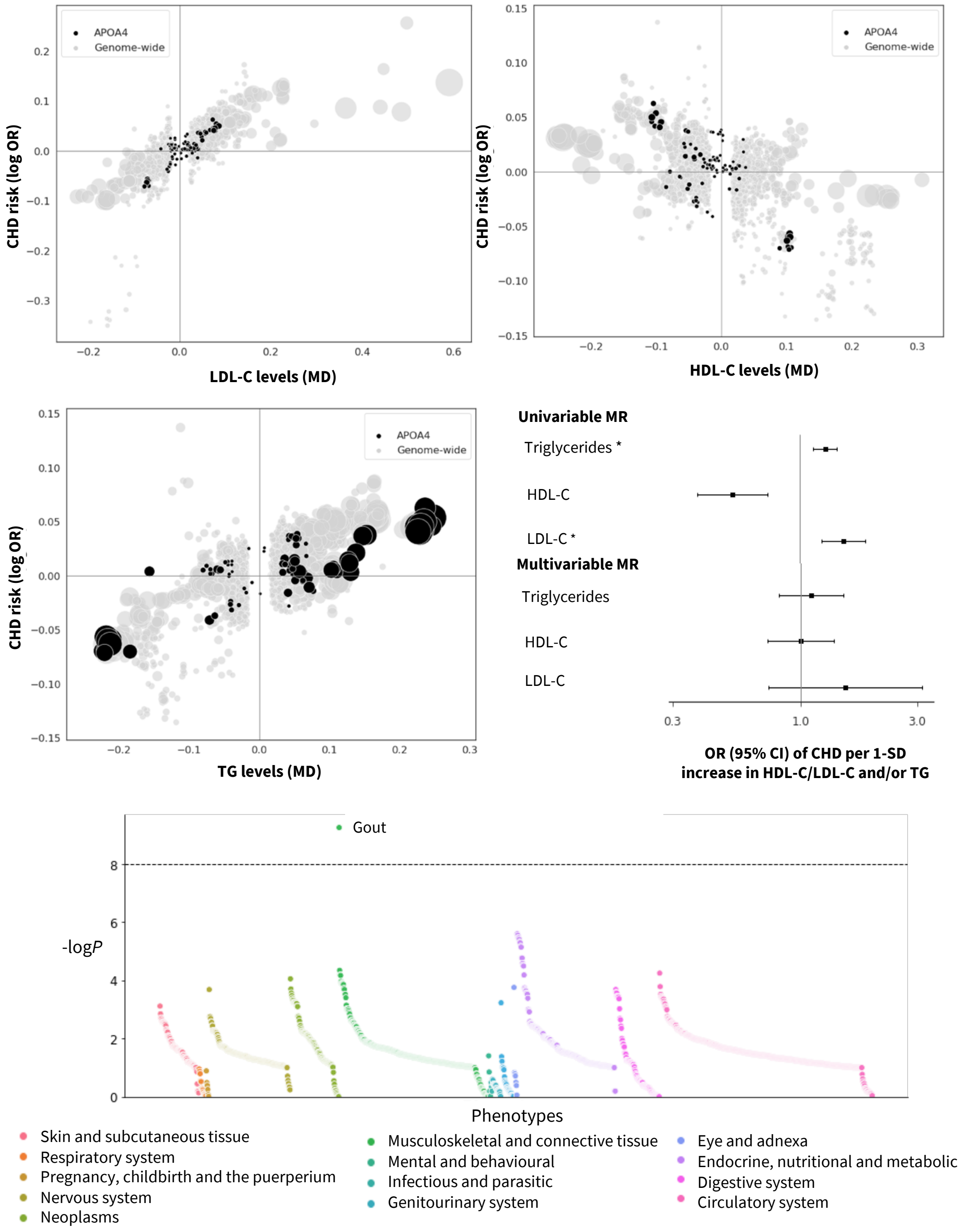


**Fig. S13. Prioritized target: APOA4.** The top and middle left panels show genetic associations at the locus (± 50kbp) in black vs genome-wide associations (grey, *P* value < 1x10$^{-6}$). The x-axis shows the per allele effect on the corresponding lipid expressed as mean difference (MD) from GLGC and the y-axis indicates the per allele effect on CHD expressed as log odds ratios (OR) from CardiogramPlusC4D. The marker size indicates the significance of the association with the lipid sub-fraction (*P*-value). The middle right panel shows the result of the univariable and multivariable (drug target) *cis*-MR results. An asterisk (*) indicates the MR estimates as being replicated, and a dagger (†) that the lipid effect and CHD signals are co-localized. The bottom panel shows disease associations at the locus with 103 clinical end points from UK Biobank and GWAS Consortia.

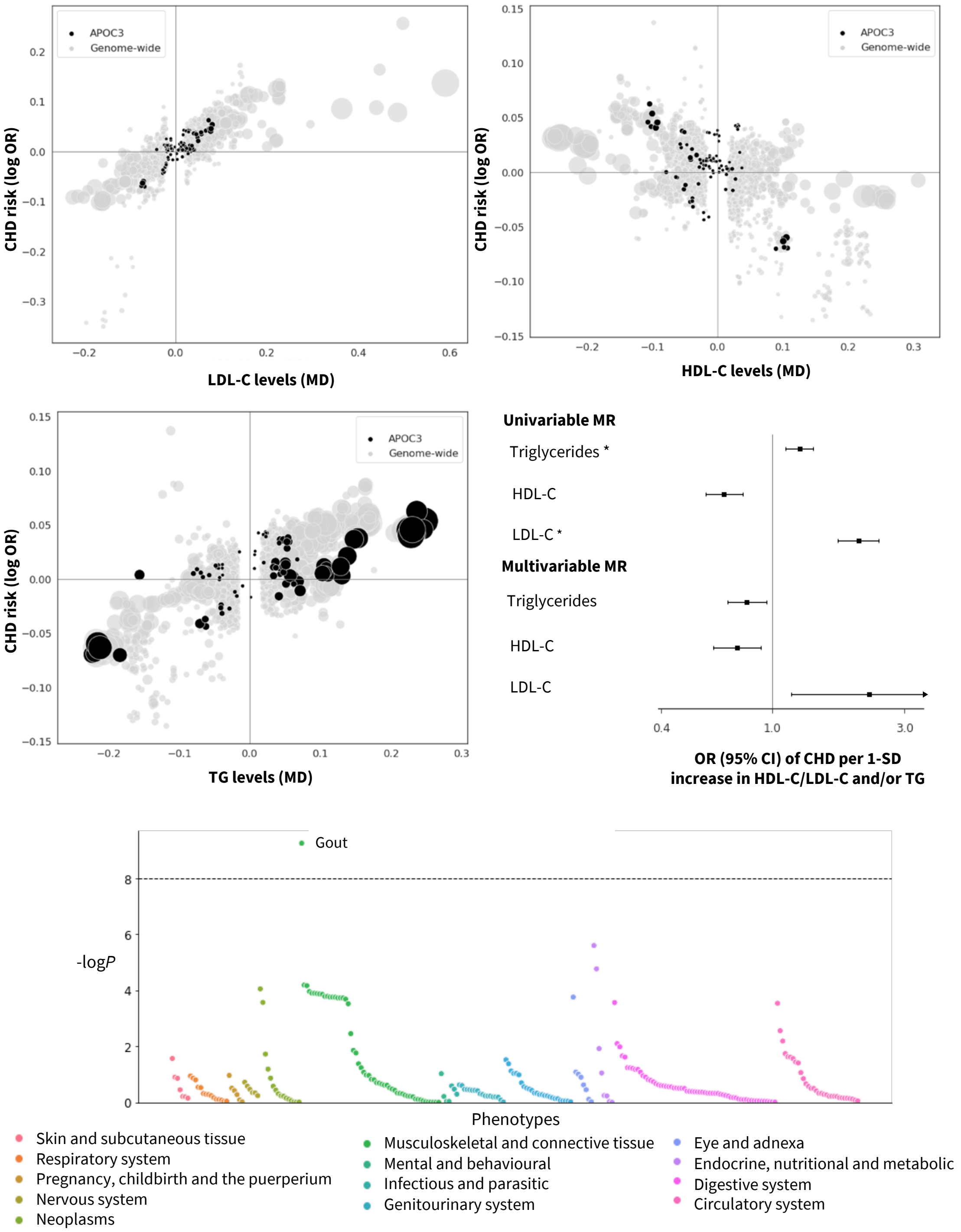


**Fig. S14. Prioritized target: APOC3.** The top and middle left panels show genetic associations at the locus (± 50kbp) in black vs genome-wide associations (grey, $P$ value < $1\text{x}10^{-6}$). The x-axis shows the per allele effect on the corresponding lipid expressed as mean difference (MD) from GLGC and the y-axis indicates the per allele effect on CHD expressed as log odds ratios (OR) from CardiogramPlusC4D. The marker size indicates the significance of the association with the lipid sub-fraction ($P$-value). The middle right panel shows the result of the univariable and multivariable (drug target) *cis*-MR results. An asterisk (*) indicates the MR estimates as being replicated, and a dagger (†) that the lipid effect and CHD signals are co-localized. The bottom panel shows disease associations at the locus with 103 clinical end points from UK Biobank and GWAS Consortia.

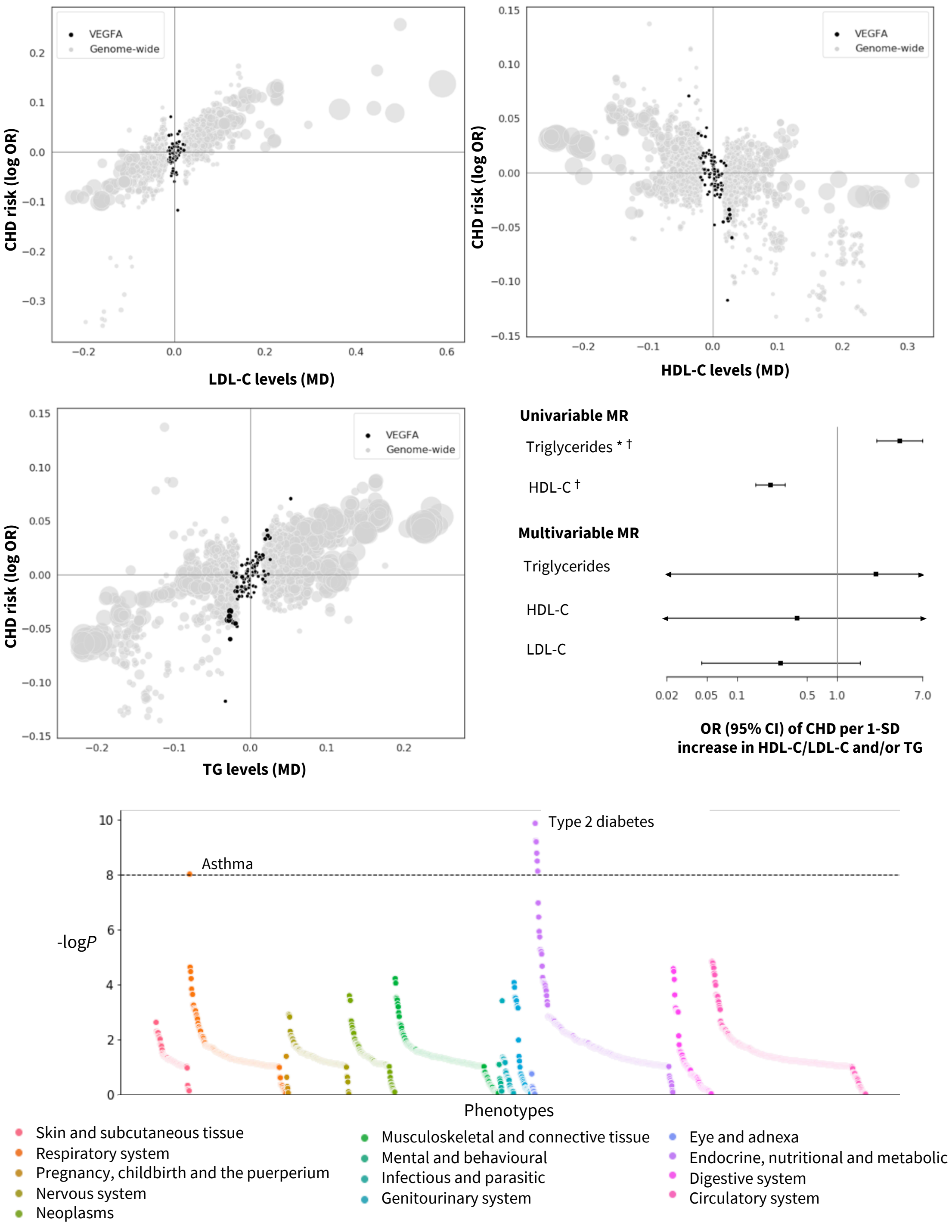


**Fig. S15. Prioritized target: VEGFA.** The top and middle left panels show genetic associations at the locus (± 50kbp) in black vs genome-wide associations (grey, $P$ value < $1x10^{-6}$). The x-axis shows the per allele effect on the corresponding lipid expressed as mean difference (MD) from GLGC and the y-axis indicates the per allele effect on CHD expressed as log odds ratios (OR) from CardiogramPlusC4D. The marker size indicates the significance of the association with the lipid sub-fraction ($P$-value). The middle right panel shows the result of the univariable and multivariable (drug target) *cis*-MR results. An asterisk (*) indicates the MR estimates as being replicated, and a dagger (†) that the lipid effect and CHD signals are co-localized. The bottom panel shows disease associations at the locus with 103 clinical end points from UK Biobank and GWAS Consortia.

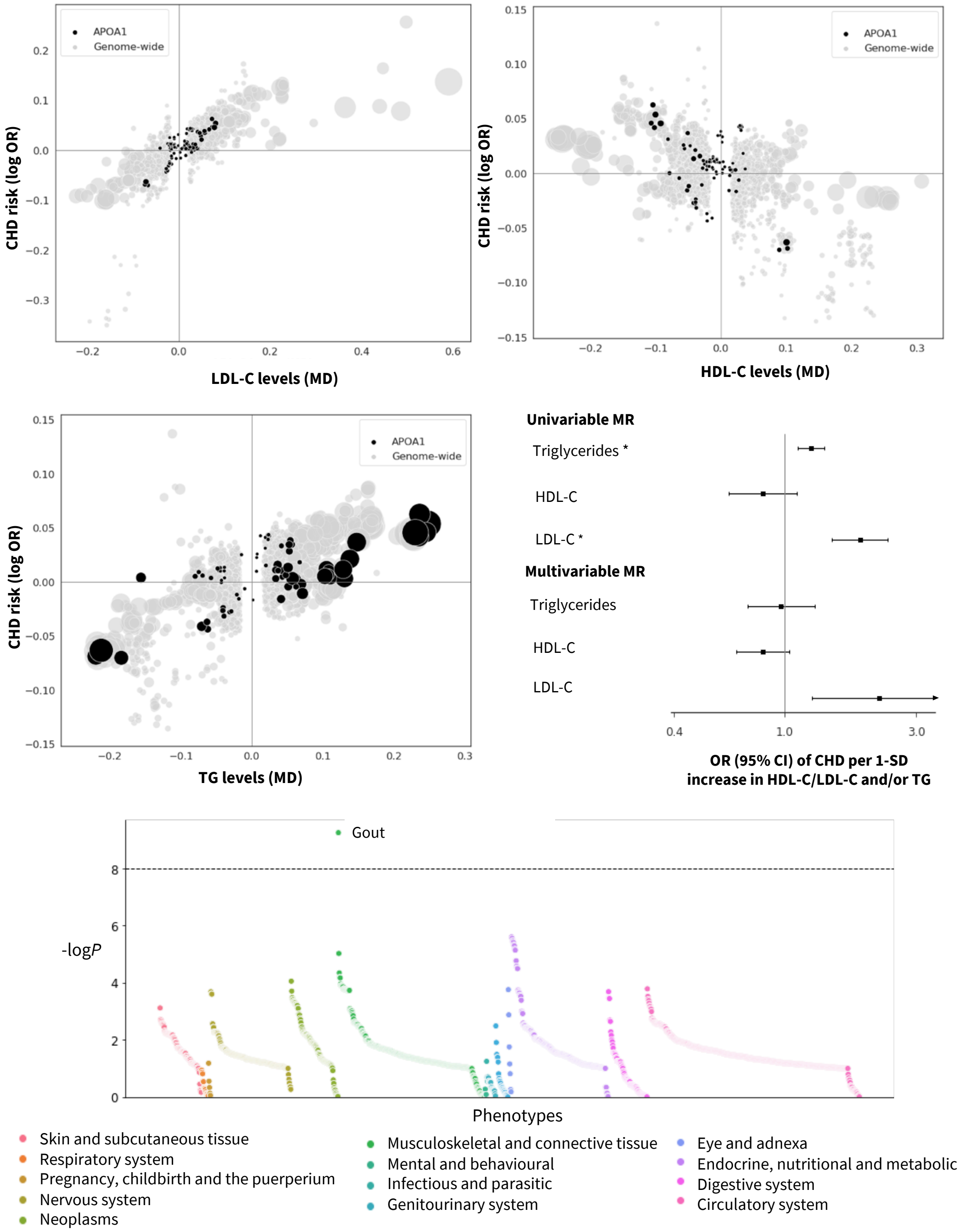


**Fig. S16. Prioritized target: APOA1.** The top and middle left panels show genetic associations at the locus (± 50kbp) in black vs genome-wide associations (grey, *P* value < $1x10^{-6}$). The x-axis shows the per allele effect on the corresponding lipid expressed as mean difference (MD) from GLGC and the y-axis indicates the per allele effect on CHD expressed as log odds ratios (OR) from CardiogramPlusC4D. The marker size indicates the significance of the association with the lipid sub-fraction (*P*-value). The middle right panel shows the result of the univariable and multivariable (drug target) *cis*-MR results. An asterisk (*) indicates the MR estimates as being replicated, and a dagger (†) that the lipid effect and CHD signals are co-localized. The bottom panel shows disease associations at the locus with 103 clinical end points from UK Biobank and GWAS Consortia.

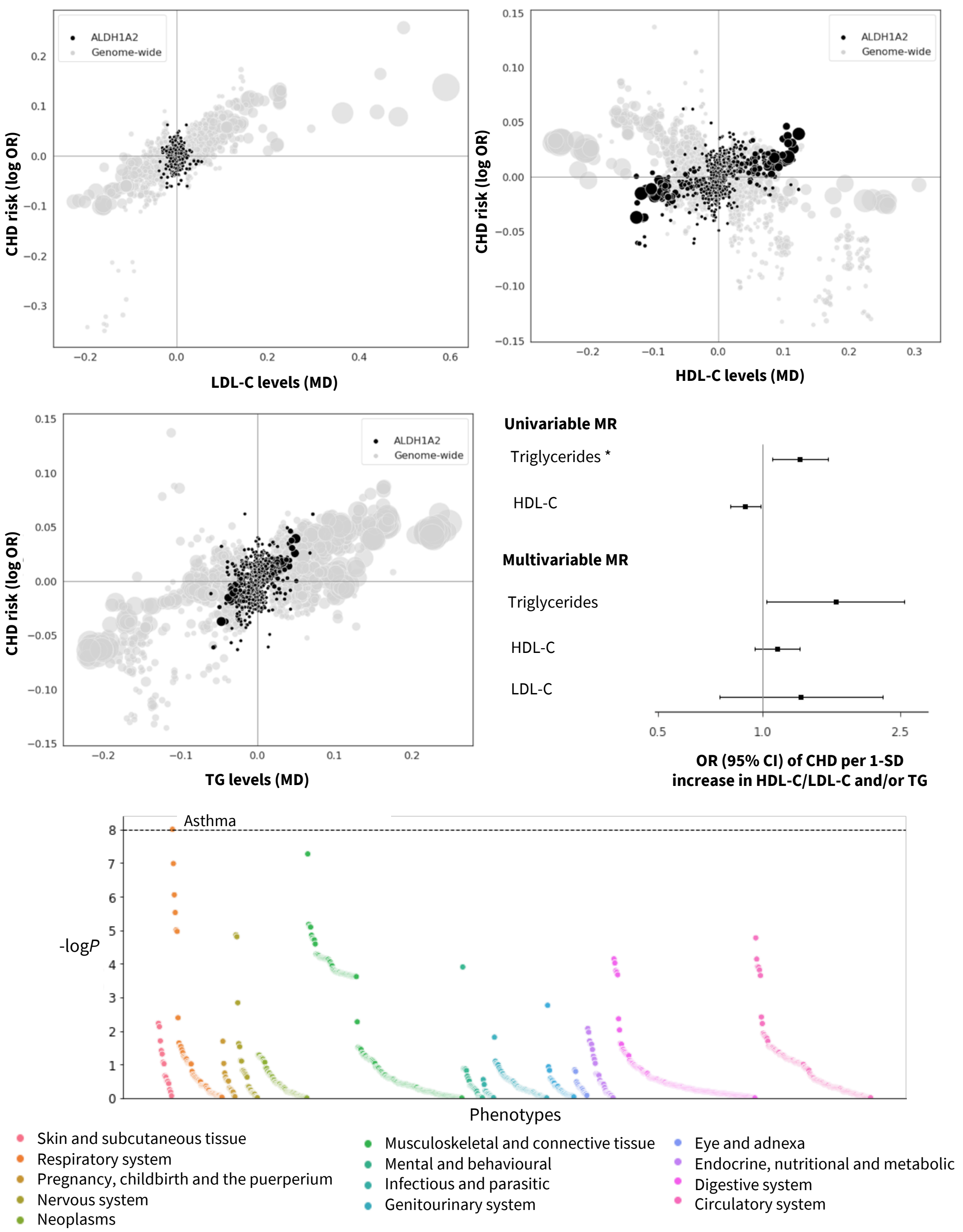


**Fig. S17. Prioritized target: ALDH1A2.** The top and middle left panels show genetic associations at the locus (± 50kbp) in black vs genome-wide associations (grey, *P* value < $1x10^{-6}$). The x-axis shows the per allele effect on the corresponding lipid expressed as mean difference (MD) from GLGC and the y-axis indicates the per allele effect on CHD expressed as log odds ratios (OR) from CardiogramPlusC4D. The marker size indicates the significance of the association with the lipid sub-fraction (*P*-value). The middle right panel shows the result of the univariable and multivariable (drug target) *cis*-MR results. An asterisk (*) indicates the MR estimates as being replicated, and a dagger (†) that the lipid effect and CHD signals are co-localized. The bottom panel shows disease associations at the locus with 103 clinical end points from UK Biobank and GWAS Consortia.

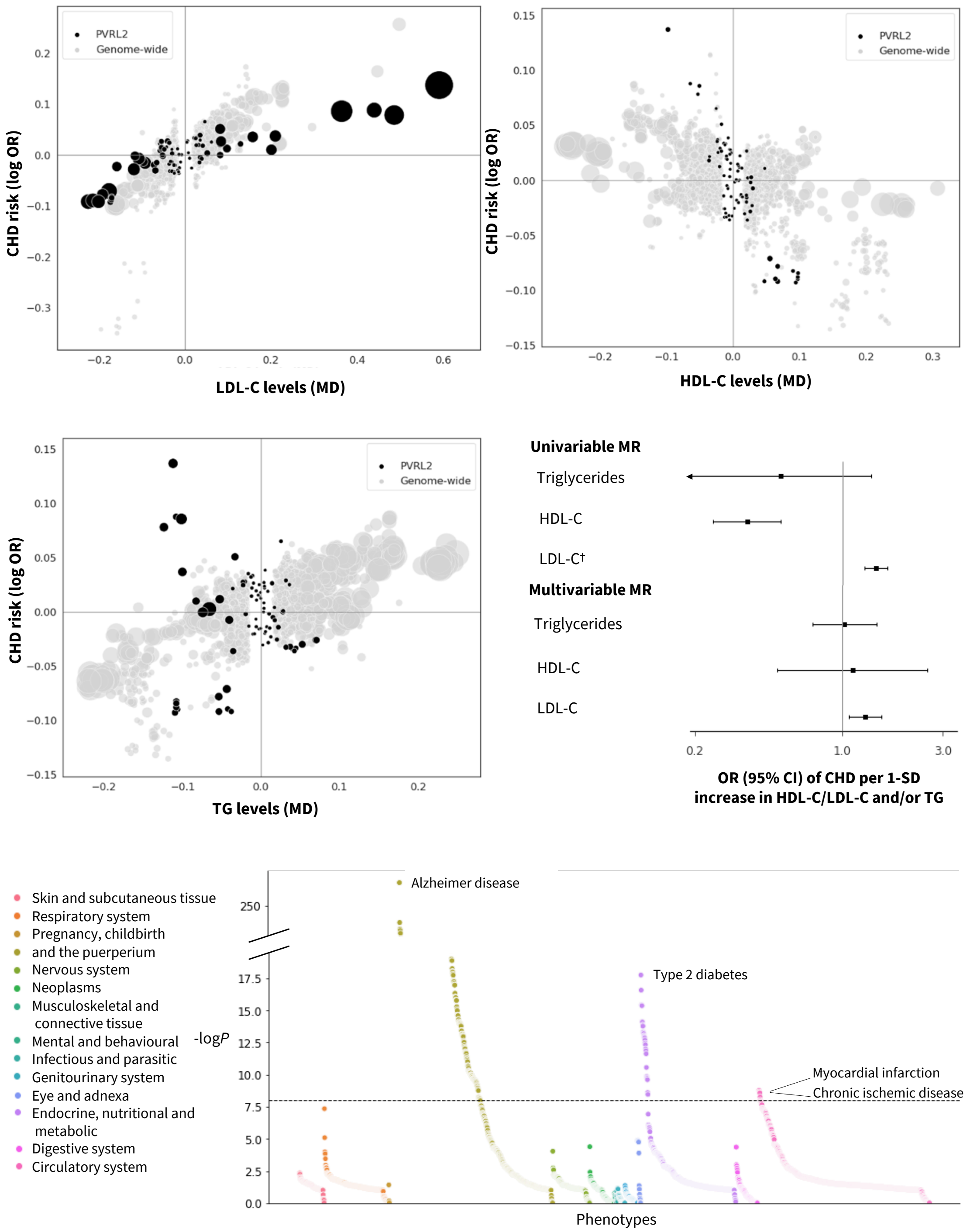


**Fig. S18. Prioritized target: PVRL2.** The top and middle left panels show genetic associations at the locus (± 50kbp) in black vs genome-wide associations (grey, *P* value < 1x10$^{-6}$). The x-axis shows the per allele effect on the corresponding lipid expressed as mean difference (MD) from GLGC and the y-axis indicates the per allele effect on CHD expressed as log odds ratios (OR) from CardiogramPlusC4D. The marker size indicates the significance of the association with the lipid sub-fraction (*P*-value). The middle right panel shows the result of the univariable and multivariable (drug target) *cis*-MR results. An asterisk (*) indicates the MR estimates as being replicated, and a dagger (†) that the lipid effect and CHD signals are co-localized. The bottom panel shows disease associations at the locus with 103 clinical end points from UK Biobank and GWAS Consortia.

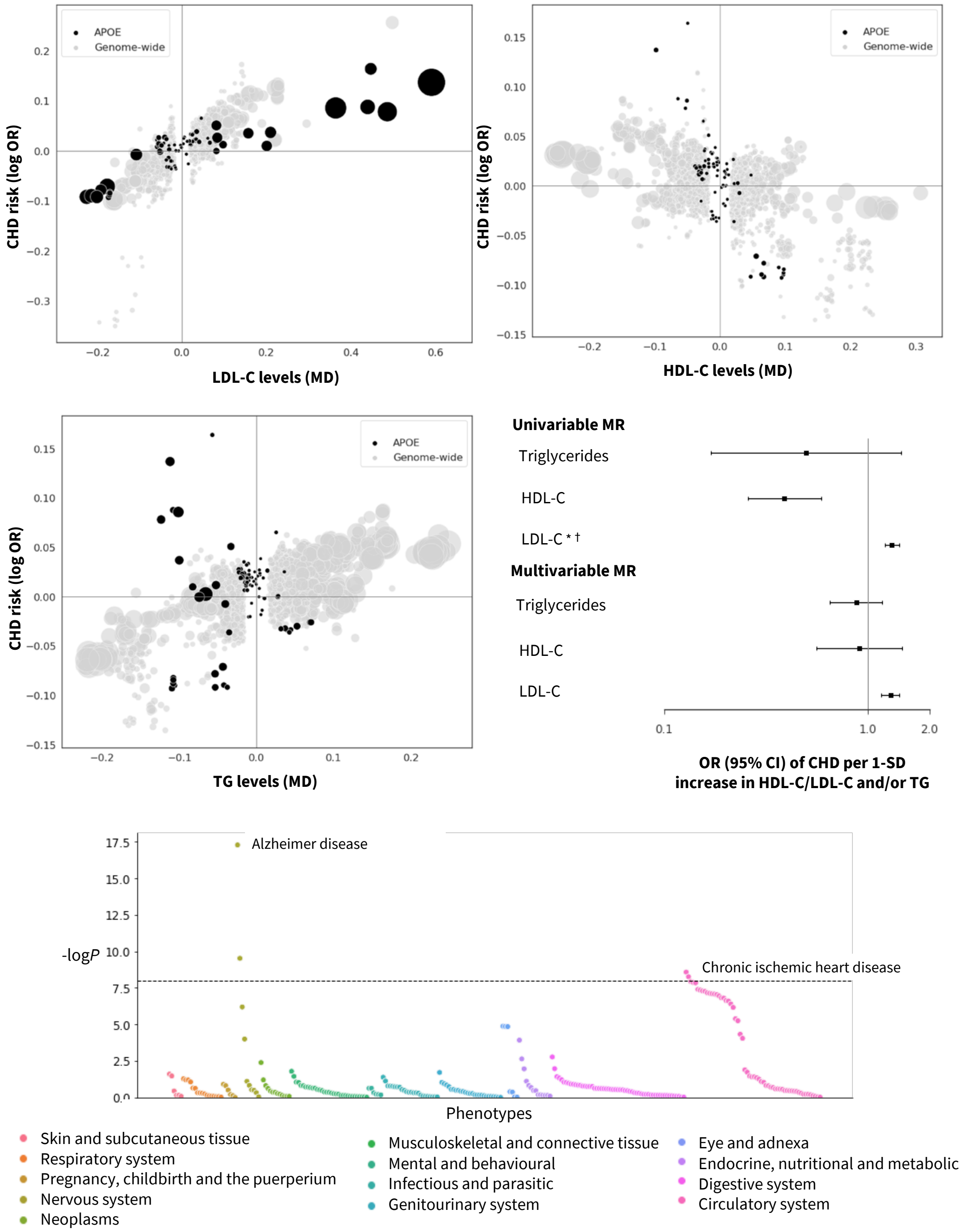


**Fig. S19. Prioritized target: APOE.** The top and middle left panels show genetic associations at the locus (± 50kbp) in black vs genome-wide associations (grey, $P$ value < $1 \times 10^{-6}$). The x-axis shows the per allele effect on the corresponding lipid expressed as mean difference (MD) from GLGC and the y-axis indicates the per allele effect on CHD expressed as log odds ratios (OR) from CardiogramPlusC4D. The marker size indicates the significance of the association with the lipid sub-fraction ($P$-value). The middle right panel shows the result of the univariable and multivariable (drug target) *cis*-MR results. An asterisk (*) indicates the MR estimates as being replicated, and a dagger (†) that the lipid effect and CHD signals are co-localized. The bottom panel shows disease associations at the locus with 103 clinical end points from UK Biobank and GWAS Consortia.

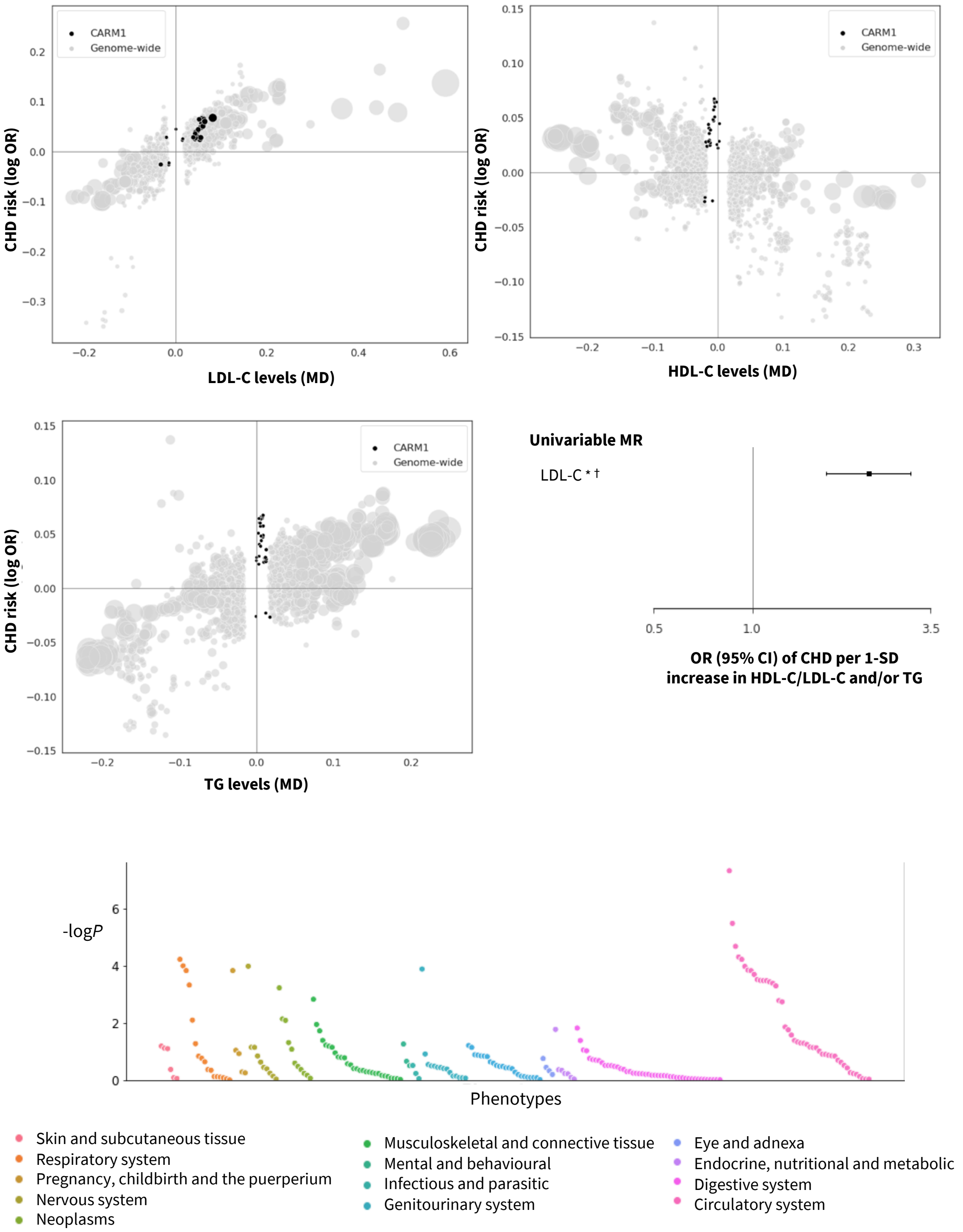


**Fig. S20. Prioritized target: CARM1.** The top and middle left panels show genetic associations at the locus (± 50kbp) in black vs genome-wide associations (grey, $P$ value < $1x10^{-6}$). The x-axis shows the per allele effect on the corresponding lipid expressed as mean difference (MD) from GLGC and the y-axis indicates the per allele effect on CHD expressed as log odds ratios (OR) from CardiogramPlusC4D. The marker size indicates the significance of the association with the lipid sub-fraction ($P$-value). The middle right panel shows the result of the univariable and multivariable (drug target) *cis*-MR results. An asterisk (*) indicates the MR estimates as being replicated, and a dagger (†) that the lipid effect and CHD signals are co-localized. The bottom panel shows disease associations at the locus with 103 clinical end points from UK Biobank and GWAS Consortia.

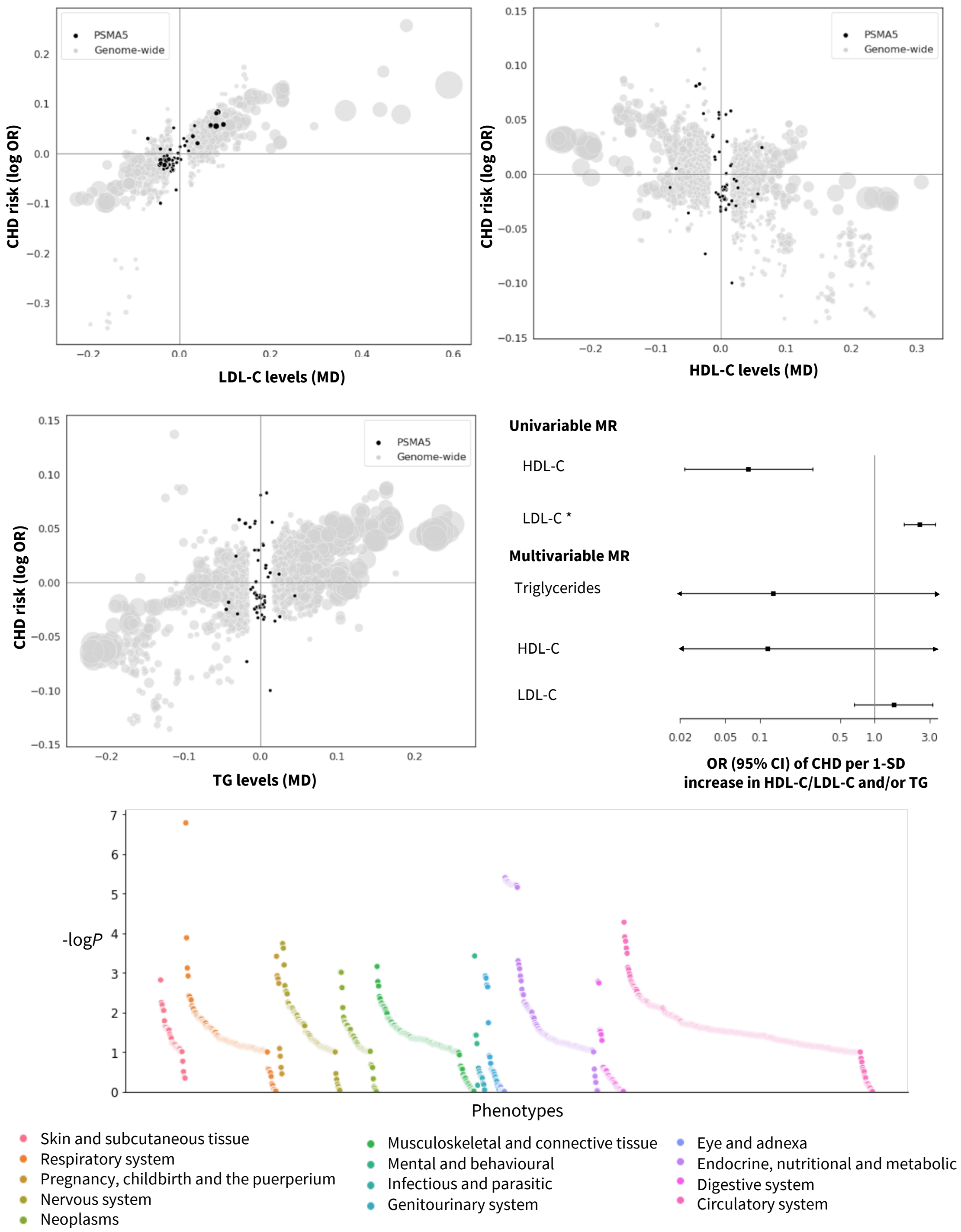


**Fig. S21. Prioritized target: PSMA5.** The top and middle left panels show genetic associations at the locus (± 50kbp) in black vs genome-wide associations (grey, $P$ value < $1x10^{-6}$). The x-axis shows the per allele effect on the corresponding lipid expressed as mean difference (MD) from GLGC and the y-axis indicates the per allele effect on CHD expressed as log odds ratios (OR) from CardiogramPlusC4D. The marker size indicates the significance of the association with the lipid sub-fraction ($P$-value). The middle right panel shows the result of the univariable and multivariable (drug target) *cis*-MR results. An asterisk (*) indicates the MR estimates as being replicated, and a dagger (†) that the lipid effect and CHD signals are co-localized. The bottom panel shows disease associations at the locus with 103 clinical end points from UK Biobank and GWAS Consortia.

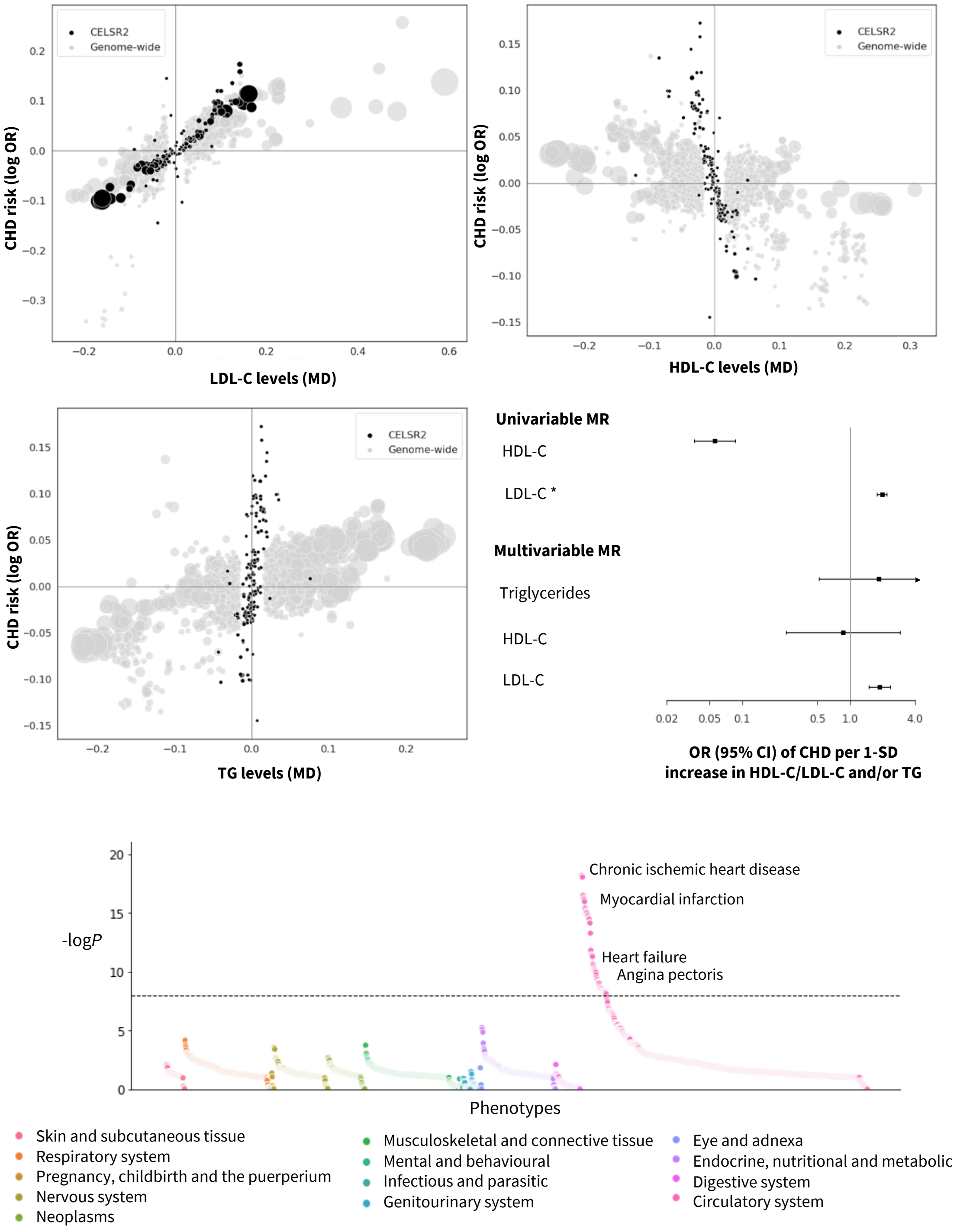


**Fig. S22. Prioritized target: CELSR2.** The top and middle left panels show genetic associations at the locus (± 50kbp) in black vs genome-wide associations (grey, *P* value < $1 \times 10^{-6}$). The x-axis shows the per allele effect on the corresponding lipid expressed as mean difference (MD) from GLGC and the y-axis indicates the per allele effect on CHD expressed as log odds ratios (OR) from CardiogramPlusC4D. The marker size indicates the significance of the association with the lipid sub-fraction (*P*-value). The middle right panel shows the result of the univariable and multivariable (drug target) *cis*-MR results. An asterisk (*) indicates the MR estimates as being replicated, and a dagger (†) that the lipid effect and CHD signals are co-localized. The bottom panel shows disease associations at the locus with 103 clinical end points from UK Biobank and GWAS Consortia.

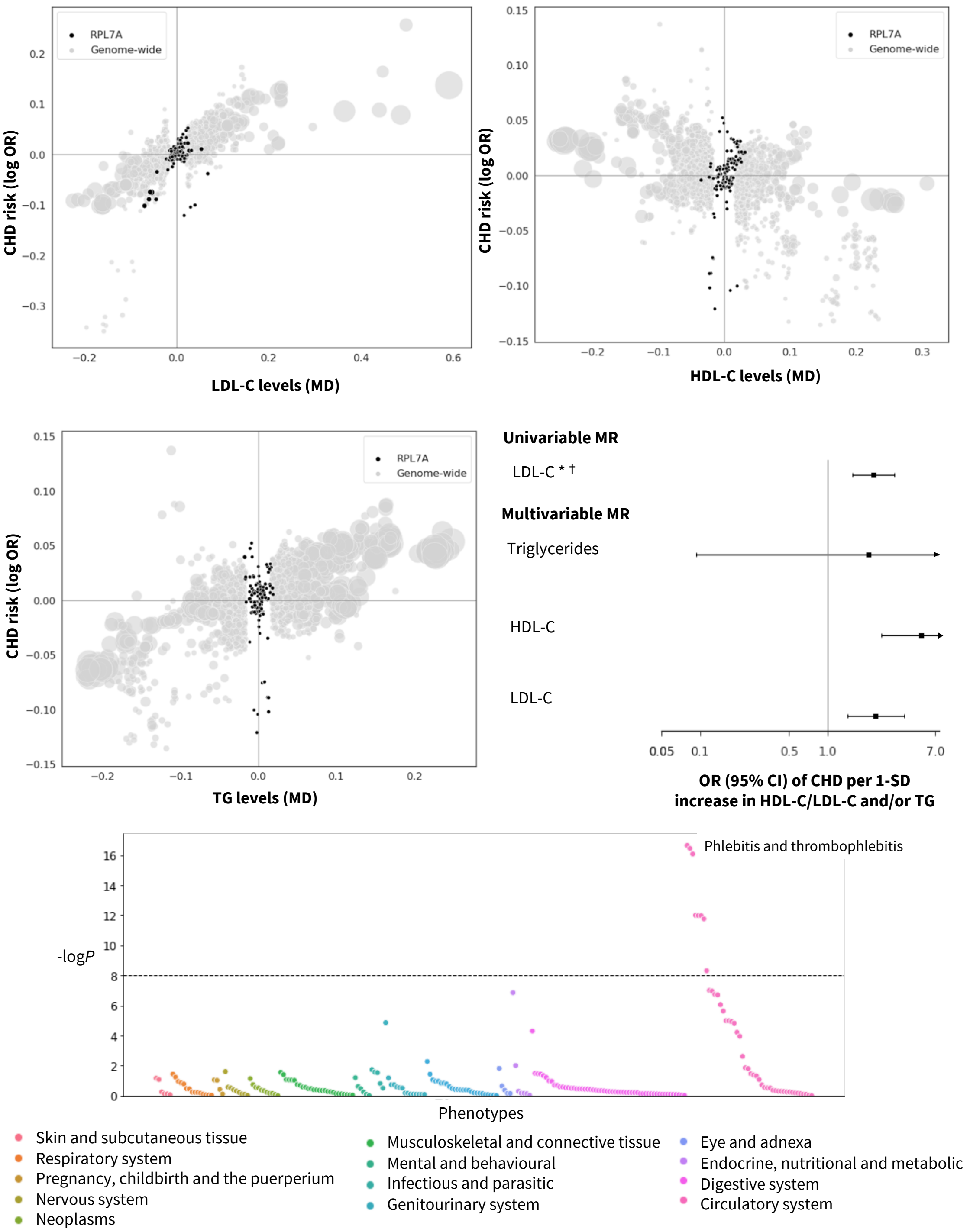


**Fig. S23. Prioritized target: RPL7A.** The top and middle left panels show genetic associations at the locus (± 50kbp) in black vs genome-wide associations (grey, $P$ value < $1x10^{-6}$). The x-axis shows the per allele effect on the corresponding lipid expressed as mean difference (MD) from GLGC and the y-axis indicates the per allele effect on CHD expressed as log odds ratios (OR) from CardiogramPlusC4D. The marker size indicates the significance of the association with the lipid sub-fraction ($P$-value). The middle right panel shows the result of the univariable and multivariable (drug target) *cis*-MR results. An asterisk (*) indicates the MR estimates as being replicated, and a dagger (†) that the lipid effect and CHD signals are co-localized. The bottom panel shows disease associations at the locus with 103 clinical end points from UK Biobank and GWAS Consortia.

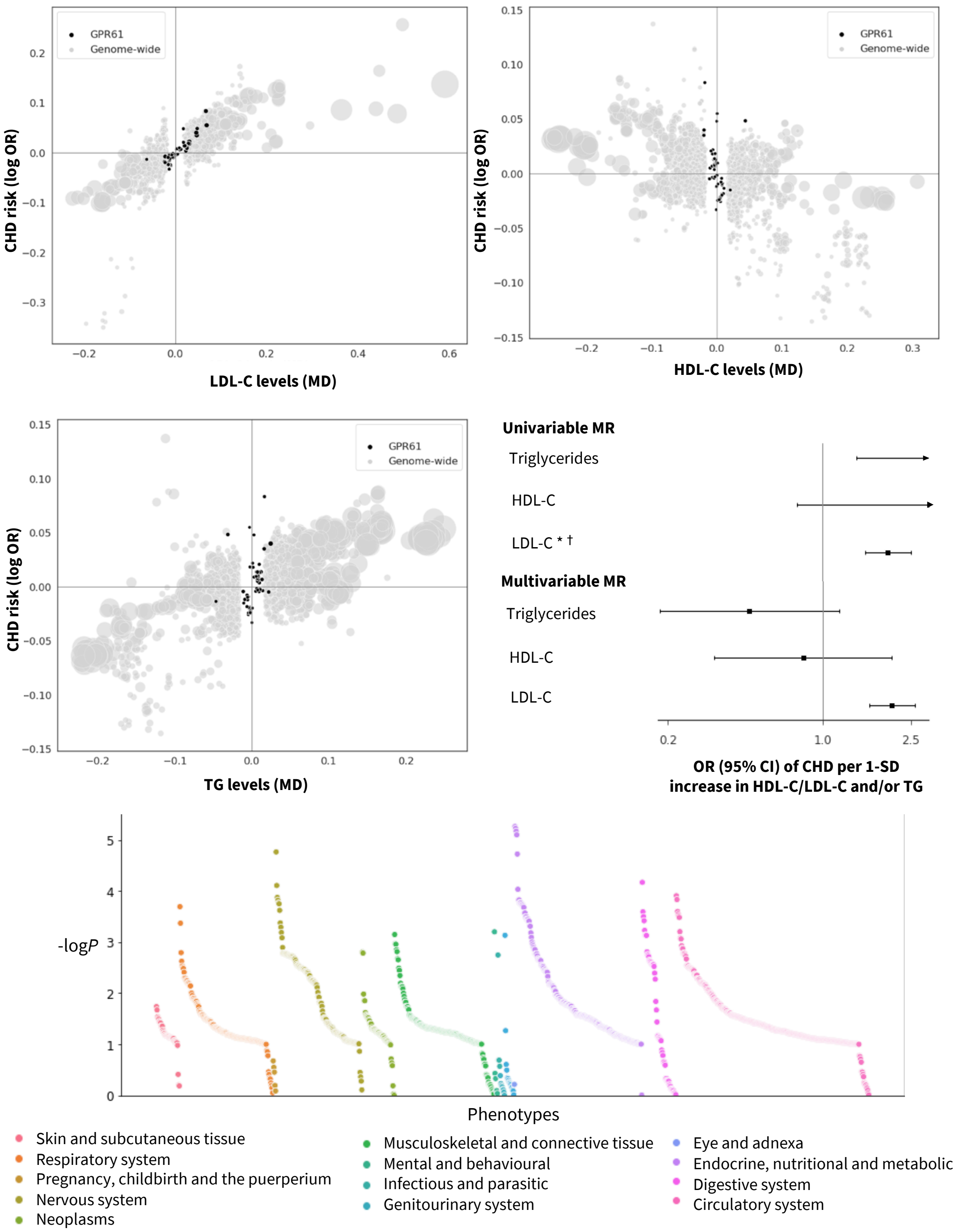


**Fig. S24. Prioritized target: GPR61.** The top and middle left panels show genetic associations at the locus (± 50kbp) in black vs genome-wide associations (grey, $P$ value < $1x10^{-6}$). The x-axis shows the per allele effect on the corresponding lipid expressed as mean difference (MD) from GLGC and the y-axis indicates the per allele effect on CHD expressed as log odds ratios (OR) from CardiogramPlusC4D. The marker size indicates the significance of the association with the lipid sub-fraction ($P$-value). The middle right panel shows the result of the univariable and multivariable (drug target) *cis*-MR results. An asterisk (*) indicates the MR estimates as being replicated, and a dagger (†) that the lipid effect and CHD signals are co-localized. The bottom panel shows disease associations at the locus with 103 clinical end points from UK Biobank and GWAS Consortia.

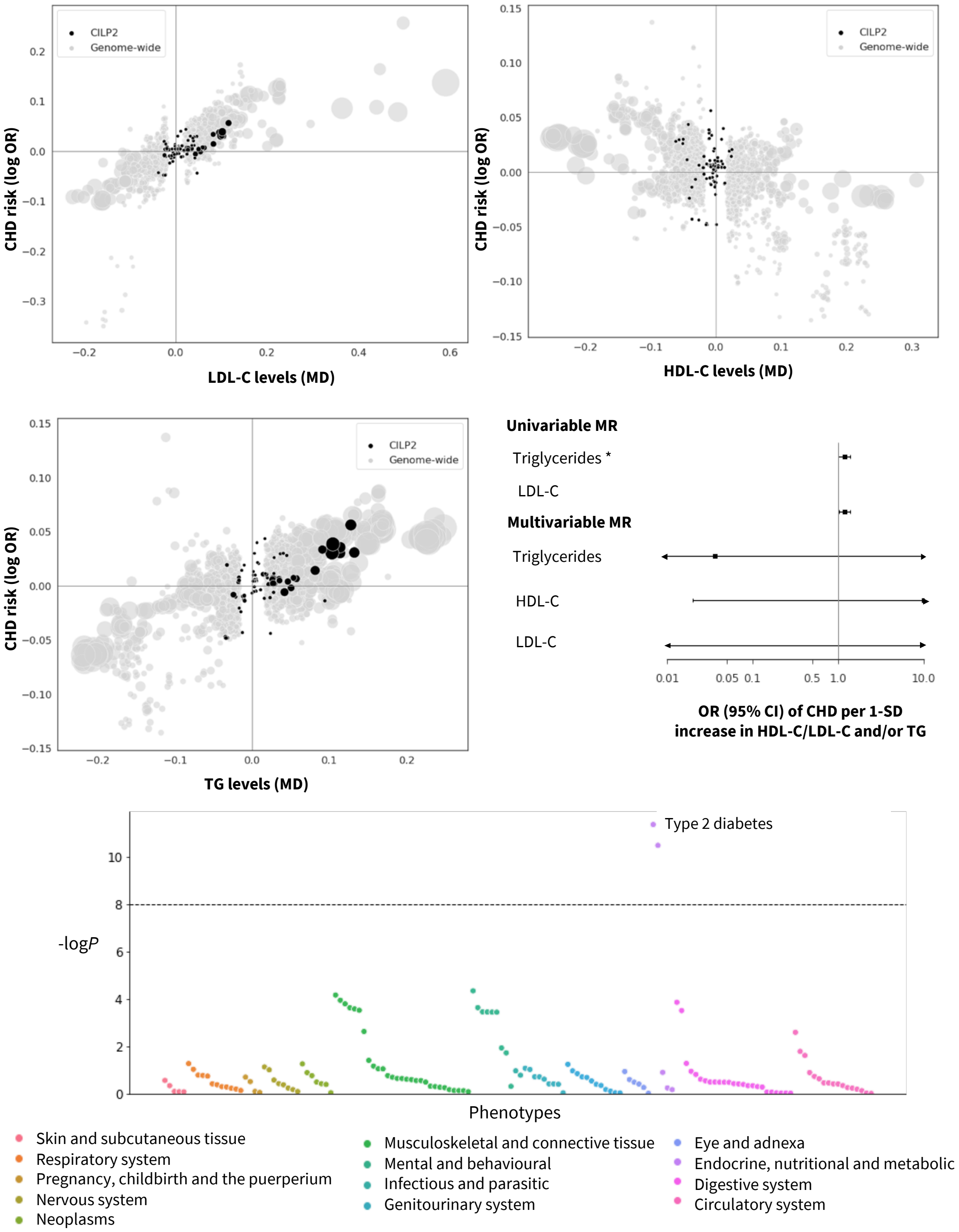


**Fig. S25. Prioritized target: CILP2.** The top and middle left panels show genetic associations at the locus (± 50kbp) in black vs genome-wide associations (grey, $P$ value < $1 \times 10^{-6}$). The x-axis shows the per allele effect on the corresponding lipid expressed as mean difference (MD) from GLGC and the y-axis indicates the per allele effect on CHD expressed as log odds ratios (OR) from CardiogramPlusC4D. The marker size indicates the significance of the association with the lipid sub-fraction ($P$-value). The middle right panel shows the result of the univariable and multivariable (drug target) *cis*-MR results. An asterisk (*) indicates the MR estimates as being replicated, and a dagger (†) that the lipid effect and CHD signals are co-localized. The bottom panel shows disease associations at the locus with 103 clinical end points from UK Biobank and GWAS Consortia.

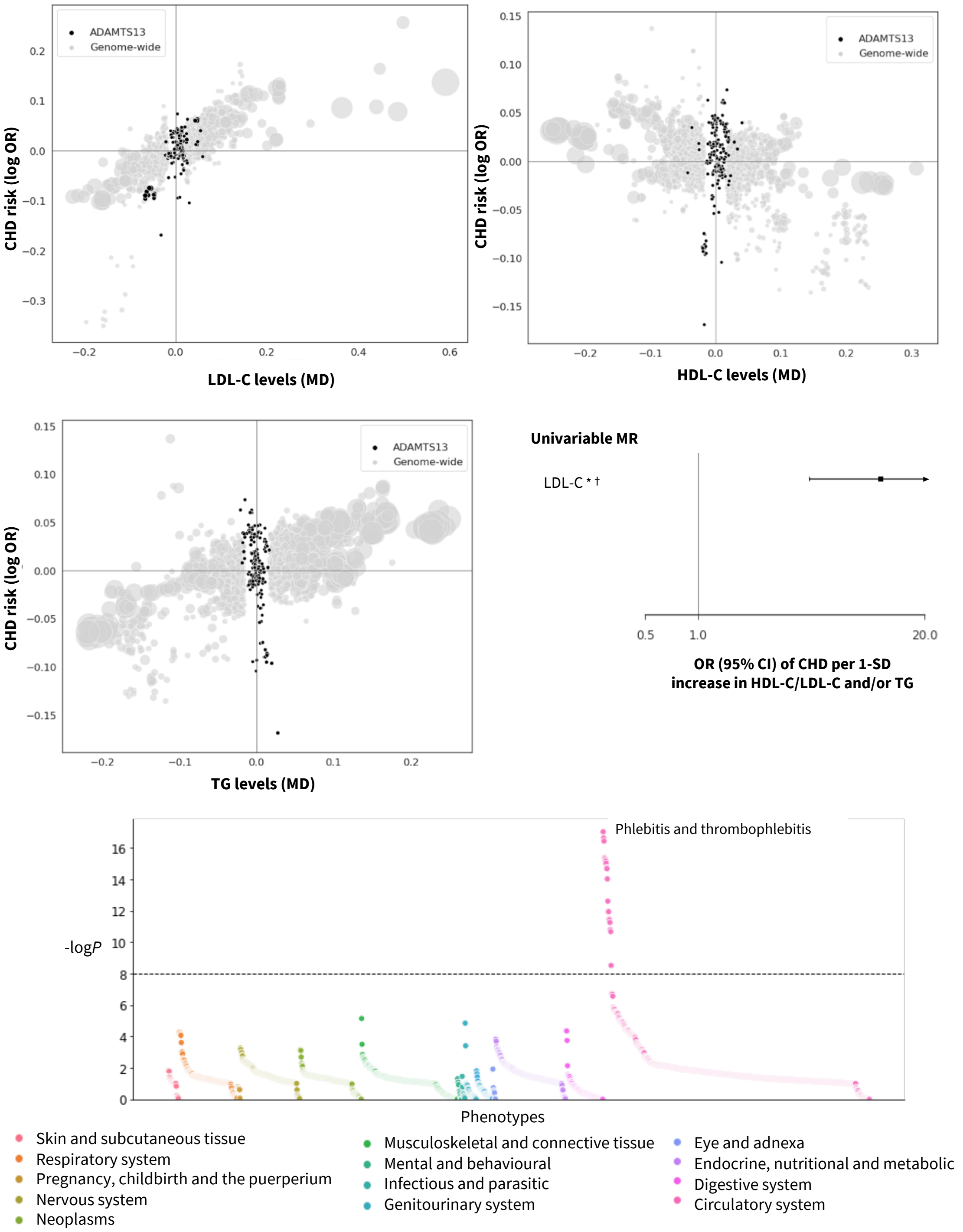


**Fig. S26. Prioritized target: ADAMTS13.** The top and middle left panels show genetic associations at the locus (± 50kbp) in black vs genome-wide associations (grey, *P* value < $1x10^{-6}$). The x-axis shows the per allele effect on the corresponding lipid expressed as mean difference (MD) from GLGC and the y-axis indicates the per allele effect on CHD expressed as log odds ratios (OR) from CardiogramPlusC4D. The marker size indicates the significance of the association with the lipid sub-fraction (*P*-value). The middle right panel shows the result of the univariable and multivariable (drug target) *cis*-MR results. An asterisk (*) indicates the MR estimates as being replicated, and a dagger (†) that the lipid effect and CHD signals are co-localized. The bottom panel shows disease associations at the locus with 103 clinical end points from UK Biobank and GWAS Consortia.

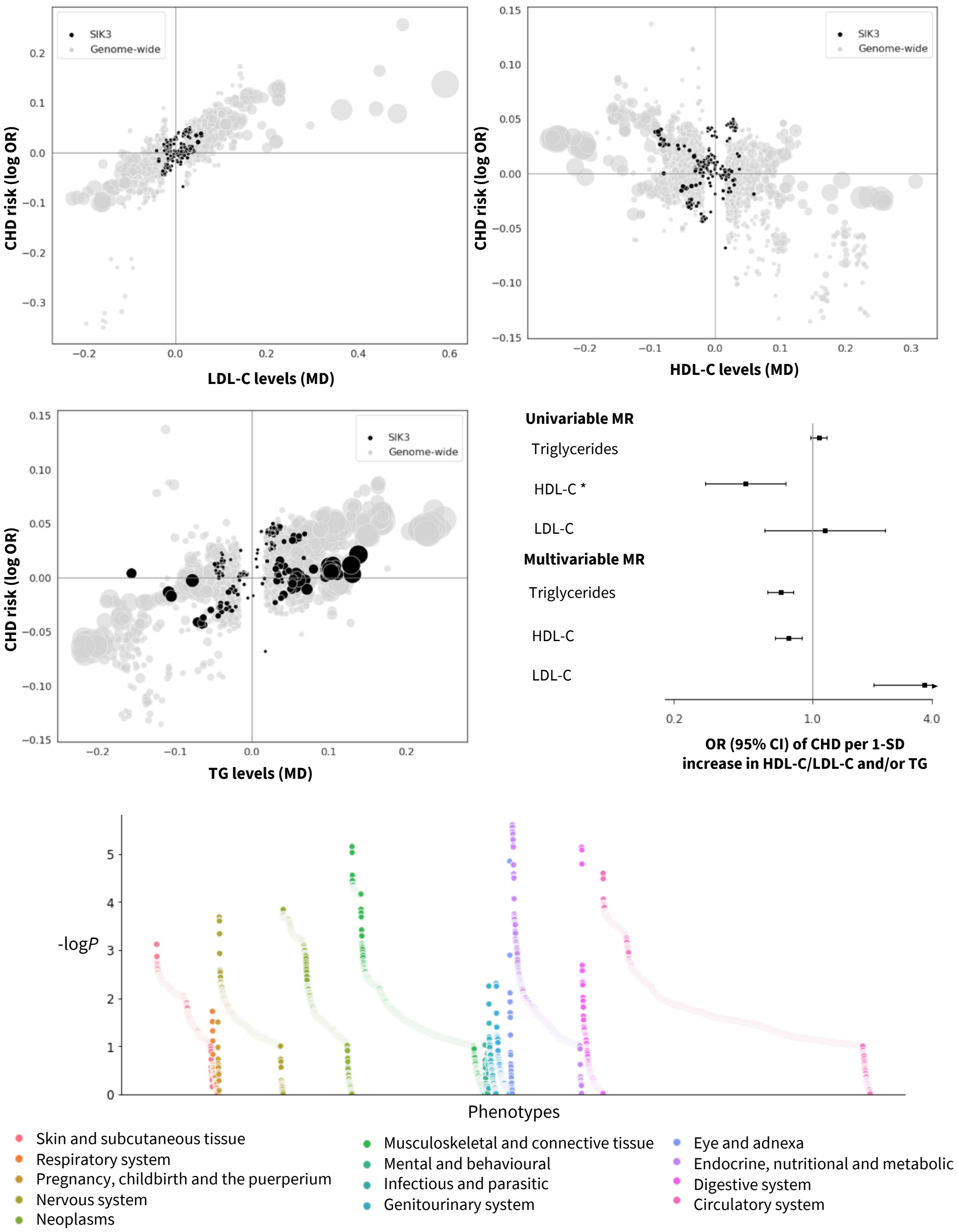


**Fig. S27. Prioritized target: SIK3.** The top and middle left panels show genetic associations at the locus (± 50kbp) in black vs genome-wide associations (grey, $P$ value < $1 \times 10^{-6}$). The x-axis shows the per allele effect on the corresponding lipid expressed as mean difference (MD) from GLGC and the y-axis indicates the per allele effect on CHD expressed as log odds ratios (OR) from CardiogramPlusC4D. The marker size indicates the significance of the association with the lipid sub-fraction ($P$-value). The middle right panel shows the result of the univariable and multivariable (drug target) *cis*-MR results. An asterisk (*) indicates the MR estimates as being replicated, and a dagger (†) that the lipid effect and CHD signals are co-localized. The bottom panel shows disease associations at the locus with 103 clinical end points from UK Biobank and GWAS Consortia.

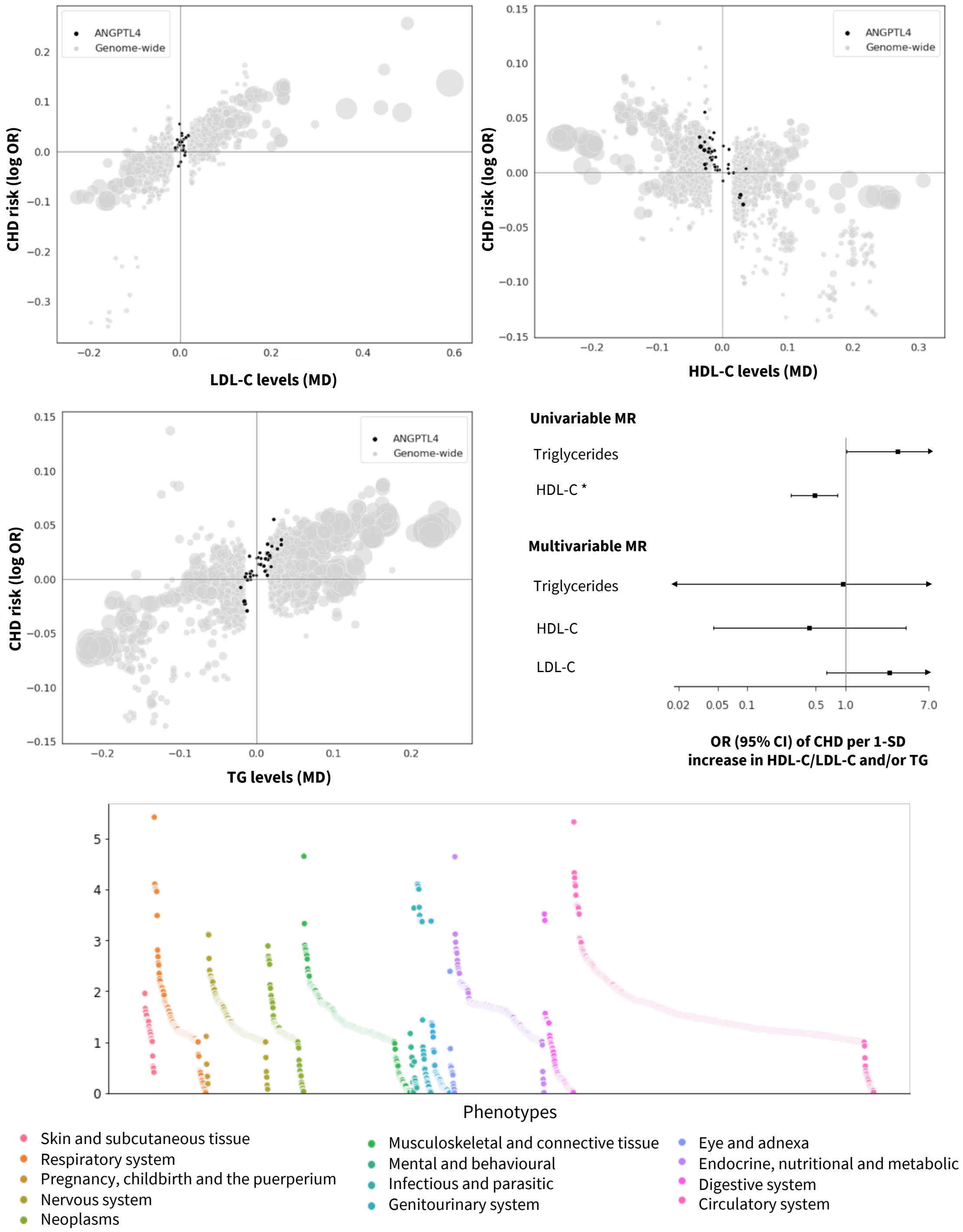


**Fig. S28. Prioritized target: ANGPTL4.** The top and middle left panels show genetic associations at the locus (± 50kbp) in black vs genome-wide associations (grey, $P$ value < $1\times10^{-6}$). The x-axis shows the per allele effect on the corresponding lipid expressed as mean difference (MD) from GLGC and the y-axis indicates the per allele effect on CHD expressed as log odds ratios (OR) from CardiogramPlusC4D. The marker size indicates the significance of the association with the lipid sub-fraction ($P$-value). The middle right panel shows the result of the univariable and multivariable (drug target) *cis*-MR results. An asterisk (*) indicates the MR estimates as being replicated, and a dagger (†) that the lipid effect and CHD signals are co-localized. The bottom panel shows disease associations at the locus with 103 clinical end points from UK Biobank and GWAS Consortia.

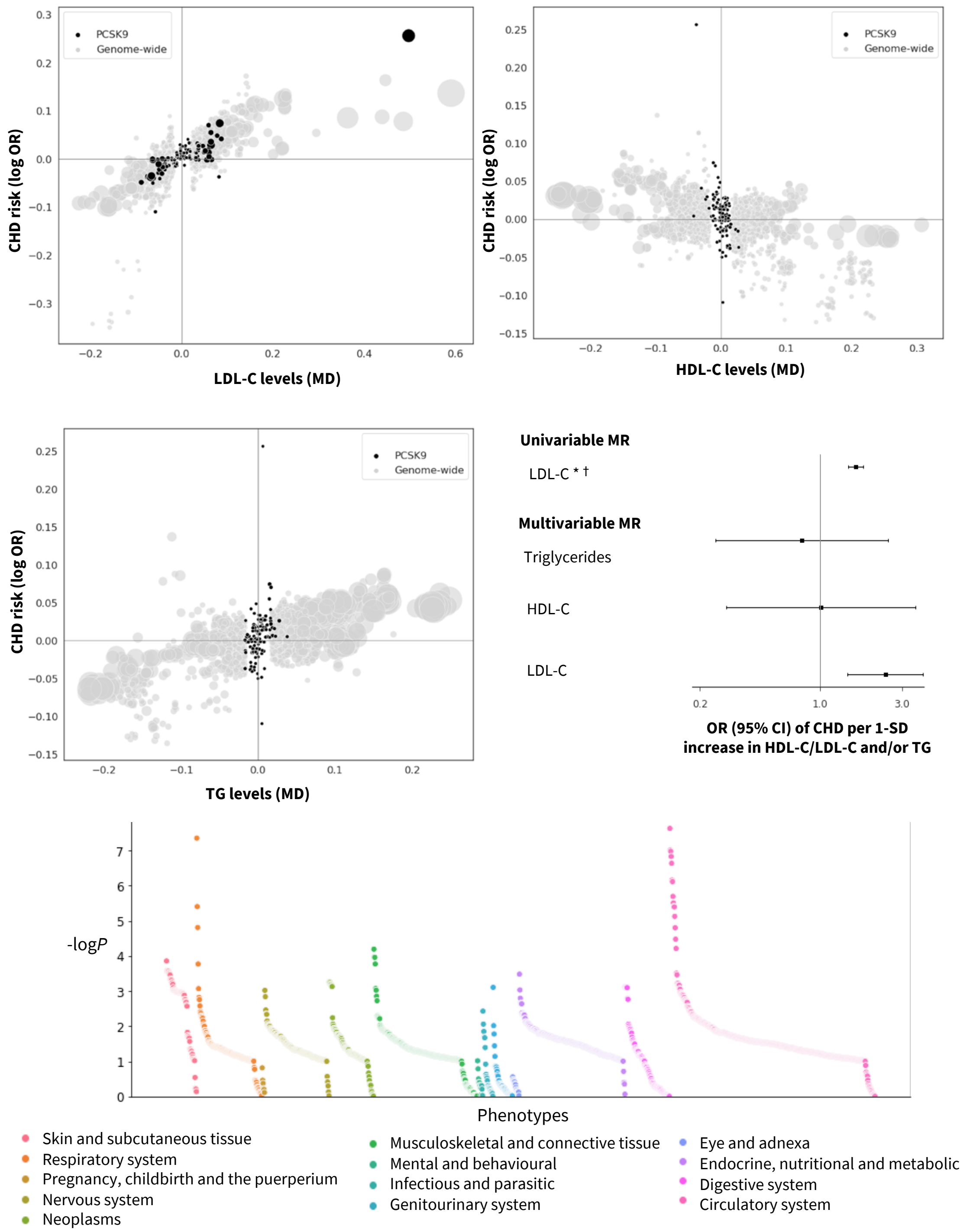


**Fig. S29. Prioritized target: PCSK9.** The top and middle left panels show genetic associations at the locus (± 50kbp) in black vs genome-wide associations (grey, $P$ value < $1x10^{-6}$). The x-axis shows the per allele effect on the corresponding lipid expressed as mean difference (MD) from GLGC and the y-axis indicates the per allele effect on CHD expressed as log odds ratios (OR) from CardiogramPlusC4D. The marker size indicates the significance of the association with the lipid sub-fraction ($P$-value). The middle right panel shows the result of the univariable and multivariable (drug target) *cis*-MR results. An asterisk (*) indicates the MR estimates as being replicated, and a dagger (†) that the lipid effect and CHD signals are co-localized. The bottom panel shows disease associations at the locus with 103 clinical end points from UK Biobank and GWAS Consortia.

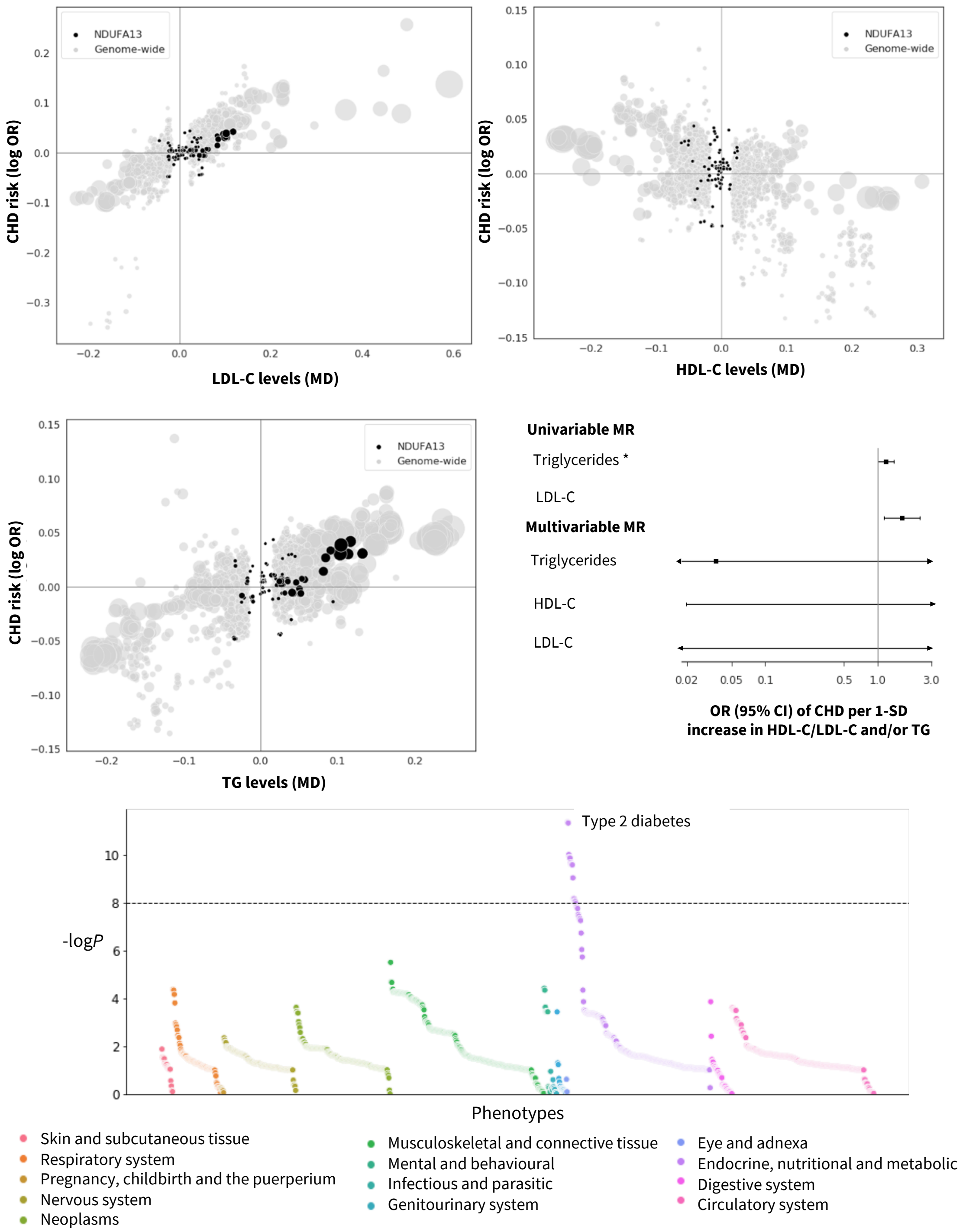


**Fig. S30. Prioritized target: NDUFA13.** The top and middle left panels show genetic associations at the locus (± 50kbp) in black vs genome-wide associations (grey, *P* value < 1x$10^{-6}$). The x-axis shows the per allele effect on the corresponding lipid expressed as mean difference (MD) from GLGC and the y-axis indicates the per allele effect on CHD expressed as log odds ratios (OR) from CardiogramPlusC4D. The marker size indicates the significance of the association with the lipid sub-fraction (*P*-value). The middle right panel shows the result of the univariable and multivariable (drug target) *cis*-MR results. An asterisk (*) indicates the MR estimates as being replicated, and a dagger (†) that the lipid effect and CHD signals are co-localized. The bottom panel shows disease associations at the locus with 103 clinical end points from UK Biobank and GWAS Consortia.

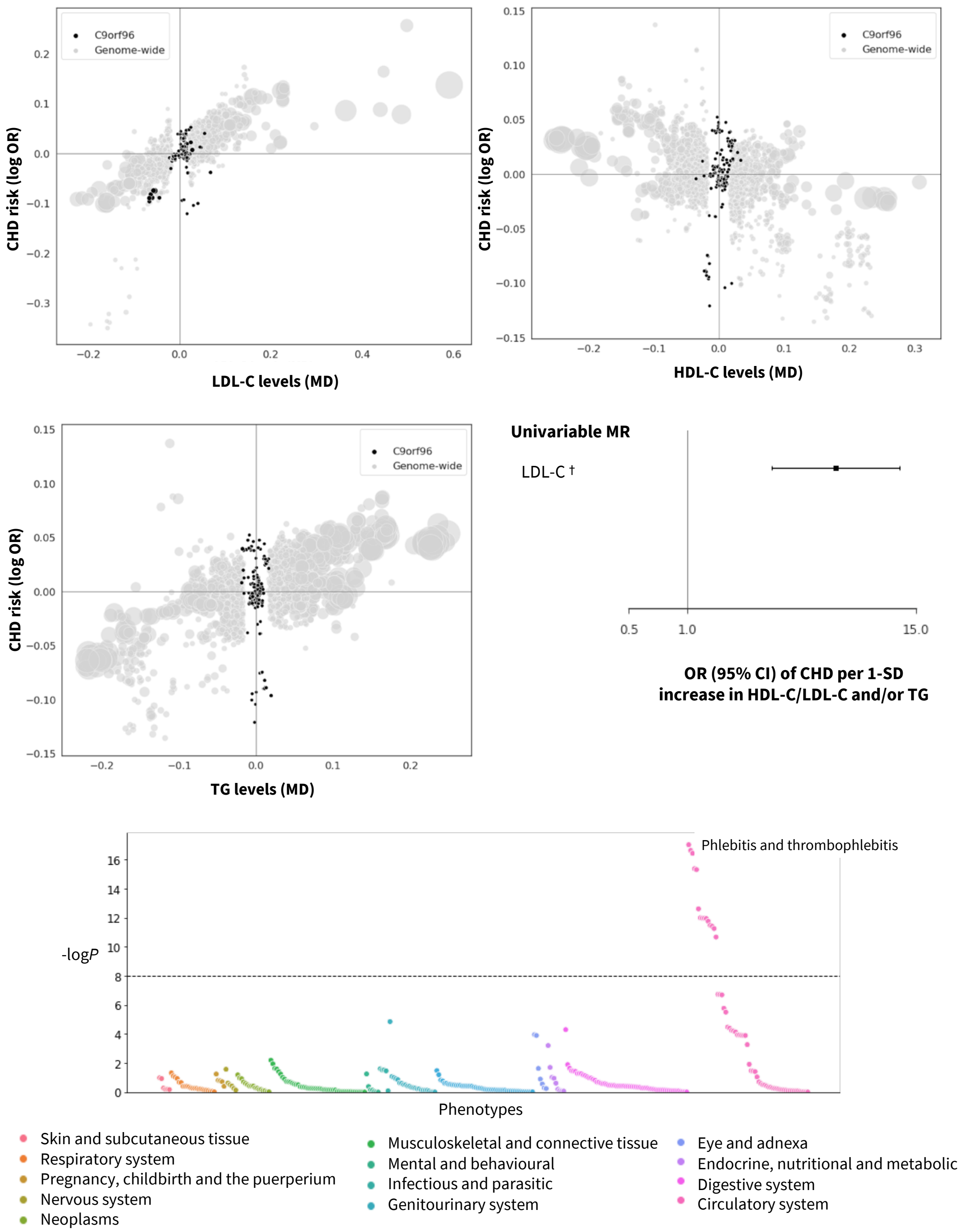


**Fig. S31. Prioritized target: C9orf96.** The top and middle left panels show genetic associations at the locus (± 50kbp) in black vs genome-wide associations (grey, $P$ value < $1x10^{-6}$). The x-axis shows the per allele effect on the corresponding lipid expressed as mean difference (MD) from GLGC and the y-axis indicates the per allele effect on CHD expressed as log odds ratios (OR) from CardiogramPlusC4D. The marker size indicates the significance of the association with the lipid sub-fraction ($P$-value). The middle right panel shows the result of the univariable and multivariable (drug target) *cis*-MR results. An asterisk (*) indicates the MR estimates as being replicated, and a dagger (†) that the lipid effect and CHD signals are co-localized. The bottom panel shows disease associations at the locus with 103 clinical end points from UK Biobank and GWAS Consortia.

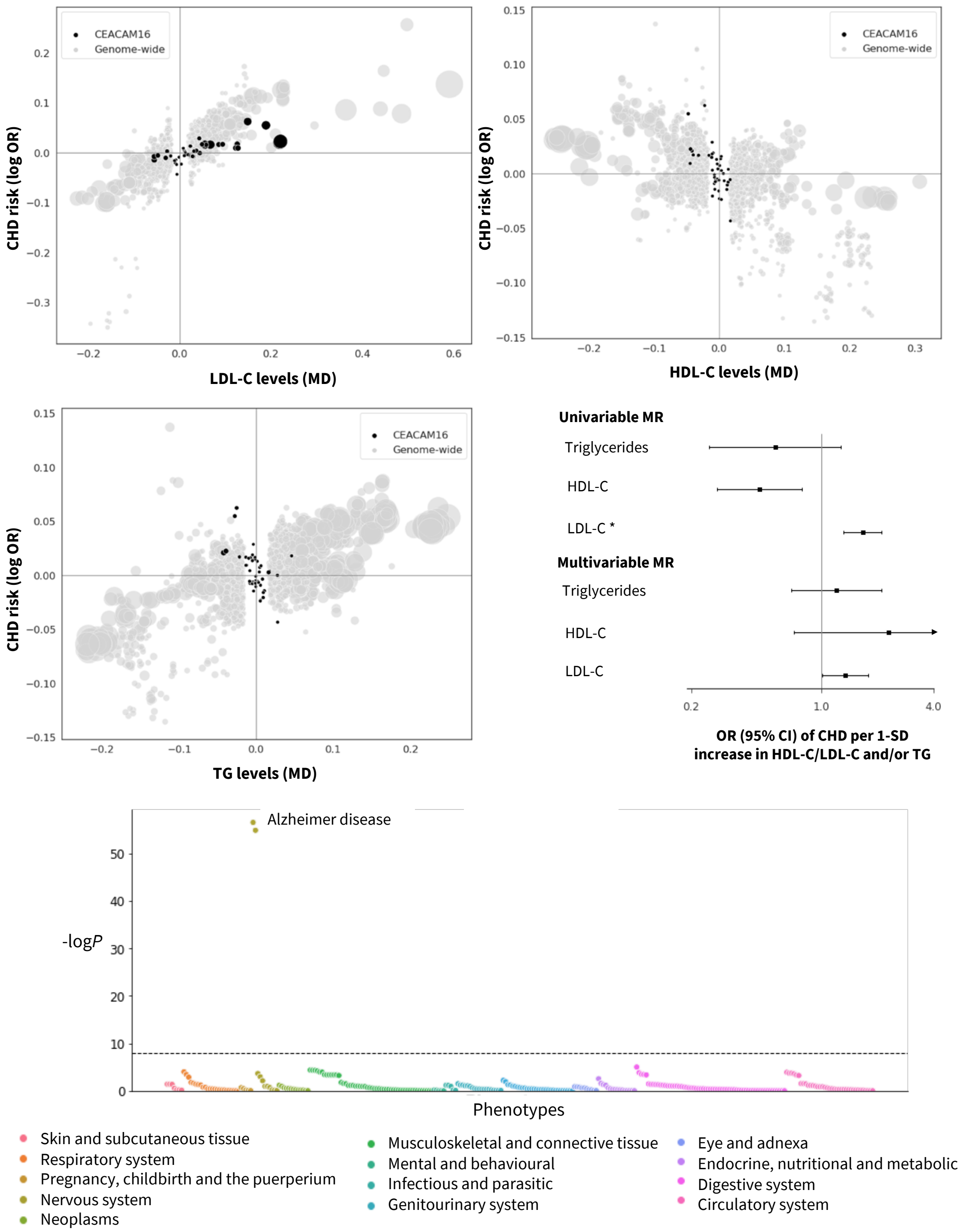


**Fig. S32. Prioritized target: CEACAM16.** The top and middle left panels show genetic associations at the locus (± 50kbp) in black vs genome-wide associations (grey, *P* value < 1x$10^{-6}$). The x-axis shows the per allele effect on the corresponding lipid expressed as mean difference (MD) from GLGC and the y-axis indicates the per allele effect on CHD expressed as log odds ratios (OR) from CardiogramPlusC4D. The marker size indicates the significance of the association with the lipid sub-fraction (*P*-value). The middle right panel shows the result of the univariable and multivariable (drug target) *cis*-MR results. An asterisk (*) indicates the MR estimates as being replicated, and a dagger (†) that the lipid effect and CHD signals are co-localized. The bottom panel shows disease associations at the locus with 103 clinical end points from UK Biobank and GWAS Consortia.

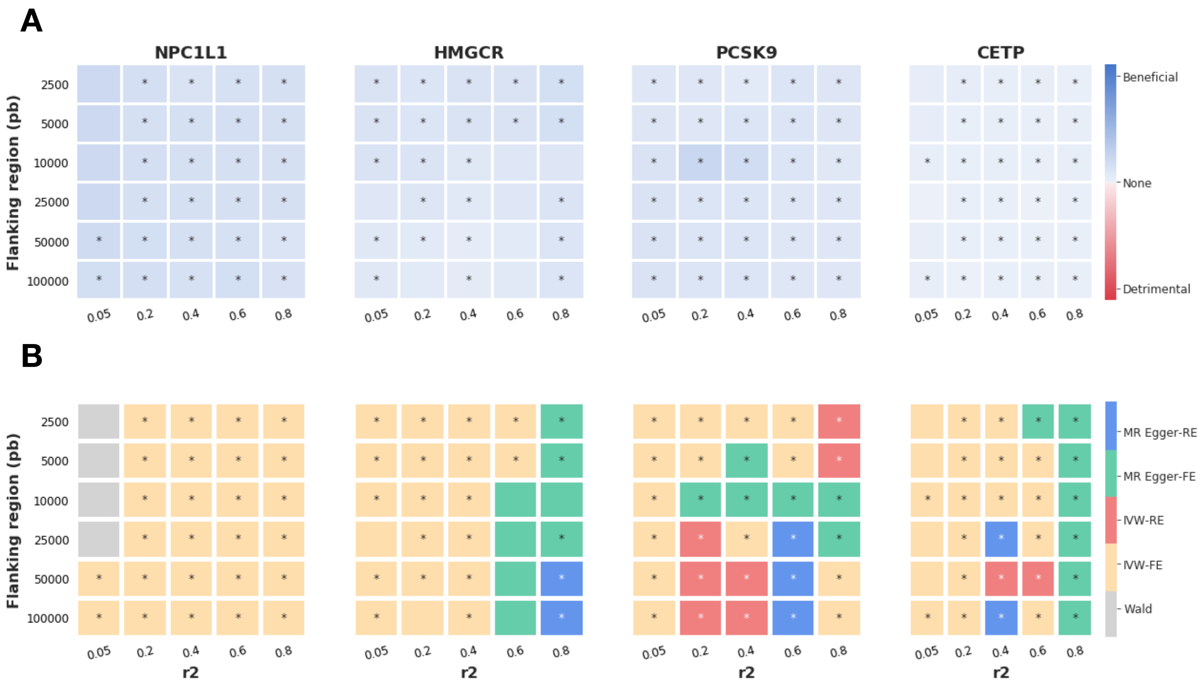


**Fig. S33. Drug target MR of positive control examples.** Grid search of LD threshold and region around the gene encoding a druggable target using genetic associations with LDL-C and HDL-C from the Global Lipid Genetic Consortium (GLGC) with CHD events from the CardiogramPlusC4D Consortium**.** MR estimates (**A**) and preferred model (**B**) for three licensed LDL-lowering drug targets and HDL-lowering CETP using lipid data from GLGC and CHD data from CardiogramPlusC4D in the discovery analysis. Models explored: MR Egger-RE (random effects), MR Egger-RE (fixed effects), inverse variance weighted (IVW)-RE (random effects), IVW-FE (fixed effects), Wald ratio.

**Table S1. Causal odds ratios (95% CI) for CHD per standard deviation increase in each lipid sub-fraction from a biomarker MR analysis.** All the studies used variants from the Global Lipid Genetic Consortium (GLGC) to instrument causal effects of LDL-C, HDL-C and triglycerides on CHD from the CardiogramPlusC4D Consortium. For details see text. OR = odds ratio per 1-SD increase in LDL-C/HDL-C or triglycerides; CI = confidence interval.

| **Method** | **LDL-C (OR, 95%CI)** | **HDL-C (OR, 95%CI)** | **Triglycerides (OR, 95%CI)** | **Ref.** |
|---|---|---|---|---|
| Regression-based method | 1.46 (1.37, 1.57)<br>No. variants = 185 | 0.96 (0.89, 1.03)<br>No. variants = 185 | 1.43 (1.28, 1.61)<br>No. variants = 185 | *(1)* |
| Multivariable IVW MR | 1.48 (1.36, 1.61)<br>No. variants = 185 | 0.93 (0.85, 1.02)<br>No. variants = 185 | 1.16 (1.04, 1.29)<br>No. variants = 185 | (2) |
| Univariable MR-Egger regression | 1.51 (1.30, 1.63)<br>No. variants = 418 | 0.95 (0.90, 1.01)<br>No. variants = 456 | 1.11 (1.01, 1.21)<br>No. variants = 381 | - |
| Multivariable MR-Egger regression | 1.53 (1.44, 1.62)<br>No. variants = 677 | 0.91 (0.86, 0.95)<br>n = 677 | 1.09 (1.01, 1.17)<br>No. variants = 677 | - |

**Table S2. Proximity to GWAS SNP, distance rank and previous evidence of druggable genes near genetic associations with LDL-C, HDL-C and TG.** For each druggable gene included in the analysis, the minimum distance from the gene to the variant (variants located within a gene were given a distance of 0bp and distance to variants upstream the gene are indicated with a negative value), a gene distance rank value according to their base pair distance, and indicated the druggable genes prioritized by GLGC are provided. OR = odds ratio per 1-SD increase in LDL-C/HDL-C or triglycerides; CI = confidence interval.

| Druggable gene | Genomic coordinates (GRCh37) | Min distance (pb) to LDL-C assoc. (distance rank) | Min distance (pb) to HDL-C assoc. (distance rank) | Min distance (pb) to TG assoc. (distance rank) | Prioritized by GLGC |
|---|---|---|---|---|---|
| ***RHD*** | 1:25598884-25656936 | 31340 (4) | - | - | No |
| ***RHCE*** | 1:25688740-25756683 | 0 (1) | - | - | No |
| ***RPS6KA1*** | 1:26856252-26901521 | - | 0 (1) | - | No |
| ***SFN*** | 1:27189633-27190947 | - | 45265 (6) | 45265 (6) | No |
| ***NR0B2*** | 1:27237980-27240457 | 44738 (4) | 44738 (4) | 44738 (4) | No |
| ***SLC9A1*** | 1:27425306-27493472 | -27702 (1) | - | - | No |
| ***BMP8A*** | 1:39957318-39991607 | - | -7238 (2) | -7238 (2) | No |
| ***PCSK9*** | 1:55505221-55530525 | 0 (1) | - | - | Yes |
| ***ANGPTL3*** | 1:63063158-63071830 | -49762 (2) | - | -7878 (2) | Yes |
| ***ATG4C*** | 1:63249806-63331184 | 0 (1) | - | 0 (1) | No |
| ***RPL5*** | 1:93297582-93307481 | -27758 (2) | - | - | No |
| ***CELSR2*** | 1:109792641-109818372 | 0 (1) | 0 (1) | - | No |
| ***PSMA5*** | 1:109941653-109969062 | -19276 (2) | - | - | No |
| ***GPR61*** | 1:110082494-110091028 | -20869 (3) | - | - | No |
| ***AMPD2*** | 1:110158726-110174673 | -49687 (4) | - | - | No |
| ***GSTM4*** | 1:110198703-110208118 | -29513 (3) | - | - | No |
| ***GSTM2*** | 1:110210644-110252171 | 44118 (4) | - | - | No |
| ***GSTM1*** | 1:110230436-110251661 | 44628 (5) | - | - | No |
| ***GSTM5*** | 1:110254864-110318050 | 0 (2) | - | - | No |
| ***GSTM3*** | 1:110276554-110284384 | 11905 (3) | - | - | No |
| ***CSF1*** | 1:110452864-110473614 | - | 5297 (1) | - | No |
| ***HDGF*** | 1:156711899-156736717 | - | -11248 (4) | - | No |
| ***GALNT2*** | 1:230193536-230417870 | - | 0 (1) | 0 (1) | Yes |
| ***GDF7*** | 2:20866424-20873418 | 13808 (2) | - | - | No |
| ***APOB*** | 2:21224301-21266945 | 29601 (1) | -28051 (1) | -28051 (1) | Yes |
| ***EMILIN1*** | 2:27301435-27309271 | - | - | 33623 (7) | No |
| ***KHK*** | 2:27309615-27323640 | - | - | 19254 (5) | No |
| ***CGREF1*** | 2:27321757-27341995 | - | - | 899 (1) | No |
| ***SLC5A6*** | 2:27422455-27435826 | - | - | 7370 (3) | No |
| ***ATRAID*** | 2:27434895-27440046 | - | - | 3150 (2) | No |
| ***CAD*** | 2:27440258-27466811 | - | - | 30737 (4) | No |
| ***UCN*** | 2:27530268-27531313 | - | - | -32720 (5) | No |
| ***NRBP1*** | 2:27650657-27665126 | - | - | -10332 (2) | No |
| ***GCKR*** | 2:27719709-27746554 | 0 (1) | - | 32018 (4) | Yes |
| ***MAP3K19*** | 2:135722061-135805038 | 0 (1) | - | - | No |
| ***LCT*** | 2:136545410-136594750 | 0 (1) | - | - | No |
| ***ABCB11*** | 2:169779448-169887832 | 0 (1) | - | - | Yes |
| ***CPS1*** | 2:211342406-211543831 | - | 0 (1) | - | Yes |
| ***FN1*** | 2:216225163-216300895 | 0 (1) | - | - | Yes |
| ***UGT1A8*** | 2:234526291-234681956 | 0 (9) | - | - | No |
| ***UGT1A10*** | 2:234545100-234681951 | 0 (8) | - | - | No |
| ***UGT1A9*** | 2:234580499-234681946 | 0 (7) | - | - | No |
| ***UGT1A7*** | 2:234590584-234681945 | 0 (4) | - | - | No |
| ***UGT1A6*** | 2:234600253-234681946 | 0 (6) | - | - | No |
| ***UGT1A5*** | 2:234621638-234681945 | 0 (5) | - | - | No |
| ***UGT1A4*** | 2:234627424-234681945 | 0 (3) | - | - | No |
| ***UGT1A3*** | 2:234637754-234681945 | 0 (2) | - | - | No |
| ***UGT1A1*** | 2:234668894-234681945 | 0 (1) | - | - | Yes |
| ***PPARG*** | 3:12328867-12475855 | 0 (1) | 0 (1) | 13487 (1) | No |
| ***RAF1*** | 3:12625100-12705725 | 0 (1) | - | - | Yes |
| ***CAMKV*** | 3:49895421-49907655 | - | 0 (1) | - | No |
| ***MST1R*** | 3:49924435-49941299 | - | 30215 (4) | - | No |
| ***SEMA3F*** | 3:50192478-50226508 | - | -20081 (2) | - | No |
| ***SEMA3G*** | 3:52467069-52479101 | - | -35398 (4) | - | No |
| ***TNNC1*** | 3:52485118-52488086 | - | 4621 (2) | - | No |
| ***NISCH*** | 3:52489134-52527087 | - | 0 (1) | - | No |

| Druggable gene | Genomic coordinates (GRCh37) | Min distance (pb) to LDL-C assoc. (distance rank) | Min distance (pb) to HDL-C assoc. (distance rank) | Min distance (pb) to TG assoc. (distance rank) | Prioritized by GLGC |
|---|---|---|---|---|---|
| ***PBRM1*** | 3:52579368-52719933 | - | 0 (1) | - | No |
| ***NEK4*** | 3:52744800-52804965 | - | 39569 (8) | - | No |
| ***ITIH1*** | 3:52811603-52826078 | - | 18456 (4) | - | No |
| ***ITIH3*** | 3:52828784-52843025 | - | 1509 (1) | - | No |
| ***ITIH4*** | 3:52846991-52865495 | - | -2457 (2) | - | No |
| ***ABHD6*** | 3:58223233-58281420 | 12447 (3) | - | - | No |
| ***NR1I2*** | 3:119499331-119537332 | - | 4965 (2) | - | No |
| ***GSK3B*** | 3:119540170-119813264 | - | 0 (1) | - | Yes |
| ***RGS12*** | 4:3294755-3441640 | 0 (1) | - | 31499 (3) | No |
| ***HGFAC*** | 4:3443614-3451211 | -8729 (2) | - | 21928 (2) | No |
| ***LRPAP1*** | 4:3508103-3534286 | -34964 (4) | - | -34964 (4) | Yes |
| ***MAPK10*** | 4:86936276-87515284 | - | - | 0 (1) | No |
| ***PTPN13*** | 4:87515468-87736324 | - | 32193 (3) | -41692 (2) | No |
| ***KLHL8*** | 4:88081255-88161466 | - | -28170 (2) | 49590 (3) | Yes |
| ***HSD17B11*** | 4:88257762-88312538 | - | - | -46706 (2) | No |
| ***METAP1*** | 4:99916771-99983964 | - | 30841 (3) | - | No |
| ***ADH5*** | 4:99992132-100009952 | - | 4853 (1) | - | Yes |
| ***ADH4*** | 4:100044808-100078949 | - | -30003 (2) | - | No |
| ***NFKB1*** | 4:103422486-103538459 | - | -49200 (2) | - | No |
| ***PDGFC*** | 4:157681606-157892546 | - | 0 (1) | - | No |
| ***HMGCR*** | 5:74632154-74657929 | -49992 (2) | - | - | Yes |
| ***CSNK1G3*** | 5:122847793-122952739 | 0 (1) | - | - | Yes |
| ***HIST1H1C*** | 6:26055968-26056699 | 36442 (6) | - | - | No |
| ***HFE*** | 6:26087509-26098571 | 0 (1) | - | - | Yes |
| ***HIST1H4C*** | 6:26104104-26104518 | -10963 (2) | - | - | No |
| ***OR11A1*** | 6:29393281-29424848 | - | - | 17853 (3) | No |
| ***OR2H1*** | 6:29424958-29432105 | - | - | 10596 (1) | No |
| ***MAS1L*** | 6:29454474-29455738 | - | - | -11773 (2) | No |
| ***HLA-G*** | 6:29794744-29798902 | - | - | 21684 (1) | No |
| ***TUBB*** | 6:30687978-30693203 | - | - | 15752 (3) | No |
| ***DDR1*** | 6:30844198-30867933 | - | - | -45501 (1) | No |
| ***SFTA2*** | 6:30899130-30899952 | - | - | 20172 (2) | No |
| ***C6orf15*** | 6:31079000-31080336 | 49371 (8) | 25077 (6) | 32878 (7) | No |
| ***HLA-C*** | 6:31236526-31239907 | 32354 (1) | - | 25632 (1) | No |
| ***HLA-B*** | 6:31321649-31324965 | 29139 (2) | 29139 (2) | -49933 (2) | No |
| ***LTA*** | 6:31539831-31542101 | -27032 (6) | - | 930 (2) | No |
| ***TNF*** | 6:31543344-31546113 | -30545 (7) | - | -313 (1) | No |
| ***LTB*** | 6:31548302-31550299 | -35503 (9) | - | 45839 (12) | No |
| ***NCR3*** | 6:31556672-31560762 | -43873 (11) | 41081 (10) | 35376 (7) | No |
| ***APOM*** | 6:31620193-31625987 | - | -18350 (4) | 39207 (17) | No |
| ***ABHD16A*** | 6:31654726-31671221 | - | -35150 (11) | 0 (1) | No |
| ***C6orf25*** | 6:31686371-31694491 | - | - | -21177 (8) | No |
| ***HSPA1A*** | 6:31783291-31785723 | - | - | 22713 (5) | No |
| ***HSPA1B*** | 6:31795512-31798031 | 48981 (9) | - | 39246 (7) | No |
| ***NEU1*** | 6:31825436-31830683 | 17537 (4) | - | 20671 (5) | No |
| ***SLC44A4*** | 6:31830969-31846823 | 1397 (2) | - | 4531 (2) | No |
| ***EHMT2*** | 6:31847536-31865464 | 0 (1) | - | 0 (1) | No |
| ***C2*** | 6:31865562-31913449 | 15565 (7) | - | -14208 (3) | No |
| ***CFB*** | 6:31895475-31919861 | 9153 (4) | - | -44121 (8) | No |
| ***C4A*** | 6:31949801-31970458 | -20787 (8) | - | -20787 (8) | No |
| ***C4B*** | 6:31982539-32003195 | -35079 (10) | - | 4264 (3) | No |
| ***CYP21A2*** | 6:32006042-32009447 | 36828 (3) | - | 0 (1) | No |
| ***TNXB*** | 6:32008931-32083111 | 0 (1) | - | -1472 (2) | No |
| ***EGFL8*** | 6:32132360-32136058 | - | - | 48287 (8) | No |
| ***AGER*** | 6:32148745-32152101 | - | 37740 (4) | 32244 (4) | No |
| ***NOTCH4*** | 6:32162620-32191844 | - | 0 (1) | 0 (1) | No |
| ***HLA-DRA*** | 6:32407619-32412823 | -48188 (3) | -44404 (3) | 0 (1) | No |
| ***HLA-DRB5*** | 6:32485120-32498064 | - | -40922 (2) | - | No |
| ***HLA-DRB1*** | 6:32546546-32557625 | 42432 (3) | 18358 (1) | 42432 (3) | No |
| ***HLA-DQA2*** | 6:32709119-32714992 | -37979 (2) | -39746 (2) | -24862 (1) | No |
| ***HLA-DOB*** | 6:32780540-32784825 | 0 (1) | - | -25250 (2) | No |
| ***PSMB8*** | 6:32808494-32812480 | -23874 (4) | - | - | No |
| ***PSMB9*** | 6:32811913-32827362 | -27293 (5) | - | - | No |
| ***SCUBE3*** | 6:35182190-35220856 | -49116 (2) | -20049 (1) | - | No |
| ***KCNK17*** | 6:39266777-39282329 | -15940 (1) | - | - | Yes |
| ***KCNK16*** | 6:39282474-39290744 | -31637 (2) | - | - | No |
| ***VEGFA*** | 6:43737921-43754224 | - | 10327 (1) | 10327 (1) | Yes |
| ***FRK*** | 6:116252312-116381921 | 0 (1) | - | - | Yes |
| ***RSPO3*** | 6:127439749-127518910 | - | 2217 (1) | -47328 (1) | Yes |
| ***L3MBTL3*** | 6:130334844-130462594 | - | - | 0 (1) | No |
| ***ESR1*** | 6:151977826-152450754 | - | 0 (1) | - | No |
| ***IGF2R*** | 6:160390131-160534539 | 8609 (2) | - | - | No |

| Druggable gene | Genomic coordinates (GRCh37) | Min distance (pb) to LDL-C assoc. (distance rank) | Min distance (pb) to HDL-C assoc. (distance rank) | Min distance (pb) to TG assoc. (distance rank) | Prioritized by GLGC |
|---|---|---|---|---|---|
| ***SLC22A1*** | 6:160542821-160579750 | 0 (1) | - | - | No |
| ***SLC22A2*** | 6:160592093-160698670 | 19974 (1) | 69 (1) | - | No |
| ***SLC22A3*** | 6:160769300-160876014 | -46295 (2) | -32178 (1) | 0 (1) | No |
| ***LPA*** | 6:160952515-161087407 | 0 (1) | 0 (1) | -45381 (2) | Yes |
| ***PLG*** | 6:161123270-161174347 | -40471 (2) | -40809 (2) | - | No |
| ***MAP3K4*** | 6:161412759-161538417 | -21972 (1) | - | - | No |
| ***GPR146*** | 7:1084212-1098897 | - | -435 (2) | - | Yes |
| ***GPER1*** | 7:1121844-1133451 | - | -38067 (3) | - | No |
| ***DAGLB*** | 7:6448757-6523821 | - | -13853 (2) | - | Yes |
| ***NPC1L1*** | 7:44552134-44580914 | 41372 (4) | - | - | Yes |
| ***FKBP6*** | 7:72742167-72772634 | - | - | 0 (1) | No |
| ***FZD9*** | 7:72848109-72850450 | - | 5980 (2) | 32656 (2) | No |
| ***STX1A*** | 7:73113536-73134002 | - | - | -53 (2) | No |
| ***MET*** | 7:116312444-116438440 | - | - | 0 (1) | Yes |
| ***AOC1*** | 7:150521715-150558592 | - | 0 (1) | - | No |
| ***TNKS*** | 8:9413424-9639856 | - | - | 0 (1) | No |
| ***BLK*** | 8:11351510-11422113 | - | - | 0 (1) | No |
| ***FDFT1*** | 8:11653082-11696818 | - | - | -36672 (4) | No |
| ***CTSB*** | 8:11700033-11726957 | - | - | -10805 (2) | No |
| ***NAT2*** | 8:18248755-18258728 | 13710 (1) | - | 9752 (1) | Yes |
| ***LPL*** | 8:19759228-19824769 | - | 46744 (1) | 46744 (1) | Yes |
| ***SLC18A*** | 8:20002366-20040717 | - | -46760 (1) | -41092 (1) | Yes |
| ***CYP7A1*** | 8:59402737-59412795 | -4276 (1) | - | -49203 (2) | Yes |
| ***GPIHBP1*** | 8:144295068-144299044 | - | -49193 (2) | - | Yes |
| ***ABCA1*** | 9:107543283-107690518 | 0 (1) | 0 (1) | 0 (1) | Yes |
| ***OBP2B*** | 9:136080664-136084630 | 47720 (1) | - | - | No |
| ***RPL7A*** | 9:136215069-136218281 | -24059 (3) | - | - | No |
| ***C9orf96*** | 9:136243117-136271220 | 0 (1) | - | - | No |
| ***ADAMTS13*** | 9:136279478-136324508 | 1740 (2) | - | - | No |
| ***AKR1C3*** | 10:5077546-5149878 | - | - | 46395 (3) | No |
| ***VIM*** | 10:17270258-17279592 | -9968 (1) | - | - | No |
| ***ALOX5*** | 10:45869661-45941561 | - | 0 (1) | - | No |
| ***CYP26C1*** | 10:94821021-94828454 | - | -15356 (2) | -15356 (2) | No |
| ***CYP26A1*** | 10:94833232-94837647 | - | -27567 (3) | -27567 (3) | Yes |
| ***TECTB*** | 10:114043493-114064793 | - | 0 (1) | 0 (1) | No |
| ***ADRB1*** | 10:115803806-115806667 | - | -11019 (1) | - | No |
| ***AMPD3*** | 11:10329860-10529126 | - | 0 (1) | - | Yes |
| ***PSMA1*** | 11:14515329-14665181 | - | -10866 (2) | - | No |
| ***LGR4*** | 11:27387508-27494322 | - | - | 31783 (2) | No |
| ***CHST1*** | 11:45670427-45687172 | - | 47215 (1) | - | No |
| ***CRY2*** | 11:45868669-45904798 | - | -28960 (2) | - | No |
| ***F2*** | 11:46740730-46761056 | - | 42729 (2) | -18509 (3) | No |
| ***ACP2*** | 11:47260853-47270457 | - | 40910 (3) | -34365 (4) | No |
| ***NR1H3*** | 11:47269851-47290396 | - | 20971 (2) | -43363 (6) | No |
| ***PSMC3*** | 11:47440320-47447993 | - | 1551 (1) | - | No |
| ***NDUFS3*** | 11:47586888-47606114 | - | -3767 (2) | - | No |
| ***PTPRJ*** | 11:48002113-48189670 | - | 0 (1) | - | No |
| ***FOLH1*** | 11:49168187-49230222 | - | 0 (1) | - | No |
| ***OR4A16*** | 11:55110627-55111707 | - | 12013 (2) | - | No |
| ***OR4C16*** | 11:55339604-55340536 | - | -15296 (2) | - | No |
| ***DAGLA*** | 11:61447905-61514473 | 49826 (6) | 49826 (6) | 49826 (6) | No |
| ***FEN1*** | 11:61560109-61564716 | 15044 (3) | 15044 (3) | 15044 (3) | No |
| ***KCNK7*** | 11:65360326-65363467 | - | 27850 (4) | - | No |
| ***MAP3K11*** | 11:65365226-65382853 | - | 8464 (2) | - | No |
| ***RELA*** | 11:65421067-65430565 | - | -29750 (5) | - | No |
| ***MOGAT2*** | 11:75428864-75444003 | - | 11018 (1) | - | No |
| ***APOA5*** | 11:116660083-116663136 | -4483 (2) | -4483 (2) | -43681 (3) | No |
| ***APOA4*** | 11:116691419-116694022 | -35819 (4) | -35819 (4) | -35819 (4) | No |
| ***APOC3*** | 11:116700422-116703788 | -44822 (5) | -44822 (5) | -44822 (5) | No |
| ***APOA1*** | 11:116706467-116708666 | -42616 (6) | -38922 (6) | -42616 (6) | Yes |
| ***SIK3*** | 11:116714118-116969153 | 0 (1) | 0 (1) | -46573 (7) | No |
| ***SIDT2*** | 11:117049449-117068160 | 7406 (3) | -3252 (2) | -23075 (2) | No |
| ***PCSK7*** | 11:117075053-117103241 | 0 (1) | -28856 (4) | -48679 (4) | No |
| ***BACE1*** | 11:117156402-117186975 | - | 0 (1) | -4990 (2) | No |
| ***DCPS*** | 11:126173647-126215644 | 26052 (2) | 12356 (2) | - | No |
| ***ST3GAL4*** | 11:126225535-126310239 | 0 (1) | 0 (1) | - | Yes |
| ***PDE3A*** | 12:20522179-20837315 | - | -48421 (1) | - | Yes |
| ***SLCO1B1*** | 12:21284136-21392180 | - | - | 0 (1) | No |
| ***BAZ2A*** | 12:56989380-57030600 | - | - | -38161 (2) | No |
| ***INHBC*** | 12:57828543-57844611 | - | 0 (1) | 0 (1) | No |
| ***INHBE*** | 12:57846106-57853063 | - | -2057 (2) | -2057 (2) | No |
| ***MARS*** | 12:57869228-57911352 | - | -25179 (6) | -25179 (6) | No |

| Druggable gene | Genomic coordinates (GRCh37) | Min distance (pb) to LDL-C assoc. (distance rank) | Min distance (pb) to HDL-C assoc. (distance rank) | Min distance (pb) to TG assoc. (distance rank) | Prioritized by GLGC |
|---|---|---|---|---|---|
| ***PIP4K2C*** | 12:57984957-57997198 | - | 15035 (5) | - | No |
| ***ALDH2*** | 12:112204691-112247782 | 37809 (2) | - | - | No |
| ***MAPKAPK5*** | 12:112279782-112334343 | 5500 (1) | - | - | No |
| ***ERP29*** | 12:112451120-112461255 | -10389 (2) | 25563 (2) | - | No |
| ***RPL6*** | 12:112842994-112856642 | -29579 (2) | 49773 (2) | - | No |
| ***PTPN11*** | 12:112856155-112947717 | 0 (1) | 0 (1) | - | No |
| ***HPD*** | 12:122277433-122301502 | - | - | -28342 (4) | No |
| ***HCAR1*** | 12:123104824-123215390 | - | 0 (2) | - | No |
| ***HCAR2*** | 12:123185840-123187890 | - | 12878 (3) | - | No |
| ***HCAR3*** | 12:123199303-123201439 | - | 0 (1) | - | No |
| ***SCARB1*** | 12:125261402-125367214 | - | 0 (1) | - | Yes |
| ***CBLN3*** | 14:24895738-24900160 | -12108 (2) | - | - | No |
| ***SERPINA10*** | 14:94749650-94759608 | 35884 (2) | - | - | No |
| ***SERPINA6*** | 14:94770585-94789731 | 5761 (1) | - | - | No |
| ***SERPINA1*** | 14:94843084-94857030 | -47592 (3) | - | - | No |
| ***AKT1*** | 14:105235686-105262088 | - | 15121 (2) | - | No |
| ***LTK*** | 15:41795836-41806085 | - | 23145 (3) | - | No |
| ***TYRO3*** | 15:41849873-41871536 | - | -20643 (2) | - | No |
| ***GANC*** | 15:42565431-42645864 | - | 37923 (3) | 37923 (3) | No |
| ***CATSPER2*** | 15:43920701-43960316 | - | - | -26883 (4) | No |
| ***PDIA3*** | 15:44038590-44065477 | - | - | -22173 (2) | No |
| ***MFAP1*** | 15:44096690-44117000 | - | - | 35817 (3) | No |
| ***ALDH1A2*** | 15:58245622-58790065 | - | 0 (1) | 0 (1) | No |
| ***LIPC*** | 15:58702768-58861151 | - | -21963 (2) | -22125 (2) | Yes |
| ***ADAM10*** | 15:58887403-59042177 | - | -34294 (2) | - | No |
| ***LACTB†*** | 15:63413999-63434260 | - | -498 (1) | -498 (1) | Yes |
| ***PKM*** | 15:72491370-72524164 | - | - | 42451 (4) | No |
| ***HSD3B7*** | 16:30996519-31000473 | - | - | 47606 (8) | No |
| ***PRSS53*** | 16:31094746-31100949 | - | - | 41044 (7) | No |
| ***VKORC1*** | 16:31102163-31107301 | - | - | 34692 (5) | No |
| ***PRSS8*** | 16:31142756-31147083 | - | - | -763 (2) | No |
| ***PRSS36*** | 16:31150246-31161415 | - | - | -8253 (3) | No |
| ***SLC12A3*** | 16:56899119-56949762 | 43263 (4) | 43263 (4) | 37253 (4) | No |
| ***CETP*** | 16:56995762-57017757 | -2737 (1) | -2737 (1) | -8747 (1) | Yes |
| ***CCL22*** | 16:57392684-57400102 | - | -38750 (2) | - | No |
| ***CES3*** | 16:66995140-67009051 | - | 42996 (3) | - | No |
| ***CES4A*** | 16:67022492-67043661 | - | 8386 (1) | - | No |
| ***HSD11B2*** | 16:67464555-67471456 | - | -45403 (4) | - | No |
| ***AGRP*** | 16:67516474-67517716 | - | 37623 (2) | - | No |
| ***GFOD2*** | 16:67708434-67753324 | - | 5454 (2) | - | No |
| ***PSKH1*** | 16:67927175-67963581 | - | -42556 (7) | - | No |
| ***CTRL*** | 16:67961543-67966317 | - | 27326 (6) | - | No |
| ***PSMB10*** | 16:67968405-67970990 | - | 22653 (4) | - | No |
| ***LCAT*** | 16:67973653-67978034 | - | 46961 (6) | - | Yes |
| ***SLC12A4*** | 16:67977377-68003504 | - | 21491 (4) | - | No |
| ***DPEP3*** | 16:68009566-68014732 | - | 10263 (3) | - | No |
| ***DPEP2*** | 16:68021297-68034489 | - | 0 (2) | - | No |
| ***PLA2G15*** | 16:68279207-68294961 | - | 0 (1) | - | No |
| ***DHODH*** | 16:72042487-72058954 | -34262 (1) | - | 49139 (6) | No |
| ***HP*** | 16:72088491-72094954 | 0 (2) | - | 13139 (3) | No |
| ***ASGR1*** | 17:7076750-7082883 | 6040 (2) | - | - | No |
| ***AURKB*** | 17:8108056-8113918 | 47231 (5) | - | - | No |
| ***CACNB1*** | 17:37329709-37353956 | - | 35453 (4) | - | No |
| ***RPL19*** | 17:37356536-37360980 | - | 28429 (3) | - | No |
| ***CDK12*** | 17:37617764-37721160 | - | 18114 (1) | - | No |
| ***PNMT*** | 17:37824234-37826728 | - | 31950 (4) | - | No |
| ***ERBB2*** | 17:37844167-37886679 | - | 0 (1) | - | No |
| ***PSMD3*** | 17:38137050-38154213 | - | -15057 (2) | - | No |
| ***CSF3*** | 17:38171614-38174066 | - | -49621 (6) | - | No |
| ***SOST*** | 17:41831099-41836156 | - | - | 42010 (4) | No |
| ***DUSP3*** | 17:41843489-41856356 | - | - | 21810 (3) | No |
| ***CD300LG*** | 17:41924516-41940997 | - | 0 (1) | 0 (1) | No |
| ***PPY*** | 17:42018172-42019836 | - | -39416 (5) | - | No |
| ***WNT9B*** | 17:44910567-44964096 | - | 46625 (4) | - | No |
| ***MYL4*** | 17:45277812-45301045 | 12133 (1) | - | - | No |
| ***ITGB3*** | 17:45331212-45421658 | 3457 (2) | - | - | No |
| ***NPEPPS*** | 17:45600308-45700642 | 49954 (3) | - | 49954 (3) | No |
| ***APOH*** | 17:64208151-64252643 | 0 (1) | - | - | No |
| ***ITGB4*** | 17:73717408-73753899 | 28292 (4) | - | - | No |
| ***GALK1*** | 17:73747675-73761792 | 20399 (3) | - | - | No |
| ***H3F3B*** | 17:73772515-73781974 | 217 (2) | - | - | No |
| ***RPL17*** | 18:47014851-47018906 | - | 10670 (1) | - | No |

| Druggable gene | Genomic coordinates (GRCh37) | Min distance (pb) to LDL-C assoc. (distance rank) | Min distance (pb) to HDL-C assoc. (distance rank) | Min distance (pb) to TG assoc. (distance rank) | Prioritized by GLGC |
|---|---|---|---|---|---|
| ***LIPG*** | 18:47087069-47119272 | 37732 (1) | 40585 (1) | - | Yes |
| ***INSR*** | 19:7112266-7294045 | - | - | 0 (1) | Yes |
| ***MAP2K7*** | 19:7968728-7979363 | - | 0 (1) | - | No |
| ***NDUFA7*** | 19:8373490-8386280 | - | 46916 (6) | - | No |
| ***RPS28*** | 19:8386042-8388224 | - | 44972 (5) | - | No |
| ***ANGPTL4*** | 19:8428173-8439257 | - | 2522 (1) | - | Yes |
| ***ADAMTS10*** | 19:8645126-8675620 | - | -34232 (3) | -34232 (3) | No |
| ***TMED1*** | 19:10943114-10946994 | 44833 (4) | - | - | No |
| ***CARM1*** | 19:10982189-11033453 | 0 (1) | - | - | No |
| ***SMARCA4*** | 19:11071598-11176071 | 0 (1) | - | - | No |
| ***LDLR*** | 19:11200038-11244492 | 0 (1) | - | - | Yes |
| ***NCAN*** | 19:19322782-19363042 | 20713 (4) | - | 20713 (4) | No |
| ***HAPLN4*** | 19:19366450-19373605 | 10150 (3) | - | 10150 (3) | No |
| ***TSSK6*** | 19:19623227-19626838 | 30794 (6) | - | 30794 (6) | No |
| ***NDUFA13*** | 19:19626545-19644285 | 13347 (4) | - | 13347 (4) | No |
| ***CILP2*** | 19:19649057-19657468 | 164 (1) | - | 164 (1) | Yes |
| ***LPAR2*** | 19:19734477-19739739 | -12501 (2) | - | -12501 (2) | No |
| ***PEPD*** | 19:33877856-34012700 | - | 0 (1) | 0 (1) | Yes |
| ***SCN1B*** | 19:35521588-35531352 | - | - | 25392 (2) | No |
| ***HPN*** | 19:35531410-35557475 | - | - | 0 (1) | No |
| ***PVR*** | 19:45147098-45166850 | 28463 (3) | - | - | No |
| ***CEACAM16*** | 19:45202421-45213986 | -7108 (1) | 33641 (3) | 33641 (3) | No |
| ***BCAM*** | 19:45312328-45324673 | 4541 (1) | 4541 (1) | 46165 (4) | No |
| ***PVRL2*** | 19:45349432-45392485 | 3134 (2) | 3134 (2) | 3134 (2) | No |
| ***APOE*** | 19:45409011-45412650 | -13392 (3) | -13392 (3) | -13392 (3) | Yes |
| ***APOC1*** | 19:45417504-45422606 | -21885 (4) | -21885 (4) | -21885 (4) | No |
| ***APOC4-APOC2*** | 19:45445495-45452822 | -49876 (5) | -49876 (5) | -49876 (5) | No |
| ***APOC2*** | 19:45449243-45452822 | -47577 (7) | -49899 (7) | 37606 (5) | No |
| ***MARK4*** | 19:45582546-45808541 | 0 (2) | - | - | No |
| ***GIPR*** | 19:46171502-46186982 | 20828 (4) | - | - | No |
| ***DMPK*** | 19:46272975-46285810 | -31643 (4) | - | - | No |
| ***SAE1*** | 19:47616531-47713886 | - | -26636 (2) | -26636 (2) | No |
| ***FGF21*** | 19:49258816-49261587 | -44542 (7) | - | - | No |
| ***BCAT2*** | 19:49298319-49314286 | -48080 (7) | - | - | No |
| ***FLT3LG*** | 19:49977464-49989488 | - | - | 38675 (7) | No |
| ***RPL13A*** | 19:49990811-49995565 | - | - | 32598 (6) | No |
| ***RPS11*** | 19:49999622-50002946 | - | - | 25217 (4) | No |
| ***FCGRT*** | 19:50010073-50029590 | - | - | 0 (1) | No |
| ***FPR1*** | 19:52248425-52307363 | - | 15725 (2) | - | No |
| ***FPR2*** | 19:52255279-52273779 | - | 49309 (4) | - | No |
| ***FPR3*** | 19:52298416-52329442 | - | 0 (1) | - | No |
| ***RPS9*** | 19:54704610-54752862 | - | 39907 (5) | - | No |
| ***LILRA6*** | 19:54720737-54746649 | - | 46120 (6) | - | No |
| ***LILRB5*** | 19:54754263-54761164 | - | 31605 (4) | - | No |
| ***LILRB2*** | 19:54777675-54785039 | - | 42441 (5) | - | No |
| ***LILRA3*** | 19:54799854-54809952 | - | 17528 (3) | - | Yes |
| ***LILRA5*** | 19:54818353-54824409 | - | 3071 (1) | - | No |
| ***LILRA4*** | 19:54844456-54850421 | - | -16976 (2) | - | No |
| ***LAIR1*** | 19:54865362-54882165 | - | -37882 (4) | - | No |
| ***GGT7*** | 20:33432523-33460663 | - | -23173 (2) | - | No |
| ***GSS*** | 20:33516236-33543620 | - | 0 (1) | 0 (1) | No |
| ***MYH7B*** | 20:33563206-33590240 | - | -33440 (3) | -37799 (3) | No |
| ***EDEM2*** | 20:33703167-33865928 | - | 0 (1) | - | No |
| ***PROCR*** | 20:33759876-33765165 | - | 12818 (2) | - | No |
| ***MMP24*** | 20:33814457-33864801 | - | -36474 (3) | - | No |
| ***GDF5*** | 20:34021145-34042568 | 3933 (2) | - | - | No |
| ***TOP1*** | 20:39657458-39753127 | 0 (1) | - | - | Yes |
| ***EMILIN3*** | 20:39988606-39995467 | -20655 (2) | - | - | No |
| ***HNF4A*** | 20:42984340-43061485 | 0 (1) | 0 (1) | - | Yes |
| ***TNNC2*** | 20:44451853-44462384 | - | -24435 (4) | -24435 (4) | No |
| ***CTSA*** | 20:44518783-44527459 | - | 4796 (2) | 20734 (3) | No |
| ***PLTP*** | 20:44527399-44540794 | - | 0 (1) | 7399 (1) | Yes |
| ***MMP9*** | 20:44637547-44645200 | - | 0 (1) | 0 (1) | No |
| ***SLC12A5*** | 20:44650356-44688784 | - | -5391 (2) | -5391 (2) | No |
| ***CD40*** | 20:44746911-44758502 | - | -12540 (1) | - | No |
| ***PPIL2*** | 22:22006559-22054304 | - | -23667 (5) | - | No |
| ***PLA2G6*** | 22:38507502-38601697 | - | - | 0 (1) | Yes |
| ***KCNJ4*** | 22:38822332-38851205 | - | - | 46846 (4) | No |
| ***PPARA*** | 22:46546424-46639653 | 0 (1) | - | - | Yes |

**Table S3. Univariable drug target MR estimates.** * - significant in the discovery analysis; †- significant in both original and validation study and concordant direction of effect. OR = odds ratio per 1-SD increase in LDL-C/HDL-C or triglycerides; CI = confidence interval.

| Druggable gene | Genomic coordinates | LDL-C (OR, 95% CI) | HDL-C (OR, 95% CI) | Triglycerides (OR, 95% CI) |
|---|---|---|---|---|
| *ABCA1* | chr9:107543283-107690518 | 2.05 (1.34, 3.15)* | 1.41 (0.66, 3.0) | 2.4 (1.29, 4.49)* |
| *ABCB11* | chr2:169779448-169887832 | 1.51 (0.7, 3.25) | - | - |
| *ABHD16A* | chr6:31654726-31671221 | 0.7 (0.14, 3.54) | 1.06 (0.21, 5.4) | 0.94 (0.33, 2.68) |
| *ABHD6* | chr3:58223233-58281420 | 2.25 (0.87, 5.87) | - | - |
| *ACP2* | chr11:47260853-47270457 | - | 1.1 (1.0, 1.2)* | 0.86 (0.58, 1.26) |
| *ADAM10* | chr15:58887403-59042177 | - | 1.87 (0.94, 3.75) | - |
| *ADAMTS10* | chr19:8645126-8675620 | - | 0.43 (0.16, 1.17) | 3.1 (1.21, 7.98)* |
| *ADAMTS13* | chr9:136279478-136324508 | 11.18 (4.37, 28.59)*† | - | - |
| *ADH4* | chr4:100044808-100078949 | - | 1.05 (0.39, 2.85) | - |
| *ADH5* | chr4:99992132-100009952 | - | 1.05 (0.39, 2.85) | - |
| *ADRB1* | chr10:115803806-115806667 | - | 1.67 (0.58, 4.8) | - |
| *AGER* | chr6:32148745-32152101 | 1.47 (0.71, 3.06) | 1.82 (1.08, 3.04)* | 1.07 (0.15, 7.64) |
| *AGRP* | chr16:67516474-67517716 | - | 0.98 (0.66, 1.45) | - |
| *AKR1C3* | chr10:5077546-5149878 | - | - | 1.05 (0.49, 2.25) |
| *AKT1* | chr14:105235686-105262088 | - | 0.49 (0.18, 1.36) | - |
| *ALDH1A2* | chr15:58245622-58790065 | - | 0.89 (0.81, 0.99)* | 1.28 (1.07, 1.54)*† |
| *ALDH2* | chr12:112204691-112247782 | 0.14 (0.07, 0.29)* | - | - |
| *ALOX5* | chr10:45869661-45941561 | - | 1.74 (1.18, 2.58)* | - |
| *AMPD2* | chr1:110158726-110174673 | 2.51 (1.53, 4.11)* | 2.97 (0.88, 10.06) | - |
| *AMPD3* | chr11:10329860-10529126 | - | 0.5 (0.27, 0.92)* | - |
| *ANGPTL3* | chr1:63063158-63071830 | 1.21 (1.11, 1.33)* | 1.61 (0.52, 5.01) | 1.16 (1.08, 1.25)* |
| *ANGPTL4* | chr19:8428173-8439257 | - | 0.48 (0.28, 0.83)*† | 3.38 (1.02, 11.22)*† |
| *AOC1* | chr7:150521715-150558592 | - | 0.81 (0.46, 1.41) | - |
| *APOA1* | chr11:116706467-116708666 | 1.88 (1.49, 2.36)*† | 0.84 (0.63, 1.11) | 1.25 (1.12, 1.4)*† |
| *APOA4* | chr11:116691419-116694022 | 1.51 (1.23, 1.86)*† | 0.53 (0.38, 0.74)* | 1.27 (1.14, 1.43)*† |
| *APOA5* | chr11:116660083-116663136 | 2.05 (1.4, 3.02)*† | 0.72 (0.6, 0.87)*† | 1.21 (1.12, 1.31)*† |
| *APOB* | chr2:21224301-21266945 | 1.5 (1.18, 1.9)*† | 1.23 (0.72, 2.12) | 0.53 (0.29, 0.98)*† |
| *APOC1* | chr19:45417504-45422606 | 1.31 (1.22, 1.41)*† | 0.39 (0.25, 0.59)* | 0.51 (0.17, 1.47) |
| *APOC2* | chr19:45449243-45452822 | 1.2 (0.87, 1.66) | 0.55 (0.26, 1.14) | 1.29 (0.31, 5.39) |
| *APOC3* | chr11:116700422-116703788 | 2.04 (1.72, 2.42)*† | 0.67 (0.58, 0.78)* | 1.26 (1.12, 1.41)*† |
| *APOC4-APOC2* | chr19:45445495-45452822 | 1.18 (0.86, 1.63) | 0.54 (0.26, 1.12) | 1.66 (0.9, 3.07) |
| *APOE* | chr19:45409011-45412650 | 1.3 (1.2, 1.41)*† | 0.39 (0.26, 0.59)* | 0.5 (0.17, 1.45) |
| *APOH* | chr17:64208151-64252643 | 1.52 (0.76, 3.02) | 0.66 (0.29, 1.53) | - |
| *APOM* | chr6:31620193-31625987 | 2.32 (0.82, 6.58) | 1.06 (0.21, 5.4) | 0.96 (0.36, 2.6) |
| *ASGR1* | chr17:7076750-7082883 | 1.39 (0.55, 3.49) | - | - |
| *ATG4C* | chr1:63249806-63331184 | 0.64 (0.31, 1.33) | - | 0.94 (0.55, 1.61) |
| *ATRAID* | chr2:27434895-27440046 | - | - | 0.85 (0.71, 1.02) |
| *AURKB* | chr17:8108056-8113918 | 1.35 (0.64, 2.84) | - | - |
| *BACE1* | chr11:117156402-117186975 | - | 2.72 (1.68, 4.39)* | 0.82 (0.04, 16.33) |
| *BAZ2A* | chr12:56989380-57030600 | - | - | 0.56 (0.2, 1.63) |
| *BCAM* | chr19:45312328-45324673 | 1.09 (0.71, 1.69) | 0.4 (0.18, 0.87)* | 0.63 (0.4, 0.97)* |
| *BCAT2* | chr19:49298319-49314286 | 0.94 (0.48, 1.83) | - | - |
| *BLK* | chr8:11351510-11422113 | - | - | 0.46 (0.31, 0.7)* |
| *BMP8A* | chr1:39957318-39991607 | - | 0.52 (0.4, 0.69)* | 1.8 (0.6, 5.46) |
| *C2* | chr6:31865562-31913449 | 1.62 (0.93, 2.8) | 1.85 (0.66, 5.17) | 0.21 (0.07, 0.6)* |
| *C4A* | chr6:31949801-31970458 | 2.26 (0.88, 5.85) | 1.04 (0.21, 5.12) | 0.22 (0.08, 0.65)* |
| *C4B* | chr6:31982539-32003195 | 2.41 (1.6, 3.63)* | 1.23 (0.33, 4.63) | 2.28 (1.11, 4.68)* |
| *C6orf15* | chr6:31079000-31080336 | 1.62 (1.23, 2.14)* | 1.08 (0.43, 2.72) | 1.5 (1.0, 2.25)* |
| *C6orf25* | chr6:31686371-31694491 | 0.7 (0.14, 3.54) | - | 1.26 (0.43, 3.69) |
| *C9orf96* | chr9:136243117-136271220 | 5.77 (2.71, 12.31)*† | - | - |
| *CACNB1* | chr17:37329709-37353956 | - | 0.38 (0.2, 0.72)* | - |
| *CAD* | chr2:27440258-27466811 | - | - | 1.01 (0.85, 1.19) |
| *CAMKV* | chr3:49895421-49907655 | - | 0.18 (0.1, 0.31)* | - |
| *CARM1* | chr19:10982189-11033453 | 2.27 (1.68, 3.05)*† | - | - |
| *CATSPER2* | chr15:43920701-43960316 | - | - | 1.05 (0.46, 2.37) |
| *CBLN3* | chr14:24895738-24900160 | 1.25 (0.61, 2.54) | - | - |
| *CCL22* | chr16:57392684-57400102 | - | 0.37 (0.11, 1.22) | - |

| Druggable gene | Genomic coordinates | LDL-C (OR, 95% CI) | HDL-C (OR, 95% CI) | Triglycerides (OR, 95% CI) |
|---|---|---|---|---|
| ***CD300LG*** | chr17:41924516-41940997 | - | 1.04 (0.57, 1.91) | 1.58 (0.58, 4.35) |
| ***CD40*** | chr20:44746911-44758502 | - | 1.32 (0.46, 3.77) | 0.75 (0.2, 2.72) |
| ***CDK12*** | chr17:37617764-37721160 | - | 0.19 (0.0, 37.54) | - |
| ***CEACAM16*** | chr19:45202421-45213986 | 1.66 (1.31, 2.11)*† | 0.46 (0.27, 0.79)* | 0.56 (0.25, 1.27) |
| ***CELSR2*** | chr1:109792641-109818372 | 1.97 (1.78, 2.18)*† | 0.06 (0.04, 0.09)* | - |
| ***CES3*** | chr16:66995140-67009051 | - | 1.15 (0.6, 2.18) | - |
| ***CES4A*** | chr16:67022492-67043661 | - | 1.15 (0.6, 2.18) | - |
| ***CETP*** | chr16:56995762-57017757 | 1.49 (1.29, 1.72)* | 0.91 (0.87, 0.95)*† | 1.98 (1.63, 2.4)*† |
| ***CFB*** | chr6:31895475-31919861 | 1.61 (0.94, 2.77) | 1.85 (0.66, 5.16) | 0.46 (0.24, 0.91)* |
| ***CGREF1*** | chr2:27321757-27341995 | - | - | 0.94 (0.66, 1.36) |
| ***CHST1*** | chr11:45670427-45687172 | - | 1.15 (0.45, 2.94) | - |
| ***CILP2*** | chr19:19649057-19657468 | 1.19 (1.01, 1.39)* | - | 1.18 (1.0, 1.39)*† |
| ***CPS1*** | chr2:211342406-211543831 | - | 2.05 (1.1, 3.82)* | - |
| ***CRY2*** | chr11:45868669-45904798 | - | 0.71 (0.48, 1.04) | - |
| ***CSF1*** | chr1:110452864-110473614 | - | 0.57 (0.28, 1.15) | - |
| ***CSF3*** | chr17:38171614-38174066 | 0.3 (0.12, 0.74)* | 0.87 (0.4, 1.89) | - |
| ***CSNK1G3*** | chr5:122847793-122952739 | 0.33 (0.2, 0.55)* | - | - |
| ***CTRL*** | chr16:67961543-67966317 | - | 1.11 (0.91, 1.35) | - |
| ***CTSA*** | chr20:44518783-44527459 | - | 1.89 (1.3, 2.75)* | 0.12 (0.05, 0.28)* |
| ***CTSB*** | chr8:11700033-11726957 | - | - | 0.65 (0.44, 0.98)* |
| ***CYP21A2*** | chr6:32006042-32009447 | 2.34 (1.49, 3.66)* | 1.23 (0.33, 4.63) | 2.22 (1.03, 4.81)* |
| ***CYP26A1*** | chr10:94833232-94837647 | 7.25 (4.25, 12.37)* | 0.22 (0.09, 0.51)* | 4.35 (2.79, 6.79)* |
| ***CYP26C1*** | chr10:94821021-94828454 | 7.25 (4.23, 12.42)* | 0.22 (0.09, 0.51)* | 4.36 (2.79, 6.83)* |
| ***CYP7A1*** | chr8:59402737-59412795 | 0.95 (0.47, 1.91) | - | 0.86 (0.33, 2.28) |
| ***DAGLA*** | chr11:61447905-61514473 | 1.67 (1.21, 2.29)* | 0.49 (0.1, 2.47) | 0.63 (0.53, 0.75)* |
| ***DAGLB*** | chr7:6448757-6523821 | - | 0.35 (0.22, 0.54)* | - |
| ***DCPS*** | chr11:126173647-126215644 | 4.96 (1.89, 13.06)* | 0.29 (0.13, 0.66)* | - |
| ***DDR1*** | chr6:30844198-30867933 | 0.95 (0.5, 1.8) | 1.62 (0.42, 6.27) | 0.99 (0.37, 2.67) |
| ***DHODH*** | chr16:72042487-72058954 | 0.66 (0.44, 1.0) | - | 7.42 (2.32, 23.71)* |
| ***DMPK*** | chr19:46272975-46285810 | 2.78 (1.64, 4.73)* | 0.4 (0.11, 1.45) | - |
| ***DPEP2*** | chr16:68021297-68034489 | - | 1.36 (0.98, 1.89) | - |
| ***DPEP3*** | chr16:68009566-68014732 | - | 1.36 (0.98, 1.9) | - |
| ***DUSP3*** | chr17:41843489-41856356 | - | 0.93 (0.25, 3.55) | 1.03 (0.39, 2.69) |
| ***EDEM2*** | chr20:33703167-33865928 | 2.37 (0.76, 7.37) | 0.0 (0.0, 0.0)* | - |
| ***EGFL8*** | chr6:32132360-32136058 | 1.47 (0.71, 3.06) | - | 1.96 (0.97, 3.97) |
| ***EHMT2*** | chr6:31847536-31865464 | 3.32 (1.25, 8.83)* | 2.66 (0.73, 9.69) | 0.23 (0.03, 1.56) |
| ***EMILIN1*** | chr2:27301435-27309271 | - | - | 0.73 (0.34, 1.57) |
| ***EMILIN3*** | chr20:39988606-39995467 | 2.51 (1.29, 4.86)* | - | - |
| ***ERBB2*** | chr17:37844167-37886679 | - | 2.82 (0.25, 31.53) | - |
| ***ERP29*** | chr12:112451120-112461255 | 0.11 (0.06, 0.18)* | 0.04 (0.02, 0.13)* | - |
| ***ESR1*** | chr6:151977826-152450754 | - | 2.11 (1.13, 3.93)* | - |
| ***F2*** | chr11:46740730-46761056 | 0.17 (0.05, 0.59)* | 0.57 (0.13, 2.43) | 0.35 (0.13, 0.94)* |
| ***FCGRT*** | chr19:50010073-50029590 | - | - | 1.95 (0.75, 5.07) |
| ***FDFT1*** | chr8:11653082-11696818 | - | - | 0.88 (0.44, 1.73) |
| ***FEN1*** | chr11:61560109-61564716 | 2.02 (0.99, 4.12) | 0.54 (0.2, 1.47) | 1.13 (0.71, 1.8) |
| ***FGF21*** | chr19:49258816-49261587 | 1.06 (0.79, 1.43) | - | 1.62 (0.46, 5.78) |
| ***FKBP6*** | chr7:72742167-72772634 | - | - | 0.52 (0.2, 1.34) |
| ***FLT3LG*** | chr19:49977464-49989488 | - | - | 1.95 (0.75, 5.07) |
| ***FN1*** | chr2:216225163-216300895 | 0.04 (0.01, 0.22)* | - | - |
| ***FOLH1*** | chr11:49168187-49230222 | - | 0.76 (0.36, 1.61) | - |
| ***FPR1*** | chr19:52248425-52307363 | - | 1.58 (1.11, 2.24)* | - |
| ***FPR2*** | chr19:52255279-52273779 | - | 1.89 (1.24, 2.87)* | - |
| ***FPR3*** | chr19:52298416-52329442 | - | 1.58 (1.1, 2.26)* | - |
| ***FRK*** | chr6:116252312-116381921 | 0.76 (0.48, 1.21) | 0.57 (0.17, 1.94) | - |
| ***FZD9*** | chr7:72848109-72850450 | - | 1.13 (0.52, 2.46) | 1.08 (0.72, 1.61) |
| ***GALK1*** | chr17:73747675-73761792 | 3.38 (1.46, 7.85)* | - | - |
| ***GALNT2*** | chr1:230193536-230417870 | - | 0.56 (0.42, 0.74)* | 2.19 (1.48, 3.25)* |
| ***GANC*** | chr15:42565431-42645864 | - | 0.72 (0.31, 1.67) | 1.32 (0.65, 2.71) |
| ***GCKR*** | chr2:27719709-27746554 | 1.49 (0.88, 2.51) | - | 0.9 (0.45, 1.83) |
| ***GDF5*** | chr20:34021145-34042568 | 2.71 (0.77, 9.5) | - | - |
| ***GDF7*** | chr2:20866424-20873418 | 0.96 (0.6, 1.53) | - | 1.33 (0.63, 2.8) |
| ***GFOD2*** | chr16:67708434-67753324 | - | 1.42 (1.01, 2.0)* | - |
| ***GGT7*** | chr20:33432523-33460663 | - | 0.43 (0.15, 1.23) | 2.61 (1.1, 6.18)* |
| ***GIPR*** | chr19:46171502-46186982 | 2.72 (1.06, 7.02)* | 0.89 (0.37, 2.12) | 4.17 (1.16, 15.02)* |
| ***GPER1*** | chr7:1121844-1133451 | 2.81 (0.85, 9.29) | 2.12 (0.84, 5.34) | - |

| Druggable gene | Genomic coordinates | LDL-C (OR, 95% CI) | HDL-C (OR, 95% CI) | Triglycerides (OR, 95% CI) |
|---|---|---|---|---|
| ***GPIHBP1*** | chr8:144295068-144299044 | - | 1.82 (1.23, 2.71)* | - |
| ***GPR146*** | chr7:1084212-1098897 | 2.81 (0.85, 9.29) | 2.12 (0.85, 5.33) | - |
| ***GPR61*** | chr1:110082494-110091028 | 1.97 (1.56, 2.5)*† | 3.02 (0.77, 11.91) | 5.14 (1.43, 18.48)* |
| ***GSK3B*** | chr3:119540170-119813264 | - | 0.45 (0.16, 1.25) | - |
| ***GSK3B*** | chr3:119540170-119813264 | - | 0.45 (0.16, 1.25) | - |
| ***GSS*** | chr20:33516236-33543620 | - | 0.57 (0.2, 1.65) | 2.25 (1.05, 4.85)* |
| ***GSTM1*** | chr1:110230436-110251661 | 3.28 (1.83, 5.87)* | - | - |
| ***GSTM2*** | chr1:110210644-110252171 | 3.28 (1.83, 5.89)* | - | - |
| ***GSTM3*** | chr1:110276554-110284384 | 0.06 (0.0, 0.79)* | - | - |
| ***GSTM4*** | chr1:110198703-110208118 | 3.41 (1.54, 7.53)* | 2.97 (0.88, 10.06) | - |
| ***GSTM5*** | chr1:110254864-110318050 | 0.06 (0.0, 0.84)* | - | - |
| ***H3F3B*** | chr17:73772515-73781974 | 3.38 (1.46, 7.85)* | - | - |
| ***HAPLN4*** | chr19:19366450-19373605 | 1.06 (0.91, 1.22) | - | 1.07 (0.93, 1.25) |
| ***HCAR1*** | chr12:123104824-123215390 | - | 0.99 (0.68, 1.44) | - |
| ***HCAR2*** | chr12:123185840-123187890 | - | 0.85 (0.52, 1.38) | - |
| ***HCAR3*** | chr12:123199303-123201439 | - | 0.85 (0.52, 1.39) | - |
| ***HDGF*** | chr1:156711899-156736717 | - | 1.18 (0.44, 3.19) | - |
| ***HFE*** | chr6:26087509-26098571 | 1.92 (0.77, 4.83) | 0.81 (0.18, 3.61) | - |
| ***HGFAC*** | chr4:3443614-3451211 | 2.07 (1.35, 3.16)* | - | 1.97 (1.34, 2.89)* |
| ***HIST1H1C*** | chr6:26055968-26056699 | 1.83 (0.79, 4.25) | 0.84 (0.18, 4.01) | - |
| ***HIST1H4C*** | chr6:26104104-26104518 | 1.92 (0.77, 4.83) | 0.81 (0.18, 3.61) | - |
| ***HLA-B*** | chr6:31321649-31324965 | 0.31 (0.08, 1.16) | 1.81 (0.84, 3.88) | 1.8 (1.46, 2.22)* |
| ***HLA-C*** | chr6:31236526-31239907 | 1.47 (0.06, 35.1) | 1.73 (0.91, 3.28) | 1.06 (0.8, 1.4) |
| ***HLA-DOB*** | chr6:32780540-32784825 | 3.57 (2.2, 5.78)* | 1.06 (0.31, 3.59) | 1.31 (0.81, 2.11) |
| ***HLA-DQA2*** | chr6:32709119-32714992 | 2.95 (2.41, 3.6)* | 2.23 (1.19, 4.19)* | 1.17 (0.82, 1.68) |
| ***HLA-DRA*** | chr6:32407619-32412823 | 2.29 (1.55, 3.37)* | 1.2 (0.78, 1.85) | 2.34 (1.41, 3.86)* |
| ***HLA-DRB1*** | chr6:32546546-32557625 | 1.37 (0.97, 1.93) | 0.92 (0.6, 1.41) | 1.44 (0.63, 3.32) |
| ***HLA-DRB5*** | chr6:32485120-32498064 | 2.75 (1.13, 6.7)* | 0.8 (0.12, 5.14) | - |
| ***HLA-G*** | chr6:29794744-29798902 | 1.39 (0.44, 4.38) | - | 1.33 (0.41, 4.3) |
| ***HMGCR*** | chr5:74632154-74657929 | 1.22 (1.03, 1.45)* | - | - |
| ***HNF4A*** | chr20:42984340-43061485 | 1.51 (0.67, 3.38) | 1.18 (0.87, 1.6) | - |
| ***HP*** | chr16:72088491-72094954 | 1.1 (0.88, 1.38) | - | 7.42 (2.32, 23.71)* |
| ***HPD*** | chr12:122277433-122301502 | - | 1.81 (0.68, 4.83) | 0.38 (0.12, 1.16) |
| ***HPN*** | chr19:35531410-35557475 | - | - | 0.61 (0.24, 1.51) |
| ***HSD11B2*** | chr16:67464555-67471456 | - | 0.58 (0.31, 1.1) | - |
| ***HSD17B11*** | chr4:88257762-88312538 | - | - | 1.36 (0.9, 2.05) |
| ***HSD3B7*** | chr16:30996519-31000473 | - | - | 1.32 (0.63, 2.78) |
| ***HSPA1A*** | chr6:31783291-31785723 | 10.64 (2.34, 48.34)* | - | 1.5 (0.44, 5.09) |
| ***HSPA1B*** | chr6:31795512-31798031 | 4.6 (1.61, 13.1)* | - | 0.02 (0.0, 2.7) |
| ***IGF2R*** | chr6:160390131-160534539 | 4.56 (2.73, 7.61)* | - | 224064.11 (5.34, 9394669504.1)* |
| ***INHBC*** | chr12:57828543-57844611 | - | 0.64 (0.41, 1.0)* | 1.95 (0.99, 3.84) |
| ***INHBE*** | chr12:57846106-57853063 | - | 0.58 (0.35, 0.97)* | 1.95 (1.0, 3.81)* |
| ***INSR*** | chr19:7112266-7294045 | - | 0.69 (0.47, 1.04) | 1.15 (0.83, 1.62) |
| ***ITGB3*** | chr17:45331212-45421658 | 1.64 (1.06, 2.52)* | 2.79 (0.81, 9.62) | - |
| ***ITGB4*** | chr17:73717408-73753899 | 3.38 (1.46, 7.85)* | - | - |
| ***ITIH1*** | chr3:52811603-52826078 | 2.07 (0.59, 7.23) | 3.11 (0.85, 11.31) | - |
| ***ITIH3*** | chr3:52828784-52843025 | 2.07 (0.59, 7.23) | 2.75 (0.81, 9.38) | - |
| ***ITIH4*** | chr3:52846991-52865495 | 2.07 (0.59, 7.23) | 2.75 (0.81, 9.38) | - |
| ***KCNJ4*** | chr22:38822332-38851205 | - | - | 0.45 (0.16, 1.3) |
| ***KCNK16*** | chr6:39282474-39290744 | 1.13 (0.42, 3.04) | - | - |
| ***KCNK17*** | chr6:39266777-39282329 | 2.89 (1.46, 5.74)* | - | - |
| ***KCNK7*** | chr11:65360326-65363467 | - | 0.09 (0.03, 0.3)* | 16.79 (4.99, 56.51)* |
| ***KHK*** | chr2:27309615-27323640 | - | - | 0.74 (0.41, 1.34) |
| ***KLHL8*** | chr4:88081255-88161466 | 0.96 (0.31, 2.93) | 0.98 (0.47, 2.05) | 1.15 (0.95, 1.41) |
| ***L3MBTL3*** | chr6:130334844-130462594 | - | - | 1.54 (0.57, 4.14) |
| ***LACTB*** | chr15:63413999-63434260 | - | 0.64 (0.48, 0.86)* | 1.43 (0.87, 2.36) |
| ***LAIR1*** | chr19:54865362-54882165 | - | 1.53 (0.87, 2.69) | - |
| ***LCAT*** | chr16:67973653-67978034 | - | 1.11 (0.92, 1.34) | - |
| ***LCT*** | chr2:136545410-136594750 | 0.61 (0.23, 1.58) | - | - |
| ***LDLR*** | chr19:11200038-11244492 | 1.37 (0.98, 1.93) | 0.04 (0.01, 0.34)* | - |
| ***LGR4*** | chr11:27387508-27494322 | - | - | 4.07 (2.03, 8.16)* |
| ***LILRA3*** | chr19:54799854-54809952 | - | 0.76 (0.44, 1.33) | - |
| ***LILRA4*** | chr19:54844456-54850421 | - | 0.87 (0.44, 1.69) | - |
| ***LILRA5*** | chr19:54818353-54824409 | - | 0.8 (0.45, 1.43) | - |

| Druggable gene | Genomic coordinates | LDL-C (OR, 95% CI) | HDL-C (OR, 95% CI) | Triglycerides (OR, 95% CI) |
|---|---|---|---|---|
| ***LILRA6*** | chr19:54720737-54746649 | - | 0.96 (0.65, 1.41) | - |
| ***LILRB2*** | chr19:54777675-54785039 | - | 0.51 (0.27, 0.96)* | - |
| ***LILRB5*** | chr19:54754263-54761164 | - | 0.96 (0.65, 1.41) | - |
| ***LIPC*** | chr15:58702768-58861151 | - | 0.99 (0.89, 1.11) | 0.91 (0.44, 1.86) |
| ***LIPG*** | chr18:47087069-47119272 | 0.41 (0.25, 0.67)* | 0.95 (0.82, 1.1) | - |
| ***LPA*** | chr6:160952515-161087407 | 13.5 (7.17, 25.42)* | 21.73 (4.69, 100.69)* | 3.77 (1.78, 7.99)* |
| ***LPAR2*** | chr19:19734477-19739739 | 1.48 (1.22, 1.8)* | - | 1.62 (1.39, 1.9)* |
| ***LPL*** | chr8:19759228-19824769 | - | 0.63 (0.49, 0.82)*† | 1.68 (1.46, 1.92)*† |
| ***LRPAP1*** | chr4:3508103-3534286 | 5.03 (1.74, 14.56)* | - | 2.59 (1.6, 4.2)* |
| ***LTA*** | chr6:31539831-31542101 | 2.03 (1.04, 3.97)* | - | 1.22 (0.78, 1.9) |
| ***LTA*** | chr6:31539831-31542101 | 2.03 (1.04, 3.97)* | - | 1.22 (0.78, 1.9) |
| ***LTB*** | chr6:31548302-31550299 | 2.01 (1.03, 3.93)* | 1.2 (0.44, 3.25) | 1.11 (0.76, 1.62) |
| ***LTK*** | chr15:41795836-41806085 | - | 0.8 (0.49, 1.29) | - |
| ***MAP2K7*** | chr19:7968728-7979363 | - | 1.24 (0.51, 3.02) | 0.73 (0.2, 2.71) |
| ***MAP3K11*** | chr11:65365226-65382853 | - | 0.09 (0.03, 0.3)* | 16.79 (4.99, 56.51)* |
| ***MAP3K19*** | chr2:135722061-135805038 | 1.21 (0.53, 2.8) | - | - |
| ***MAP3K4*** | chr6:161412759-161538417 | 2.89 (1.75, 4.78)* | - | - |
| ***MAPK10*** | chr4:86936276-87515284 | - | 0.75 (0.47, 1.19) | 1.22 (0.53, 2.77) |
| ***MAPKAPK5*** | chr12:112279782-112334343 | 0.26 (0.12, 0.57)* | - | - |
| ***MARK4*** | chr19:45582546-45808541 | 1.02 (0.73, 1.41) | 0.72 (0.18, 2.87) | - |
| ***MARS*** | chr12:57869228-57911352 | - | 0.63 (0.39, 1.03) | 1.95 (1.0, 3.83) |
| ***MAS1L*** | chr6:29454474-29455738 | - | - | 1.32 (0.62, 2.83) |
| ***MET*** | chr7:116312444-116438440 | - | 0.66 (0.43, 1.01) | 1.09 (0.42, 2.87) |
| ***METAP1*** | chr4:99916771-99983964 | - | 1.05 (0.39, 2.85) | - |
| ***MFAP1*** | chr15:44096690-44117000 | - | 0.72 (0.24, 2.17) | 1.33 (0.56, 3.19) |
| ***MMP24*** | chr20:33814457-33864801 | 2.37 (0.76, 7.37) | 1.56 (0.93, 2.63) | - |
| ***MMP9*** | chr20:44637547-44645200 | - | 1.07 (0.14, 8.55) | 0.46 (0.24, 0.85)* |
| ***MOGAT2*** | chr11:75428864-75444003 | - | 0.96 (0.43, 2.14) | - |
| ***MST1R*** | chr3:49924435-49941299 | - | 0.18 (0.11, 0.31)* | - |
| ***MYH7B*** | chr20:33563206-33590240 | - | 0.5 (0.19, 1.3) | 2.25 (1.05, 4.85)* |
| ***MYL4*** | chr17:45277812-45301045 | 2.05 (0.77, 5.49) | - | - |
| ***NAT2*** | chr8:18248755-18258728 | 3.37 (2.07, 5.49)* | - | 3.33 (1.94, 5.71)* |
| ***NCAN*** | chr19:19322782-19363042 | 1.04 (0.9, 1.2) | - | 1.1 (0.86, 1.39) |
| ***NCR3*** | chr6:31556672-31560762 | 2.15 (1.09, 4.25)* | 1.3 (0.46, 3.71) | 1.03 (0.69, 1.55) |
| ***NDUFA13*** | chr19:19626545-19644285 | 1.63 (1.13, 2.35)* | - | 1.18 (1.0, 1.39)*† |
| ***NDUFA7*** | chr19:8373490-8386280 | - | 0.57 (0.3, 1.07) | - |
| ***NDUFS3*** | chr11:47586888-47606114 | - | 1.18 (0.83, 1.69) | - |
| ***NEK4*** | chr3:52744800-52804965 | 1.58 (0.48, 5.24) | 1.28 (0.43, 3.77) | - |
| ***NEU1*** | chr6:31825436-31830683 | 3.32 (1.24, 8.84)* | 2.66 (0.73, 9.69) | 0.18 (0.02, 1.59) |
| ***NFKB1*** | chr4:103422486-103538459 | - | 1.25 (0.5, 3.08) | - |
| ***NISCH*** | chr3:52489134-52527087 | - | 0.57 (0.35, 0.93)* | 1.16 (0.31, 4.34) |
| ***NOTCH4*** | chr6:32162620-32191844 | 1.47 (0.7, 3.07) | 1.9 (1.17, 3.1)* | 0.87 (0.67, 1.13) |
| ***NPC1L1*** | chr7:44552134-44580914 | 2.01 (1.48, 2.73)*† | - | 2.56 (0.75, 8.68) |
| ***NPEPPS*** | chr17:45600308-45700642 | 1.43 (0.67, 3.07) | 3.28 (0.96, 11.23) | 0.6 (0.22, 1.63) |
| ***NR0B2*** | chr1:27237980-27240457 | 1.38 (0.6, 3.18) | 0.67 (0.27, 1.64) | 1.91 (0.71, 5.17) |
| ***NR1H3*** | chr11:47269851-47290396 | - | 1.07 (0.97, 1.18) | 0.86 (0.58, 1.27) |
| ***NR1I2*** | chr3:119499331-119537332 | - | 0.43 (0.17, 1.08) | - |
| ***NRBP1*** | chr2:27650657-27665126 | - | - | 0.89 (0.72, 1.1) |
| ***OBP2B*** | chr9:136080664-136084630 | 0.47 (0.06, 3.46) | - | - |
| ***OR11A1*** | chr6:29393281-29424848 | - | - | 2.42 (0.53, 11.09) |
| ***OR2H1*** | chr6:29424958-29432105 | - | - | 2.24 (0.55, 9.16) |
| ***OR4A16*** | chr11:55110627-55111707 | - | 0.83 (0.36, 1.93) | - |
| ***OR4C16*** | chr11:55339604-55340536 | - | 0.76 (0.32, 1.84) | - |
| ***PBRM1*** | chr3:52579368-52719933 | 1.58 (0.48, 5.24) | 0.77 (0.41, 1.44) | 2.21 (0.66, 7.34) |
| ***PCSK7*** | chr11:117075053-117103241 | 1.62 (0.85, 3.11) | 0.78 (0.51, 1.21) | 2.57 (0.09, 73.1) |
| ***PCSK9*** | chr1:55505221-55530525 | 1.6 (1.45, 1.77)*† | - | - |
| ***PDE3A*** | chr12:20522179-20837315 | - | 0.79 (0.4, 1.58) | - |
| ***PDGFC*** | chr4:157681606-157892546 | - | 0.52 (0.19, 1.39) | - |
| ***PDIA3*** | chr15:44038590-44065477 | - | - | 1.33 (0.56, 3.19) |
| ***PEPD*** | chr19:33877856-34012700 | - | 0.36 (0.25, 0.51)* | 3.41 (1.92, 6.08)* |
| ***PIP4K2C*** | chr12:57984957-57997198 | - | 1.14 (0.27, 4.85) | - |
| ***PKM*** | chr15:72491370-72524164 | - | - | 1.54 (0.59, 4.03) |
| ***PLA2G15*** | chr16:68279207-68294961 | - | 1.45 (1.06, 1.97)* | - |
| ***PLA2G6*** | chr22:38507502-38601697 | - | 1.01 (0.57, 1.78) | 1.09 (0.4, 3.0) |
| ***PLG*** | chr6:161123270-161174347 | 18.35 (5.47, 61.6)* | 5.48 (0.07, 456.86) | 0.75 (0.18, 3.14) |

| Druggable gene | Genomic coordinates | LDL-C (OR, 95% CI) | HDL-C (OR, 95% CI) | Triglycerides (OR, 95% CI) |
|---|---|---|---|---|
| ***PLTP*** | chr20:44527399-44540794 | - | 0.67 (0.1, 4.53) | 0.25 (0.02, 2.61) |
| ***PNMT*** | chr17:37824234-37826728 | - | 0.64 (0.36, 1.15) | - |
| ***PPARA*** | chr22:46546424-46639653 | 3.77 (1.44, 9.85)* | - | - |
| ***PPARG*** | chr3:12328867-12475855 | 1.67 (1.04, 2.68)* | 0.71 (0.35, 1.48) | 2.18 (1.14, 4.15)* |
| ***PPIL2*** | chr22:22006559-22054304 | - | 0.82 (0.47, 1.43) | - |
| ***PPY*** | chr17:42018172-42019836 | - | 1.34 (0.73, 2.47) | 0.48 (0.13, 1.75) |
| ***PROCR*** | chr20:33759876-33765165 | - | 0.0 (0.0, 0.0)* | - |
| ***PRSS36*** | chr16:31150246-31161415 | 1.52 (0.43, 5.39) | - | 1.16 (0.4, 3.32) |
| ***PRSS53*** | chr16:31094746-31100949 | 1.62 (0.45, 5.86) | - | 1.32 (0.52, 3.35) |
| ***PRSS8*** | chr16:31142756-31147083 | 1.52 (0.43, 5.39) | - | 1.32 (0.52, 3.36) |
| ***PSKH1*** | chr16:67927175-67963581 | - | 1.18 (0.92, 1.52) | - |
| ***PSMA1*** | chr11:14515329-14665181 | - | 1.43 (0.5, 4.11) | - |
| ***PSMA5*** | chr1:109941653-109969062 | 2.47 (1.8, 3.39)*† | 0.08 (0.02, 0.29)* | - |
| ***PSMB10*** | chr16:67968405-67970990 | - | 1.11 (0.91, 1.35) | - |
| ***PSMB8*** | chr6:32808494-32812480 | 2.23 (0.99, 4.98) | 1.06 (0.31, 3.59) | 1.09 (0.6, 1.97) |
| ***PSMC3*** | chr11:47440320-47447993 | - | 1.12 (0.9, 1.4) | 0.68 (0.18, 2.53) |
| ***PSMD3*** | chr17:38137050-38154213 | 0.3 (0.12, 0.74)* | 0.85 (0.55, 1.31) | - |
| ***PTPN11*** | chr12:112856155-112947717 | 1259419.6 (0.0, 4107788603997230.5) | 0.05 (0.02, 0.16)* | - |
| ***PTPN13*** | chr4:87515468-87736324 | - | 1.03 (0.64, 1.65) | 0.99 (0.7, 1.39) |
| ***PTPRJ*** | chr11:48002113-48189670 | - | 0.5 (0.35, 0.72)* | - |
| ***PVR*** | chr19:45147098-45166850 | 1.31 (1.12, 1.54)*† | 0.32 (0.11, 0.91)* | - |
| ***PVRL2*** | chr19:45349432-45392485 | 1.43 (1.27, 1.63)*† | 0.35 (0.24, 0.51)* | 0.51 (0.19, 1.37) |
| ***RAF1*** | chr3:12625100-12705725 | 2.06 (1.48, 2.86)* | - | 2.63 (0.79, 8.83) |
| ***RAF1*** | chr3:12625100-12705725 | 2.06 (1.48, 2.86)* | - | 2.63 (0.79, 8.83) |
| ***RELA*** | chr11:65421067-65430565 | - | 0.09 (0.03, 0.3)* | 16.79 (4.99, 56.51)* |
| ***RGS12*** | chr4:3294755-3441640 | 1.99 (1.31, 3.03)* | - | 1.98 (1.35, 2.89)* |
| ***RHCE*** | chr1:25688740-25756683 | 0.71 (0.4, 1.26) | - | - |
| ***RHD*** | chr1:25598884-25656936 | 0.27 (0.07, 1.07) | - | - |
| ***RPL13A*** | chr19:49990811-49995565 | - | - | 1.95 (0.75, 5.07) |
| ***RPL17*** | chr18:47014851-47018906 | 0.61 (0.21, 1.81) | 0.75 (0.52, 1.08) | - |
| ***RPL19*** | chr17:37356536-37360980 | - | 0.45 (0.23, 0.9)* | - |
| ***RPL5*** | chr1:93297582-93307481 | 0.43 (0.27, 0.68)* | - | - |
| ***RPL6*** | chr12:112842994-112856642 | 0.13 (0.06, 0.28)* | 0.05 (0.02, 0.16)* | - |
| ***RPL7A*** | chr9:136215069-136218281 | 2.29 (1.57, 3.36)*† | - | - |
| ***RPS11*** | chr19:49999622-50002946 | - | - | 1.95 (0.75, 5.07) |
| ***RPS28*** | chr19:8386042-8388224 | - | 0.57 (0.3, 1.07) | - |
| ***RPS6KA1*** | chr1:26856252-26901521 | - | 1.7 (0.62, 4.68) | - |
| ***RPS9*** | chr19:54704610-54752862 | - | 0.95 (0.64, 1.41) | - |
| ***RSPO3*** | chr6:127439749-127518910 | - | 0.69 (0.45, 1.07) | 1.1 (0.74, 1.65) |
| ***SAE1*** | chr19:47616531-47713886 | - | 0.23 (0.08, 0.65)* | 3.19 (1.82, 5.58)* |
| ***SCARB1*** | chr12:125261402-125367214 | 5.16 (1.85, 14.39)* | 0.42 (0.14, 1.26) | 8.0 (2.41, 26.56)* |
| ***SCN1B*** | chr19:35521588-35531352 | - | - | 0.61 (0.24, 1.51) |
| ***SCUBE3*** | chr6:35182190-35220856 | 0.2 (0.06, 0.66)* | 0.0 (0.0, 0.13)* | - |
| ***SEMA3F*** | chr3:50192478-50226508 | - | 0.4 (0.18, 0.89)* | - |
| ***SEMA3G*** | chr3:52467069-52479101 | - | 1.07 (0.45, 2.55) | 1.16 (0.31, 4.34) |
| ***SERPINA1*** | chr14:94843084-94857030 | 0.65 (0.22, 1.9) | - | - |
| ***SERPINA10*** | chr14:94749650-94759608 | 0.73 (0.26, 2.1) | - | - |
| ***SERPINA6*** | chr14:94770585-94789731 | 0.65 (0.22, 1.9) | - | - |
| ***SFN*** | chr1:27189633-27190947 | 1.51 (0.67, 3.41) | 0.49 (0.25, 0.97)* | 1.91 (0.71, 5.17) |
| ***SFTA2*** | chr6:30899130-30899952 | 0.85 (0.4, 1.78) | 1.97 (0.61, 6.41) | 0.69 (0.25, 1.9) |
| ***SIDT2*** | chr11:117049449-117068160 | 1.62 (0.85, 3.11) | 0.79 (0.51, 1.21) | 1.03 (0.19, 5.6) |
| ***SIK3*** | chr11:116714118-116969153 | 1.15 (0.57, 2.31) | 0.46 (0.29, 0.73)*† | 1.08 (0.98, 1.18) |
| ***SLC12A3*** | chr16:56899119-56949762 | 1.94 (1.43, 2.63)* | 0.89 (0.86, 0.93)*† | 0.75 (0.24, 2.33) |
| ***SLC12A4*** | chr16:67977377-68003504 | - | 1.11 (0.92, 1.33) | - |
| ***SLC12A5*** | chr20:44650356-44688784 | - | 2.06 (0.18, 22.96) | 0.4 (0.21, 0.76)* |
| ***SLC18A1*** | chr8:20002366-20040717 | - | 0.22 (0.08, 0.61)* | 2.41 (1.66, 3.5)* |
| ***SLC22A1*** | chr6:160542821-160579750 | 4.39 (2.62, 7.36)* | - | 224064.11 (5.16, 9728155751.75)* |
| ***SLC22A2*** | chr6:160592093-160698670 | 2.48 (1.85, 3.32)* | 2.55 (1.16, 5.63)* | 6.47 (2.56, 16.37)* |
| ***SLC22A3*** | chr6:160769300-160876014 | 5.13 (3.44, 7.66)* | 2.4 (1.27, 4.53)* | 4.42 (2.62, 7.45)* |
| ***SLC44A4*** | chr6:31830969-31846823 | 3.32 (1.25, 8.83)* | 2.66 (0.73, 9.69) | 0.23 (0.03, 1.59) |
| ***SLC5A6*** | chr2:27422455-27435826 | - | - | 0.86 (0.71, 1.05) |
| ***SLC9A1*** | chr1:27425306-27493472 | 1.03 (0.43, 2.46) | 0.96 (0.22, 4.13) | - |
| ***SLCO1B1*** | chr12:21284136-21392180 | - | - | 0.22 (0.07, 0.65)* |

| Druggable gene | Genomic coordinates | LDL-C (OR, 95% CI) | HDL-C (OR, 95% CI) | Triglycerides (OR, 95% CI) |
|---|---|---|---|---|
| ***SMARCA4*** | chr19:11071598-11176071 | 2.22 (1.98, 2.49)*† | 0.01 (0.0, 0.02)* | - |
| ***SOST*** | chr17:41831099-41836156 | - | 0.93 (0.25, 3.55) | 1.03 (0.39, 2.69) |
| ***ST3GAL4*** | chr11:126225535-126310239 | 2.25 (1.16, 4.39)* | 0.03 (0.0, 0.17)* | - |
| ***STX1A*** | chr7:73113536-73134002 | - | - | 0.89 (0.62, 1.28) |
| ***TECTB*** | chr10:114043493-114064793 | 0.88 (0.3, 2.64) | 0.63 (0.27, 1.5) | 1.49 (0.9, 2.49) |
| ***TMED1*** | chr19:10943114-10946994 | 2.06 (1.5, 2.83)*† | - | - |
| ***TNF*** | chr6:31543344-31546113 | 2.03 (1.05, 3.93)* | - | 1.21 (0.78, 1.9) |
| ***TNKS*** | chr8:9413424-9639856 | - | - | 0.79 (0.54, 1.16) |
| ***TNNC1*** | chr3:52485118-52488086 | - | 0.57 (0.36, 0.93)* | 1.73 (0.56, 5.36) |
| ***TNNC2*** | chr20:44451853-44462384 | - | 1.21 (0.69, 2.12) | 0.81 (0.43, 1.52) |
| ***TNXB*** | chr6:32008931-32083111 | 2.15 (1.55, 2.97)* | 0.98 (0.21, 4.67) | 1.64 (0.95, 2.82) |
| ***TOP1*** | chr20:39657458-39753127 | 2.3 (0.15, 35.62) | - | 16.72 (4.19, 66.8)* |
| ***TSSK6*** | chr19:19623227-19626838 | 1.64 (1.14, 2.37)* | - | 1.17 (0.99, 1.39) |
| ***TUBB*** | chr6:30687978-30693203 | - | 7.56 (1.18, 48.38)* | 4.46 (2.13, 9.36)* |
| ***TYRO3*** | chr15:41849873-41871536 | - | 0.8 (0.49, 1.29) | - |
| ***UCN*** | chr2:27530268-27531313 | - | - | 1.35 (0.53, 3.44) |
| ***UGT1A1*** | chr2:234668894-234681945 | 1.33 (0.74, 2.39) | - | - |
| ***UGT1A10*** | chr2:234545100-234681951 | 1.95 (1.25, 3.05)* | - | - |
| ***UGT1A3*** | chr2:234637754-234681945 | 2.04 (1.27, 3.25)* | - | - |
| ***UGT1A4*** | chr2:234627424-234681945 | 2.03 (1.27, 3.23)* | - | - |
| ***UGT1A5*** | chr2:234621638-234681945 | 2.03 (1.27, 3.25)* | - | - |
| ***UGT1A6*** | chr2:234600253-234681946 | 1.91 (1.22, 3.0)* | - | - |
| ***UGT1A7*** | chr2:234590584-234681945 | 1.94 (1.22, 3.07)* | - | - |
| ***UGT1A8*** | chr2:234526291-234681956 | 1.95 (1.23, 3.08)* | - | - |
| ***UGT1A9*** | chr2:234580499-234681946 | 1.94 (1.24, 3.05)* | - | - |
| ***VEGFA*** | chr6:43737921-43754224 | - | 0.22 (0.15, 0.3)* | 4.16 (2.45, 7.08)*† |
| ***VIM*** | chr10:17270258-17279592 | 1.02 (0.34, 3.06) | 0.77 (0.21, 2.78) | - |
| ***VKORC1*** | chr16:31102163-31107301 | 1.62 (0.45, 5.86) | - | 1.32 (0.52, 3.35) |
| ***WNT9B*** | chr17:44910567-44964096 | - | 6.95 (2.1, 23.05)* | |

**Table S4. Univariable MR estimates of drug targets with lipid records in clinicaltrials.gov and/or the British National Formulary (BNF).** * indicates significance in the discovery analysis; † indicates significance in both original and validation study and concordant direction of effect. OR = CHD odds ratio per 1-SD increase in LDL-C/HDL-C or triglycerides; CI = confidence interval.

| Gene | LDL-C (OR, 95% CI) | HDL -C (OR, 95% CI) | Triglycerides (OR, 95% CI) | Clinical trial (Record type) | BNF (Record type) | Phase | Mechanism of action | Indication |
|---|---|---|---|---|---|---|---|---|
| ADRB1 | - | 1.67 (0.58, 4.8) | - | Outcome | Side effect | 4 | AGONIST | Pain, Asthma, Nasal Obstruction, Glaucoma, Obstructive Lung Diseases, Hemorrhage, Cardiovascular Diseases, Serum Sickness, Bronchial Spasm, Rhinitis, Seasonal Allergic, Urticaria, Heart Arrest, Angioedema, Sinusitis, Sepsis, Hypotension, Orthostatic |
| ADRB1 | - | 1.67 (0.58, 4.8) | - | Outcome | Side effect | 4 | ANTAGONIST | Angina Pectoris, Hypertension, Myocardial Infarction, Cardiovascular Diseases, Arrhythmias, Cardiac, Migraine Disorders, Open-Angle Glaucoma, Ocular Hypertension, Glaucoma, Heart Failure, Left Ventricular Dysfunction |
| ADRB1 | - | 1.67 (0.58, 4.8) | - | Outcome | Side effect | 4 | PARTIAL AGONIST | Cardiovascular Diseases |
| ESR1 | - | 2.11 (1.13, 3.93)* | - | Outcome | Side effect | 4 | AGONIST | Neoplasms, Hypogonadism, Menorrhagia, Primary Ovarian Insufficiency, Acne Vulgaris, Osteoporosis, Postmenopausal |
| ESR1 | - | 2.11 (1.13, 3.93)* | - | Outcome | Side effect | 4 | ANTAGONIST | Breast Neoplasms, Neoplasms |
| ESR1 | - | 2.11 (1.13, 3.93)* | - | Outcome | Side effect | 4 | MODULATOR | Infertility, Dyspareunia, Breast Neoplasms, Osteoporosis, Postmenopausal |
| TNF | 2.03 (1.05, 3.93)* | - | 1.21 (0.78, 1.9) | Outcome | Side effect | 4 | INHIBITOR | Ankylosing Spondylitis, Crohn Disease, Psoriasis, Rheumatoid Arthritis, Colitis, Ulcerative, Psoriatic Arthritis, Immune System Diseases, Juvenile Arthritis |
| FRK | 0.76 (0.48, 1.21) | 0.57 (0.17, 1.94) | - | Outcome | Side effect | 4 | INHIBITOR | Neoplasms, Precursor Cell Lymphoblastic Leukemia-Lymphoma |
| BLK | - | - | 0.46 (0.31, 0.7)* | Outcome | Side effect | 4 | INHIBITOR | Precursor Cell Lymphoblastic Leukemia-Lymphoma, Neoplasms |
| DHODH | 0.66 (0.44, 1.0) | - | 7.42 (2.32, 23.71)* | Adverse event | Side effect | 4 | INHIBITOR | Rheumatoid Arthritis, Immune System Diseases, Multiple Sclerosis |

| Gene | LDL-C (OR, 95% CI) | HDL -C (OR, 95% CI) | Triglycerides (OR, 95% CI) | Clinical trial (Record type) | BNF (Record type) | Phase | Mechanism of action | Indication |
|---|---|---|---|---|---|---|---|---|
| HMGCR | 1.22 (1.03, 1.45)* | - | - | Outcome | Indication | 4 | INHIBITOR | Cardiovascular Diseases, Hypercholesterolemia, Dyslipidemias, Hyperlipidemias, Coronary Artery Disease, Hyperlipoproteinemia Type II, Myocardial Infarction, Heart Failure, Hypertension, Stroke, Stable Angina, Angina Pectoris, Type 2 Diabetes Mellitus |
| NPC1L1 | 2.01 (1.48, 2.73)*† | - | 2.56 (0.75, 8.68) | Outcome | Indication | 4 | INHIBITOR | Hypercholesterolemia, Hyperlipidemias, Cardiovascular Diseases |
| PPARG | 1.67 (1.04, 2.68)* | 0.71 (0.35, 1.48) | 2.18 (1.14, 4.15)* | Outcome | Indication | 4 | AGONIST | Type 2 Diabetes Mellitus, Diabetes Mellitus, Colitis, Ulcerative, Cardiovascular Diseases |
| PPARA | 3.77 (1.44, 9.85)* | - | - | Outcome | Indication | 4 | AGONIST | Cardiovascular Diseases, Hypercholesterolemia, Dyslipidemias |
| PCSK9 | 1.6 (1.45, 1.77)*† | - | - | Outcome | Indication | 4 | INHIBITOR | Hyperlipoproteinemia Type II, Coronary Artery Disease, Cardiovascular Diseases |
| INSR | - | 0.69 (0.47, 1.04) | 1.15 (0.83, 1.62) | Outcome | - | 4 | AGONIST | Diabetes Mellitus, Type 2 Diabetes Mellitus, Type 1 Diabetes Mellitus |
| NDUFS3 | - | 1.18 (0.83, 1.69) | - | Outcome | - | 4 | INHIBITOR | Diabetes Mellitus, Type 2 Diabetes Mellitus |
| NDUFA7 | - | 0.57 (0.3, 1.07) | - | Outcome | - | 4 | INHIBITOR | Diabetes Mellitus, Type 2 Diabetes Mellitus |
| NDUFA13 | 1.63 (1.13, 2.35)* | - | 1.18 (1.0, 1.39)*† | Outcome | - | 4 | INHIBITOR | Diabetes Mellitus, Type 2 Diabetes Mellitus |
| ALDH2 | 0.14 (0.07, 0.29)* | - | - | Outcome | - | 4 | INHIBITOR | Ectoparasitic Infestations, Alcoholism |
| NISCH | - | 0.57 (0.35, 0.93)* | 1.16 (0.31, 4.34) | Outcome | - | 4 | AGONIST | Hypertension |
| ABCA1 | 2.05 (1.34, 3.15)* | 1.41 (0.66, 3.0) | 2.4 (1.29, 4.49)* | Outcome | - | 4 | INHIBITOR | Cardiovascular Diseases |
| PDE3A | - | 0.79 (0.4, 1.58) | - | Outcome | - | 4 | INHIBITOR | Thrombosis, Obstructive Lung Diseases, Essential Thrombocythemia, Asthma, Cardiovascular Diseases, Coronary Artery Disease, Stroke |
| F2 | 0.17 (0.05, 0.59)* | 0.57 (0.13, 2.43) | 0.35 (0.13, 0.94)* | Outcome | - | 4 | INHIBITOR | Venous Thrombosis, Thrombosis, Unstable Angina, Thrombocytopenia, Atrial Fibrillation, Embolism, Stroke |
| TUBB | - | 7.56 (1.18, 48.38)* | 4.46 (2.13, 9.36)* | Adverse event | - | 4 | INHIBITOR | Breast Neoplasms, Neoplasms, Hodgkin Disease, Large-Cell Anaplastic Lymphoma, Non-Small-Cell Lung Carcinoma, Gout, Familial Mediterranean Fever |

| Gene | LDL-C (OR, 95% CI) | HDL -C (OR, 95% CI) | Triglycerides (OR, 95% CI) | Clinical trial (Record type) | BNF (Record type) | Phase | Mechanism of action | Indication |
|---|---|---|---|---|---|---|---|---|
| VEGFA | - | 0.22 (0.15, 0.3)* | 4.16 (2.45, 7.08)*† | Adverse event | - | 4 | ANTAGONIST | Retinal Neovascularization |
| VEGFA | - | 0.22 (0.15, 0.3)* | 4.16 (2.45, 7.08)*† | Adverse event | - | 4 | INHIBITOR | Diabetic Retinopathy, Retinal Neovascularization, Wet Macular Degeneration, Macular Edema, Colorectal Neoplasms, Neoplasms, Glioblastoma, Renal Cell Carcinoma, Non-Small-Cell Lung Carcinoma, Uterine Cervical Neoplasms |
| ERBB2 | - | 2.82 (0.25, 31.53) | - | Adverse event | - | 4 | INHIBITOR | Breast Neoplasms, Neoplasms, Non-Small-Cell Lung Carcinoma, Thyroid Neoplasms |
| RAF1 | 2.06 (1.48, 2.86)* | - | 2.63 (0.79, 8.83) | Adverse event | - | 4 | INHIBITOR | Neoplasms |
| PSMC3 | - | 1.12 (0.9, 1.4) | 0.68 (0.18, 2.53) | Adverse event | - | 4 | INHIBITOR | Multiple Myeloma, Neoplasms, Mantle-Cell Lymphoma |
| PSMA1 | - | 1.43 (0.5, 4.11) | - | Adverse event | - | 4 | INHIBITOR | Multiple Myeloma, Neoplasms, Mantle-Cell Lymphoma |
| PSMB8 | 2.23 (0.99, 4.98) | 1.06 (0.31, 3.59) | 1.09 (0.6, 1.97) | Adverse event | - | 4 | INHIBITOR | Multiple Myeloma, Neoplasms, Mantle-Cell Lymphoma |
| PSMB9 | 2.23 (0.99, 4.98) | 1.06 (0.31, 3.59) | 1.1 (0.6, 1.99) | Adverse event | - | 4 | INHIBITOR | Multiple Myeloma, Neoplasms, Mantle-Cell Lymphoma |
| PSMA5 | 2.47 (1.8, 3.39)*† | 0.08 (0.02, 0.29)* | - | Adverse event | - | 4 | INHIBITOR | Multiple Myeloma, Neoplasms, Mantle-Cell Lymphoma |
| PSMB10 | - | 1.11 (0.91, 1.35) | - | Adverse event | - | 4 | INHIBITOR | Multiple Myeloma, Neoplasms, Mantle-Cell Lymphoma |
| ALOX5 | - | 1.74 (1.18, 2.58)* | - | Adverse event | - | 4 | INHIBITOR | Asthma, Ulcerative Colitis, Rheumatoid Arthritis, Juvenile Arthritis |
| CACNB1 | - | 0.38 (0.2, 0.72)* | - | Adverse event | - | 4 | BLOCKER | Cardiovascular Diseases |
| CACNB1 | - | 0.38 (0.2, 0.72)* | - | Adverse event | - | 4 | MODULATOR | Fibromyalgia, Seizures, Epilepsy, Neuralgia, Restless Legs Syndrome, Postherpetic Neuralgia |
| PLG | 18.35 (5.47, 61.6)* | 5.48 (0.07, 456.86) | 0.75 (0.18, 3.14) | Adverse event | - | 4 | ACTIVATOR | Thrombosis, Pulmonary Embolism, Stroke, Myocardial Infarction, Heart Failure, Hepatic Veno-Occlusive Disease |
| PLG | 18.35 (5.47, 61.6)* | 5.48 (0.07, 456.86) | 0.75 (0.18, 3.14) | Adverse event | - | 4 | INHIBITOR | Hemorrhage, Menorrhagia |

| Gene | LDL-C (OR, 95% CI) | HDL -C (OR, 95% CI) | Triglycerides (OR, 95% CI) | Clinical trial (Record type) | BNF (Record type) | Phase | Mechanism of action | Indication |
|---|---|---|---|---|---|---|---|---|
| ITGB3 | 1.64 (1.06, 2.52)* | 2.79 (0.81, 9.62) | - | Adverse event | - | 4 | INHIBITOR | Thrombosis, Unstable Angina |
| MET | - | 0.66 (0.43, 1.01) | 1.09 (0.42, 2.87) | Adverse event | - | 4 | INHIBITOR | Thyroid Neoplasms, Non-Small-Cell Lung Carcinoma, Neoplasms |
| GSK3B | - | 0.45 (0.16, 1.25) | - | Adverse event | - | 4 | INHIBITOR | Bipolar Disorder, Psychotic Disorders |
| FDFT1 | - | - | 0.88 (0.44, 1.73) | Outcome | - | 3 | INHIBITOR | Hypercholesterolemia, Lipid Metabolism Disorders, Type 2 Diabetes Mellitus |
| CETP | 1.49 (1.29, 1.72)* | 0.91 (0.87, 0.95)*† | 1.98 (1.63, 2.4)*† | Outcome | - | 3 | INHIBITOR | Hypercholesterolemia, Lipid Metabolism Disorders, Hyperlipoproteinemia Type II, Coronary Disease, Cardiovascular Diseases, Acute Coronary Syndrome, Hyperlipidemias |
| ANGPTL3 | 1.21 (1.11, 1.33)* | 1.61 (0.52, 5.01) | 1.16 (1.08, 1.25)* | Outcome | - | 3 | INHIBITOR | Hyperlipoproteinemia Type II |
| AKT1 | - | 0.49 (0.18, 1.36) | - | Adverse event | - | 3 | INHIBITOR | Prostatic Neoplasms |
| SOST | - | 0.93 (0.25, 3.55) | 1.03 (0.39, 2.69) | Adverse event | - | 3 | INHIBITOR | Osteoporosis, Postmenopausal, Osteoporosis, Bone Diseases |
| CYP26A1 | 7.25 (4.25, 12.37)* | 0.22 (0.09, 0.51)* | 4.35 (2.79, 6.79)* | Adverse event | - | 2 | INHIBITOR | Psoriasis, Acne Vulgaris |
| LTA | 2.03 (1.04, 3.97)* | - | 1.22 (0.78, 1.9) | Adverse event | - | 2 | INHIBITOR | Rheumatoid Arthritis, Sjogren's Syndrome |
| LTB | 2.01 (1.03, 3.93)* | 1.2 (0.44, 3.25) | 1.11 (0.76, 1.62) | Adverse event | - | 2 | INHIBITOR | Sjogren's Syndrome, Rheumatoid Arthritis |
| NR1H3 | - | 1.07 (0.97, 1.18) | 0.86 (0.58, 1.27) | Outcome | - | 1 | AGONIST | Hypercholesterolemia |
| NR1H3 | - | 1.07 (0.97, 1.18) | 0.86 (0.58, 1.27) | Outcome | - | 1 | MODULATOR | Hypercholesterolemia |
| TOP1 | 2.3 (0.15, 35.62) | - | 16.72 (4.19, 66.8)* | Adverse event | - | 4 | INHIBITOR | Neoplasms |

**Table S5. Multivariable drug target MR estimates.** OR = CHD odds ratio per 1-SD increase in LDL-C/HDL-C or triglycerides; CI = confidence interval.

| Drug target gene | No. variants | Heterogeneity P-value | LDL-C (OR, 95% CI) | HDL-C (OR, 95% CI) | Triglycerides (OR, 95% CI) |
|---|---|---|---|---|---|
| *MARK4* | 6 | 7.47e-01 | 1.08 (0.82, 1.42) | 1.01 (0.33, 3.12) | 0.82 (0.18, 3.65) |
| *GIPR* | 4 | 3.61e-01 | 2.49 (1.2, 5.19)* | 0.45 (0.1, 2.07) | 1.91 (0.32, 11.34) |
| *NPC1L1* | 6 | 3.40e-01 | 1.5 (0.76, 2.96) | 0.76 (0.11, 5.15) | 1.11 (0.21, 5.92) |
| *NR1H3* | 8 | 8.01e-01 | 1.4 (0.45, 4.34) | 0.88 (0.73, 1.05) | 0.51 (0.37, 0.7)* |
| *SLC18A1* | 7 | 2.32e-01 | 5.31 (0.45, 62.01) | 0.21 (0.01, 3.79) | 0.37 (0.02, 9.01) |
| *LPAR2* | 5 | 2.93e-01 | 0.07 (0.0, 25.72) | 4.89 (0.02, 957.6) | 16.68 (0.07, 4147.03) |
| *CTSA* | 5 | 7.66e-01 | 0.09 (0.02, 0.48)* | 77.68 (6.48, 931.29)* | 117.3 (7.26, 1896.01)* |
| *SLC12A3* | 19 | 4.16-01 | 4.02 (2.34, 6.93)* | 1.16 (0.96, 1.4) | 0.88 (0.34, 2.25) |
| *PVR* | 10 | 2.15e-01 | 1.41 (0.93, 2.14) | 1.71 (0.54, 5.41) | 1.18 (0.3, 4.68) |
| *SCARB1* | 10 | 2.48e-03 | 25.58 (1.06, 614.7)* | 0.79 (0.47, 1.33) | 0.06 (0.0, 1.2) |
| *FDFT1* | 5 | 9.72e-01 | 1.87 (0.3, 11.58) | 0.94 (0.1, 8.91) | 0.93 (0.57, 1.5) |
| *APOB* | 16 | 5.48e-04 | 1.54 (1.02, 2.33)* | 2.48 (0.66, 9.29) | 2.37 (0.78, 7.19) |
| *GCKR* | 10 | 3.45e-01 | 3.11 (0.8, 12.13) | 1.79 (0.24, 13.49) | 0.84 (0.58, 1.24) |
| *CAD* | 11 | 7.56e-01 | 4.64 (0.9, 24.08) | 1.26 (0.26, 6.07) | 0.8 (0.66, 0.97)* |
| *CETP* | 36 | 3.00e-02 | 1.89 (0.83, 4.3) | 1.0 (0.84, 1.19) | 0.96 (0.46, 2.0) |
| *PLTP* | 5 | 6.02e-01 | 0.18 (0.04, 0.8)* | 24.18 (4.51, 129.56)* | 28.65 (4.7, 174.74)* |
| *MMP9* | 5 | 1.01e-01 | 0.01 (0.0, 0.06)* | 5.65 (0.72, 44.19) | 17.38 (1.99, 152.02)* |
| *LIPG* | 20 | 8.64e-02 | 0.24 (0.13, 0.47)* | 0.9 (0.74, 1.09) | 5.09 (3.36, 7.7)* |
| *DHODH* | 11 | 8.39e-01 | 2.18 (1.61, 2.95)* | 2.08 (0.94, 4.59) | 0.44 (0.16, 1.22) |
| *LACTB* | 6 | 7.157e-01 | 0.21 (0.0, 14.0) | 1.35 (0.11, 16.87) | 1.61 (0.06, 43.74) |
| *LILRB5* | 6 | 8.18e-01 | 1.63 (0.13, 20.4) | 0.82 (0.5, 1.36) | 1.1 (0.12, 10.51) |
| *STX1A* | 5 | 7.75e-01 | 0.51 (0.13, 2.04) | 0.34 (0.03, 3.29) | 0.7 (0.26, 1.87) |
| *HGFAC* | 7 | 9.98e-01 | 1.19 (0.13, 10.74) | 0.15 (0.01, 2.31) | 0.9 (0.07, 11.59) |
| *DCPS* | 9 | 2.46e-01 | 0.72 (0.41, 1.23) | 0.33 (0.16, 0.69)* | 6.7 (1.54, 29.28)* |
| *ST3GAL4* | 10 | 5.14e-03 | 1.46 (0.32, 6.8) | 1.03 (0.15, 6.97) | 6.72 (0.29, 156.11) |
| *APOA5* | 14 | 6.87e-01 | 10.23 (6.21, 16.84)* | 1.23 (0.93, 1.62) | 0.73 (0.61, 0.88)* |
| *APOA4* | 15 | 5.17e-01 | 1.53 (0.74, 3.14) | 1.0 (0.74, 1.37) | 1.11 (0.82, 1.5) |
| *APOC3* | 18 | 1.86e-03 | 2.24 (1.18, 4.27)* | 0.75 (0.62, 0.91)* | 0.81 (0.69, 0.95)* |
| *SLC22A2* | 16 | 2.07e-04 | 2.73 (1.66, 4.47)* | 1.26 (0.4, 3.93) | 1.38 (0.34, 5.63) |
| *VEGFA* | 4 | 4.39e-01 | 0.27 (0.04, 1.67) | 0.39 (0.0, 346.13) | 2.39 (0.01, 1013.57) |
| *HMGCR* | 7 | 4.30e-01 | 1.79 (1.28, 2.5)* | 0.13 (0.01, 1.38) | 0.31 (0.02, 4.26) |
| *NRBP1* | 10 | 8.43e-01 | 0.68 (0.06, 7.53) | 1.39 (0.09, 21.94) | 0.91 (0.47, 1.76) |
| *AMPD2* | 4 | 4.22e-01 | 2.43 (1.01, 5.81)* | 0.05 (0.0, 15.18) | 0.01 (0.0, 70.8) |
| *APOA1* | 17 | 1.76e-02 | 2.21 (1.26, 3.87)* | 0.84 (0.67, 1.04) | 0.97 (0.74, 1.29) |
| *PLG* | 5 | 5.49e-01 | 21.26 (12.83, 35.23)* | 0.17 (0.04, 0.7)* | 0.15 (0.03, 0.86)* |
| *SLC12A5* | 4 | 9.71e-01 | 0.0 (0.0, 0.16)* | 15.94 (0.81, 312.7) | 73.81 (1.12, 4881.0)* |
| *PEPD* | 6 | 8.91e-01 | 2.24 (0.66, 7.57) | 0.48 (0.21, 1.09) | 1.74 (0.72, 4.22) |
| *ATG4C* | 8 | 6.51e-01 | 0.42 (0.12, 1.47) | 4.32 (1.24, 15.04)* | 0.96 (0.41, 2.25) |
| *SMARCA4* | 15 | 8.61e-02 | 1.95 (1.8, 2.11)* | 0.97 (0.49, 1.92) | 0.13 (0.04, 0.44)* |
| *ALDH1A2* | 42 | 4.49e-01 | 1.29 (0.75, 2.22) | 1.1 (0.95, 1.28) | 1.63 (1.03, 2.57)* |
| *LDLR* | 18 | 6.01e-04 | 1.37 (1.15, 1.62)* | 0.23 (0.05, 1.11) | 0.12 (0.01, 1.03) |
| *PVRL2* | 15 | 3.29e-02 | 1.29 (1.08, 1.54)* | 1.12 (0.49, 2.53) | 1.03 (0.72, 1.46) |
| *APOE* | 14 | 6.96e-03 | 1.28 (1.16, 1.42)* | 0.9 (0.56, 1.46) | 0.87 (0.65, 1.17) |
| *APOC1* | 14 | 2.86e-03 | 1.3 (1.17, 1.46)* | 0.9 (0.52, 1.58) | 0.81 (0.6, 1.08) |
| *NCAN* | 6 | 4.55e-01 | 1.32 (0.2, 8.87) | 0.42 (0.06, 2.82) | 0.9 (0.14, 6.02) |
| *LILRB2* | 8 | 7.81e-01 | 0.87 (0.18, 4.19) | 1.02 (0.7, 1.49) | 0.91 (0.13, 6.51) |
| *PPARG* | 14 | 1.20e-02 | 2.77 (0.99, 7.76) | 0.38 (0.15, 0.96)* | 0.54 (0.15, 1.87) |
| *ANGPTL3* | 5 | 6.85e-01 | 7.52 (0.07, 826.16) | 0.43 (0.09, 1.97) | 0.35 (0.01, 10.44) |
| *AMPD3* | 5 | 7.32e-01 | 0.02 (0.0, 0.21)* | 1.52 (0.55, 4.22) | 5.21 (0.4, 67.63) |
| *ACP2* | 9 | 6.75e-01 | 0.77 (0.38, 1.54) | 0.84 (0.71, 0.98)* | 0.56 (0.43, 0.73)* |
| *DAGLA* | 9 | 4.84e-01 | 0.95 (0.67, 1.35) | 1.48 (0.51, 4.3) | 0.92 (0.43, 1.98) |
| *BLK* | 4 | 2.53e-01 | 0.04 (0.0, 0.61)* | 0.8 (0.01, 91.76) | 0.35 (0.15, 0.84)* |
| *CGREF1* | 5 | 5.99e-01 | 0.59 (0.1, 3.58) | 1.68 (0.11, 26.04) | 1.1 (0.87, 1.38) |
| *SLC5A6* | 11 | 7.23e-01 | 2.7 (0.54, 13.63) | 1.77 (0.36, 8.66) | 0.83 (0.69, 1.0) |
| *ATRAID* | 11 | 4.64e-01 | 2.49 (0.39, 15.91) | 1.62 (0.27, 9.53) | 0.86 (0.62, 1.18) |
| *CBLN3* | 4 | 9.61e-02 | 0.71 (0.27, 1.91) | 4.05 (0.08, 211.75) | 51.45 (0.28, 9606.89) |
| *PSMA5* | 4 | 7.79e-01 | 1.46 (0.66, 3.21) | 0.12 (0.0, 5.13) | 0.13 (0.0, 56.08) |
| *CELSR2* | 23 | 2.91e-02 | 1.88 (1.5, 2.34)* | 0.86 (0.25, 2.88) | 1.84 (0.52, 6.56) |
| *GALNT2* | 17 | 6.60e-03 | 0.98 (0.16, 6.02) | 0.61 (0.12, 3.13) | 0.94 (0.14, 6.14) |
| *GDF7* | 4 | 1.89e-01 | 0.87 (0.39, 1.98) | 1.16 (0.04, 37.71) | 2.44 (0.05, 127.94) |
| *KLHL8* | 8 | 9.95e-01 | 0.5 (0.07, 3.56) | 1.51 (0.28, 8.02) | 1.95 (1.08, 3.54)* |
| *RSPO3* | 5 | 5.57e-01 | 0.02 (0.0, 0.7)* | 11.04 (0.44, 279.24) | 100.3 (0.84, 11945.38) |
| *SLC22A3* | 11 | 9.02e-02 | 4.7 (3.44, 6.43)* | 2.64 (1.45, 4.81)* | 3.5 (1.96, 6.24)* |
| *RPL7A* | 6 | 5.17e-01 | 2.39 (1.44, 3.99)* | 5.41 (2.64, 11.1)* | 2.09 (0.09, 46.73) |
| *PTPRJ* | 4 | 8.41e-02 | 12.48 (1.22, 127.5)* | 0.51 (0.28, 0.92)* | 0.04 (0.0, 0.58)* |
| *SIDT2* | 10 | 1.41e-05 | 5.26 (0.2, 139.08) | 1.34 (0.52, 3.46) | 1.08 (0.34, 3.41) |
| *NAT2* | 9 | 6.67e-01 | 1.46 (0.2, 10.91) | 3.77 (1.16, 12.22)* | 1.39 (0.4, 4.86) |
| *GPR61* | 8 | 9.57e-01 | 2.05 (1.62, 2.6)* | 0.82 (0.32, 2.05) | 0.47 (0.18, 1.19) |

| Drug target gene | No. variants | Heterogeneity P-value | LDL-C (OR, 95% CI) | HDL-C (OR, 95% CI) | Triglycerides (OR, 95% CI) |
|---|---|---|---|---|---|
| ***RGS12*** | 8 | 9.93e-01 | 0.7 (0.2, 2.44) | 0.3 (0.07, 1.26) | 1.76 (0.56, 5.6) |
| ***CILP2*** | 4 | 7.39e-01 | 55.61 (0.0, 15957345.62) | 10.05 (0.02, 5027.9) | 0.04 (0.0, 3522.79) |
| ***SIK3*** | 17 | 1.57e-01 | 3.68 (2.02, 6.69)* | 0.76 (0.65, 0.88)* | 0.69 (0.59, 0.8)* |
| ***PCSK7*** | 10 | 1.17e-03 | 22.24 (2.45, 201.73)* | 0.96 (0.46, 2.02) | 0.53 (0.25, 1.11) |
| ***PTPN13*** | 5 | 1.26e-01 | 2.62 (0.34, 20.08) | 3.67 (0.15, 91.55) | 2.38 (0.19, 30.33) |
| ***UCN*** | 8 | 7.82e-01 | 10.72 (1.76, 65.49)* | 0.36 (0.03, 4.02) | 0.57 (0.41, 0.78)* |
| ***CTSB*** | 4 | 9.44e-01 | 1.68 (0.06, 47.93) | 1.9 (0.01, 327.35) | 0.9 (0.36, 2.26) |
| ***ABCA1*** | 21 | 1.11e-02 | 2.09 (0.59, 7.36) | 0.83 (0.58, 1.19) | 3.33 (1.39, 7.96)* |
| ***LIPC*** | 26 | 4.95e-01 | 1.45 (0.76, 2.76) | 1.09 (0.93, 1.29) | 1.72 (1.05, 2.81)* |
| ***C2*** | 5 | 1.34e-01 | 0.05 (0.0, 22.82) | 1.16 (0.4, 3.36) | 0.41 (0.07, 2.33) |
| ***ANGPTL4*** | 5 | 8.41e-01 | 2.8 (0.63, 12.39) | 0.43 (0.05, 4.09) | 0.94 (0.01, 122.83) |
| ***TNXB*** | 5 | 7.22e-01 | 2.54 (1.45, 4.43)* | 0.53 (0.07, 3.85) | 1.05 (0.28, 3.92) |
| ***FEN1*** | 11 | 6.08e-01 | 0.94 (0.69, 1.28) | 1.81 (0.81, 4.06) | 1.06 (0.62, 1.81) |
| ***GSTM4*** | 4 | 6.94e-01 | 3.46 (2.01, 5.94)* | 0.28 (0.07, 1.09) | 0.14 (0.01, 3.47) |
| ***PCSK9*** | 20 | 5.21e-03 | 2.39 (1.45, 3.96)* | 1.01 (0.28, 3.6) | 0.78 (0.24, 2.49) |
| ***LILRA3*** | 9 | 5.22e-01 | 0.07 (0.01, 0.81)* | 1.01 (0.65, 1.56) | 0.87 (0.12, 6.31) |
| ***RPS9*** | 6 | 8.20e-01 | 1.67 (0.17, 16.86) | 0.83 (0.46, 1.47) | 1.24 (0.1, 14.84) |
| ***FPR1*** | 5 | 3.47e-01 | 0.65 (0.2, 2.12) | 1.6 (0.31, 8.36) | 0.66 (0.01, 32.0) |
| ***OBP2B*** | 9 | 9.50e-01 | 1.25 (0.87, 1.8) | 2.13 (0.75, 6.1) | 0.07 (0.01, 0.62)* |
| ***INSR*** | 6 | 7.00e-01 | 6.78 (0.6, 76.86) | 19.37 (0.94, 400.01) | 16.08 (1.15, 224.46)* |
| ***TNKS*** | 4 | 2.69e-01 | 1.16 (0.05, 24.78) | 0.25 (0.0, 14.36) | 0.44 (0.12, 1.56) |
| ***SLC22A1*** | 13 | 5.143e-05 | 2.73 (1.53, 4.9)* | 1.16 (0.16, 8.61) | 0.3 (0.04, 2.2) |
| ***LPL*** | 27 | 6.78e-03 | 0.17 (0.03, 1.07) | 0.28 (0.1, 0.78)* | 0.48 (0.17, 1.35) |
| ***TSSK6*** | 4 | 7.39e-01 | 55.37 (0.0, 16133127.76) | 10.04 (0.02, 5084.4) | 0.04 (0.0, 3586.37) |
| ***EMILIN3*** | 7 | 7.29e-02 | 1.28 (0.69, 2.36) | 0.4 (0.02, 7.21) | 4.03 (0.18, 90.18) |
| ***NDUFA13*** | 4 | 7.39e-01 | 55.95 (0.0, 16832999.76) | 10.07 (0.02, 5139.88) | 0.04 (0.0, 3654.32) |
| ***BACE1*** | 6 | 4.52e-02 | 5.17 (0.34, 79.25) | 1.7 (0.78, 3.73) | 0.43 (0.17, 1.07) |
| ***LILRA5*** | 9 | 7.70e-01 | 0.08 (0.01, 0.59)* | 0.92 (0.59, 1.44) | 0.58 (0.08, 4.38) |
| ***BCAM*** | 12 | 1.28e-01 | 1.23 (0.99, 1.53) | 1.42 (0.62, 3.28) | 0.68 (0.49, 0.95)* |
| ***FPR3*** | 5 | 3.56e-01 | 0.66 (0.2, 2.15) | 1.58 (0.31, 7.93) | 0.63 (0.01, 28.98) |
| ***HAPLN4*** | 4 | 6.89e-02 | 4.0 (0.2, 80.12) | 46.52 (0.79, 2724.55) | 0.48 (0.03, 8.07) |
| ***HLA-DRB1*** | 8 | 4.04e-02 | 1.01 (0.47, 2.17) | 1.0 (0.57, 1.75) | 1.3 (0.51, 3.3) |
| ***SFTA2*** | 4 | 8.37e-02 | 0.62 (0.24, 1.63) | 1.36 (0.35, 5.34) | 1.92 (0.25, 14.86) |
| ***IGF2R*** | 16 | 1.23e-04 | 3.75 (2.25, 6.25)* | 0.35 (0.07, 1.73) | 0.24 (0.07, 0.8)* |
| ***HSD17B11*** | 4 | 4.23e-01 | 0.4 (0.02, 6.55) | 0.61 (0.08, 4.8) | 1.57 (0.33, 7.53) |
| ***LPA*** | 9 | 1.47e-02 | 4.44 (2.43, 8.13)* | 1.01 (0.47, 2.21) | 1.03 (0.37, 2.91) |
| ***TOP1*** | 7 | 6.86e-01 | 1.32 (0.81, 2.14) | 0.47 (0.09, 2.45) | 4.61 (0.49, 43.81) |
| ***PSMB8*** | 7 | 8.47e-01 | 3.36 (1.76, 6.4)* | 0.85 (0.21, 3.41) | 0.73 (0.39, 1.38) |
| ***HLA-DRA*** | 6 | 3.49e-01 | 1.06 (0.57, 1.95) | 0.86 (0.39, 1.91) | 7.48 (3.04, 18.39)* |
| ***NOTCH4*** | 15 | 3.53e-02 | 1.1 (0.46, 2.67) | 2.03 (1.5, 2.75)* | 0.99 (0.54, 1.8) |
| ***AGER*** | 11 | 6.88e-03 | 2.73 (0.8, 9.29) | 3.13 (1.89, 5.19)* | 2.27 (0.71, 7.25) |
| ***EHMT2*** | 5 | 1.05=e-01 | 328.24 (7.85, 13716.69)* | 0.53 (0.2, 1.39) | 0.04 (0.0, 0.38)* |
| ***SLC44A4*** | 5 | 1.07e-01 | 322.41 (7.63, 13629.22)* | 0.53 (0.2, 1.4) | 0.04 (0.0, 0.39)* |
| ***NEU1*** | 4 | NA | 1.18 (0.0, 690.62) | 0.91 (0.31, 2.68) | 0.09 (0.01, 0.91)* |
| ***HSPA1B*** | 4 | NA | 615016.56 (120.95, 3127274364.99)* | 0.21 (0.01, 3.43) | 0.11 (0.01, 1.23) |
| ***HSPA1A*** | 4 | NA | 224722364.93 (797.77, 63301607264984.07)* | 0.0 (0.0, 0.09)* | 48.9 (1.62, 1472.63)* |
| ***C6orf25*** | 4 | NA | 0.0 (0.0, 1.05) | 194.1 (0.82, 45848.5) | 0.3 (0.07, 1.29) |
| ***ABHD16A*** | 4 | NA | 641.94 (13.43, 30693.48)* | 0.04 (0.0, 0.39)* | 2.54 (0.53, 12.2) |
| ***APOM*** | 4 | NA | 292.41 (8.42, 10153.82)* | 0.6 (0.19, 1.96) | 0.26 (0.05, 1.23) |
| ***NCR3*** | 4 | 4.22e-01 | 5.55 (0.71, 43.25) | 0.21 (0.04, 0.97)* | 0.63 (0.15, 2.66) |
| ***HLA-C*** | 15 | 4.35e-04 | 1.31 (0.77, 2.22) | 2.72 (1.09, 6.79)* | 0.93 (0.7, 1.24) |
| ***C6orf15*** | 11 | 8.77e-01 | 5.24 (2.57, 10.65)* | 0.66 (0.45, 0.96)* | 0.37 (0.21, 0.64)* |
| ***DDR1*** | 4 | 1.52e-01 | 0.74 (0.25, 2.2) | 7.09 (0.34, 146.43) | 0.67 (0.06, 8.05) |
| ***GSTM2*** | 4 | 7.11e-01 | 4.4 (0.99, 19.64) | 0.28 (0.07, 1.13) | 0.03 (0.0, 8312.8) |
| ***CEACAM16*** | 13 | 3.85e-01 | 1.34 (1.01, 1.79)* | 2.29 (0.72, 7.3) | 1.21 (0.69, 2.11) |
| ***C4B*** | 5 | 9.04e-01 | 2.29 (1.25, 4.18)* | 0.34 (0.06, 1.97) | 1.35 (0.77, 2.39) |
| ***APOC4-APOC2*** | 11 | 1.35e-02 | 1.37 (1.21, 1.56)* | 0.62 (0.39, 0.98)* | 0.58 (0.44, 0.77)* |
| ***LTA*** | 4 | 3.72e-01 | 7.6 (1.01, 57.24)* | 0.26 (0.06, 1.12) | 0.44 (0.11, 1.74) |
| ***LTB*** | 4 | 3.76e-01 | 7.61 (1.04, 55.69)* | 0.26 (0.06, 1.11) | 0.44 (0.11, 1.69) |
| ***CYP21A2*** | 4 | 6.71e-01 | 2.22 (1.09, 4.55)* | 0.32 (0.05, 2.06) | 1.47 (0.44, 4.93) |
| ***TNF*** | 4 | 3.73e-01 | 7.69 (1.03, 57.64)* | 0.26 (0.06, 1.11) | 0.44 (0.11, 1.71) |
| ***HLA-B*** | 10 | 2.98e-02 | 1.85 (1.2, 2.84)* | 1.36 (0.65, 2.84) | 1.1 (0.85, 1.42) |

| Drug target gene | No. variants | Heterogeneity P-value | LDL-C (OR, 95% CI) | HDL-C (OR, 95% CI) | Triglycerides (OR, 95% CI) |
|---|---|---|---|---|---|
| ***APOC2*** | 12 | 4.74e-14 | 1.14 (0.71, 1.82) | 0.48 (0.14, 1.67) | 0.66 (0.32, 1.36) |
| ***HLA-DQA2*** | 13 | 2.58e-01 | 1.06 (0.77, 1.44) | 3.59 (2.12, 6.09)* | 1.82 (1.22, 2.72)* |
| ***LILRA4*** | 6 | 8.07e-01 | 0.06 (0.0, 0.73)* | 0.83 (0.49, 1.41) | 0.53 (0.05, 5.16) |
| ***PSMB9*** | 7 | 8.47e-01 | 3.35 (1.75, 6.41)* | 0.85 (0.22, 3.38) | 0.73 (0.38, 1.39) |
| ***HLA-DOB*** | 8 | 9.52e-01 | 3.27 (1.97, 5.45)* | 1.02 (0.41, 2.49) | 0.75 (0.49, 1.15) |
| ***EGFL8*** | 6 | 5.22e-04 | 1.49 (0.02, 110.15) | 8.25 (0.57, 118.64) | 0.72 (0.02, 24.37) |
| ***CFB*** | 5 | 1.33e-01 | 0.05 (0.0, 20.51) | 1.14 (0.39, 3.31) | 0.4 (0.07, 2.29) |
| ***LILRA6*** | 5 | 5.37e-01 | 1.63 (0.07, 39.37) | 0.72 (0.19, 2.73) | 0.46 (0.0, 112.3) |
| ***HP*** | 11 | 8.77e-02 | 1.82 (1.15, 2.88)* | 4.88 (1.89, 12.6)* | 0.58 (0.13, 2.64) |
| ***ITGB3*** | 4 | 7.49e-01 | 2.65 (0.34, 20.42) | 0.64 (0.01, 61.4) | 1.59 (0.17, 15.05) |
| ***RPL17*** | 4 | 5.36e-01 | 0.86 (0.09, 8.06) | 1.0 (0.47, 2.12) | 8.34 (0.45, 155.7) |

**Table S6. Publicly available GWAS data used in the phenome-wide association analysis (PheWAS).**

| Trait | No. Events | Sample size | URL |
|---|---|---|---|
| **Rheumatoid arthritis** | 14361 | 57284 | https://grasp.nhlbi.nih.gov/downloads/ResultsOctober2016/Okada/ |
| **Juvenile arthritis** | 2816 | 15872 | http://ftp.ebi.ac.uk/pub/databases/gwas/summary_statistics/HinksA_23603761_GCST005528/ |
| **Ankylosing spondylitis** | 10619 | 15145 | https://ftp.ebi.ac.uk/pub/databases/gwas/summary_statistics/CortesA_23749187_GCST005529/ |
| **Ulcerative colitis** | 6968 | 27432 | http://ftp.ebi.ac.uk/pub/databases/gwas/summary_statistics/LiuJZ_26192919_GCST003045/ |
| **Psoriasis** | 10588 | 33394 | http://ftp.ebi.ac.uk/pub/databases/gwas/summary_statistics/TsoiLC_23143594_GCST005527/ |
| **Crohn disease** | 22575 | 69268 | http://ftp.ebi.ac.uk/pub/databases/gwas/summary_statistics/LiuJZ_26192919 |
| **Stroke** | 34217 | 440328 | http://www.megastroke.org/index.html |
| **Asthma** | 5135 | 30810 | https://www.thelancet.com/journals/lanres/article/PIIS2213-2600(18)30389-8/fulltext |
| **Multiple sclerosis** | 14498 | 38589 | https://www.ncbi.nlm.nih.gov/pmc/articles/PMC3832895/ |
| **Gout** | 13179 | 882413 | http://ftp.ebi.ac.uk/pub/databases/gwas/summary_statistics/TinA_31578528_GCST008970/ |
| **Ovarian neoplasms** | 16924 | 85426 | http://ftp.ebi.ac.uk/pub/databases/gwas/summary_statistics/PhelanCM_28346442_GCST004462/ |
| **Parkinson disease** | 15056 | 27693 | https://drive.google.com/drive/folders/10bGj6HfAXgl-JslpI9ZJIL_JIgZyktxn |
| **Alzheimer disease** | 71880 | 455258 | https://www.ncbi.nlm.nih.gov/pubmed/30617256 |
| **Type 2 diabetes mellitus** | 74124 | 898130 | https://www.nature.com/articles/s41588-018-0241-6 |
| **Myocardial infarction** | 40149 | 126310 | http://www.cardiogramplusc4d.org/data-downloads/ |
| **Heart failure** | 47309 | 977323 | https://www.nature.com/articles/s41467-019-13690-5 |
| **Atrial fibrillation** | 60620 | 1030836 | https://www.nature.com/articles/s41588-018-0171-3 |
| **Diabetic nephropathies** | 5908 | 10875 | http://ftp.ebi.ac.uk/pub/databases/gwas/summary_statistics/vanZuydamNR_29703844_GCST005881 |
| **Chronic kidney failure** | 64164 | 625219 | https://www.ncbi.nlm.nih.gov/pmc/articles/PMC6698888/ |
| **Schizophrenia** | 35476 | 46839 | http://www.med.unc.edu/pgc/files/resultfiles/ |
| **Narcolepsy** | 1886 | 12307 | http://ftp.ebi.ac.uk/pub/databases/gwas/summary_statistics/FaracoJ_23459209_GCST005522/ |
| **Atopic dermatitis** | 21399 | 95464 | http://ftp.ebi.ac.uk/pub/databases/gwas/summary_statistics/PaternosterL_26482879_GCST003184 |
| **Biliary liver cirrhosis** | 2764 | 13239 | http://ftp.ebi.ac.uk/pub/databases/gwas/summary_statistics/CordellHJ_26394269_GCST003129 |
